# Highly Multiplexed 3D Profiling of Cell States and Immune Niches in Human Tumours

**DOI:** 10.1101/2023.11.10.566670

**Authors:** Clarence Yapp, Ajit J. Nirmal, Felix Zhou, Alex Y.H. Wong, Juliann B. Tefft, Yi Daniel Lu, Zhiguo Shang, Zoltan Maliga, Paula Montero Llopis, George F. Murphy, Christine G. Lian, Gaudenz Danuser, Sandro Santagata, Peter K. Sorger

## Abstract

Diseases like cancer involve alterations in in cell proportions, states, and local interactions as well as complex changes in 3D tissue architecture. However, disease diagnosis and most multiplexed spatial profiling studies rely on inspecting thin (4-5 micron) tissue specimens. Here, we use confocal microscopy and cyclic immunofluorescence (3D CyCIF) to show that few if any cells are intact in these thin sections; this reduces the accuracy of cell phenotyping and interaction analysis. In contrast, high-plex 3D CyCIF imaging of intact cells in thick tissue sections enables accurate quantification of marker proteins and detailed analysis of intracellular structures and organelles. Precise imaging of cell membranes also makes it possible to detect juxtacrine signalling among interacting tumour and immune cells and reveals the formation of spatially-restricted cytokine niches. Thus, 3D CyCIF provides insights into cell states and morphologies in preserved human tissues at a level of detail previously limited to cultured cells.

## MAIN

Precise characterization of mammalian cell morphology using molecular markers and optical microscopy has enabled detailed analysis of processes such as organelle dynamics, cell migration, and intracellular trafficking^1^. Rigorous assessment of morphology also plays a central role in the conventional histopathological diagnosis of disease, as performed by pathologists using haematoxylin and eosin (H&E) stained tissue sections^2^. In research settings, rapid innovations in confocal, structured illumination, stochastic reconstruction, and deconvolution microscopy have enabled ever more precise 3D characterization of cultured cells and model organisms^3^. The development of highly multiplexed tissue imaging (“spatial proteomics”^4^) potentially extends such approaches by enabling the measurement (most commonly using labelled antibodies) of a large number of cellular features in a preserved tissue environment.

However, with a few noteworthy exceptions (e.g. IBEX)^5–7^, most contemporary spatial proteomics is performed at resolutions sufficient for segmenting cells from each other (commonly 0.6 to 2.0 µm laterally) but not for resolving fine morphological and intracellular details. Additionally, most new development focuses on increasing assay plex^8^ rather than resolution. An emphasis on assay plex is logical for spatial transcriptomics^9^, where the ability to determine cell types and states depends primarily on the number of genes measured. However, when imaging with antibodies, the distributions of antigens represent an invaluable source of additional information about cell types and states.^10,11^ A substantial opportunity therefore exists to marry high-resolution 3D microscopy with high-plex profiling to study intact tissue and tumours. This is particularly true for methods compatible with the formaldehyde fixed paraffin-embedded (FFPE) tissue specimens that are used for patient diagnosis^12^ and the analysis of murine models^13^.

An additional complication is that almost all immunohistochemistry (IHC) and histopathological analysis of H&E-stained specimens is performed on ∼5-micron thick FFPE tissue sections. This provides a balance between visualizing detailed morphology and maintaining tissue integrity^14^. Conversely, imaging thicker sections using conventional wide field (2D) microscopy complicates the interpretation of IHC data due to interference from out of focus light^15^. In principle, this can be addressed by collecting multiple optical planes from each specimen followed by 3D reconstruction. Optical sectioning requires high numerical aperture (NA) objectives with high lateral resolution, making 3D optical microscopy ideal for precise, high-resolution study of cell morphology and cell-cell contacts.

In this paper, we investigate how detailed profiling of cells in a native tissue environment can be extended to 3D based on widely available commercial instruments and a modified version of the public domain cyclic immunofluorescence (CyCIF)^7^ protocol. In CyCIF and similar methods, high-plex images are generated by repeated rounds of 4-6 plex antibody staining, imaging, fluorophore inactivation (or antibody stripping) and then incubation with another set of antibodies. Two possible approaches to 3D CyCIF emerged from our preliminary studies: (i) one based on multi-spectral laser-scanning confocal microscopy, which enables 4-8 channels of imaging per cycle in FFPE specimens up to ∼50 μm thick at lateral resolutions as low as ∼120 nm (in super-resolution mode)^16^ and (ii) a second approach based on light sheet fluorescence microscopy^17^ able to image specimens up to ∼1 mm thick at roughly micron resolution. The former is ideal for detailed analysis of cell morphology and is the focus of the current paper; the latter is better suited to studying tissue ultrastructure and is still in development.

At the outset of our work, it was not obvious that subcellular structures would be preserved in FPPE specimens that have been stored in archives for many years, However, consistent with previous low-plex super resolution imaging studies^18^, we found that it was possible to resolve multiple organelles in FFPE specimens, including mitochondria, peroxisomes, secretory granules, and microtubule organizing centres in a manner qualitatively similar to tissue culture cells. Visualizing plasma membrane proteins at high-resolution improved cell type determination, uncovered new features of the surface proteome, and revealed the locations of juxtacrine signalling complexes. By examining specimens cut at different thicknesses, we found that that nearly all cells (and most nuclei) are incomplete in standard 5 μm tissue sections. Because many proteins are polarized on the plasma membrane, measuring only a slice through a cell (as in 5 μm section) can result in inaccurate phenotyping and obscure many cell-cell interactions within the dense tissue microenvironment. We conclude that thick section 3D CyCIF reveals new features of the tissue microenvironment and that understanding cells and tissue in 3D is essential for correctly interpreting data from conventional 2D imaging.

## RESULTS

3D CyCIF^7^ was used to image FFPE specimens representing five tissue types and spanning normal, cancerous, and pre-cancerous tissues. Each specimen was subjected to 8-18 rounds of cyclic imaging using a Zeiss LSM980 laser scanning confocal microscope, resulting in 20 to 54-plex images with 140 x 140 x 280 nm voxels (200 to 500 voxels per cell). Extracellular matrix (collagen) was imaged with Second Harmonic Generation by Fluorescence Lifetime Imaging Microscopy (SHG). Datasets averaged ∼500 gigabyte per mm^2^ of tissue (z-projections of each tissue can be viewed in **Supplementary Figs. 1-11** or at full-resolution online via Minerva^19^, see **Supplementary Table 1** for links and metadata).

To study the impact of tissue thickness on imaging, 5 µm to 50 µm thick sections were cut from FFPE blocks, mounted on glass slides and subjected to dewaxing and antigen retrieval^20^. We found that some tissues – particularly those cut thicker than ∼35 µm – did not maintain their integrity across multiple imaging cycles. To enable multiplexed imaging of friable tissue specimens, we developed an alternative approach in which specimens were mounted on coverslips, held in place with an “adhesive” coating (Matrigel) or 3D printed micro-mesh, and then placed in 3D printed carriers that fit on a standard microscope stage (*see Methods*; **Extended Data Fig. 1a-c**). Volumetric reconstruction of confocal image stacks was performed using Imaris® (RRID:SCR_007370) software; visualization involved inspection of primary data slices and 3D surface renderings. A newly developed 3D segmentation algorithm was used to identify individual cells, generate UMAP and similar embeddings, and distinguish among major immune and tumour cell types (**Extended Data Fig. 1d**)^21^.

### Standard 5 µm histological sections contain few intact cells and nuclei

Dehydration is a component of the paraffin embedding process that is known to change the volume of FFPE specimens^22^. We found that sections cut at 5 µm on a microtome expanded to ∼9 µm following rehydration, antigen retrieval, and antibody labelling; similar proportional expansion was observed over a 5 to 35 µm range of section thicknesses (slope ∼1.5). Conversely, flash-frozen tissue shrank ∼1.5 fold upon dehydration but then expanded to its original thickness upon rehydrated during antigen retrieval (**Extended Data Figure 1e-f**). When we measured the aspect ratios of tumour cell nuclei, which were ellipsoidal in primary melanoma, we did not observe any systematic bias along the imaging (and re-hydration) axis **(Extended Data Figure 1g-h)**. We conclude that rehydrated sections are likely to be representative of native tissue in 3D, at least on a local scale, but that sections processed for H&E imaging are ∼1.5-fold thinner. Thicknesses reported in the remainder of this paper represent the hydrated thickness as measured during image acquisition; when necessary “cut@” is used to describe thickness at the time of sectioning in the FFPE state (as in “cut@5 µm).

Multiple layers of intact nuclei were visible in 30-40 μm tissue sections (**Fig. 1a)** but fewer than 5% of nuclei were intact in sections cut@5 μm thickness (**Extended Data Figure 1i-j**). To study the impact of this on interaction analysis and cell phenotyping, volumetric reconstruction was performed on communities of cells; **Figure 1b-c** shows a small community, comprising a dendritic cell (*D*) and two T cells (*T1, T2*), from a 54-plex CyCIF image of a 35 µm thick section of invasive (vertical growth phase; VGP) primary melanoma; the cells spanned ∼25 µm along the optical axis (Z; upper image) and a similar distance in the plane of the specimen (X,Y; lower image). Immune cell phenotypes were assigned based on patterns of expression of CD antigens and immune regulatory proteins; we used this approach on 3D data to create ground-truth phenotypes. To simulate thin section images, we then generated 2D maximum intensity projections from 9 µm virtual sections (**Fig. 1d**; equivalent to cut@5 μm sections and labelled I to V) and again performed cell type calling. Many discrepancies were observed between the 2D and 3D data. In virtual section III, for example, *T1 was* incorrectly scored as positive for PD1 due to overlap with cell *D* along the Z axis. In section I, true positive staining from *D* (CD11c and MX1; see **Supplementary Table 2** for protein nomenclature) was scored as background because the corresponding nuclei were largely absent from the section (segmentation methods rely on nuclei to locate cells^23^ and 12% of true cytoplasmic signals lacked a detectable nucleus in a 2D virtual section; **Fig. 1e**). The magnitude of the cell type assignment error varied based on marker distribution: polarized proteins (i.e., LAG3 and MX1) were most likely to result in false negative calls (30-40% of cells) whereas uniformly distributed proteins such as MART1 resulted in ∼5% false negative calls (**Fig 1e)**. Thus, imaging incomplete cells can result in inaccurate phenotyping, especially for markers with non-uniform distributions. Fragmentation of cell in cut@5 µm sections also impacted the number of cell-cell interactions identified by proximity analysis, which fell as section thickness decreased (**Extended Data Fig. 1k-m** and consistent with data from serial section reconstruction of skin)^24^.

**Figure 1:**
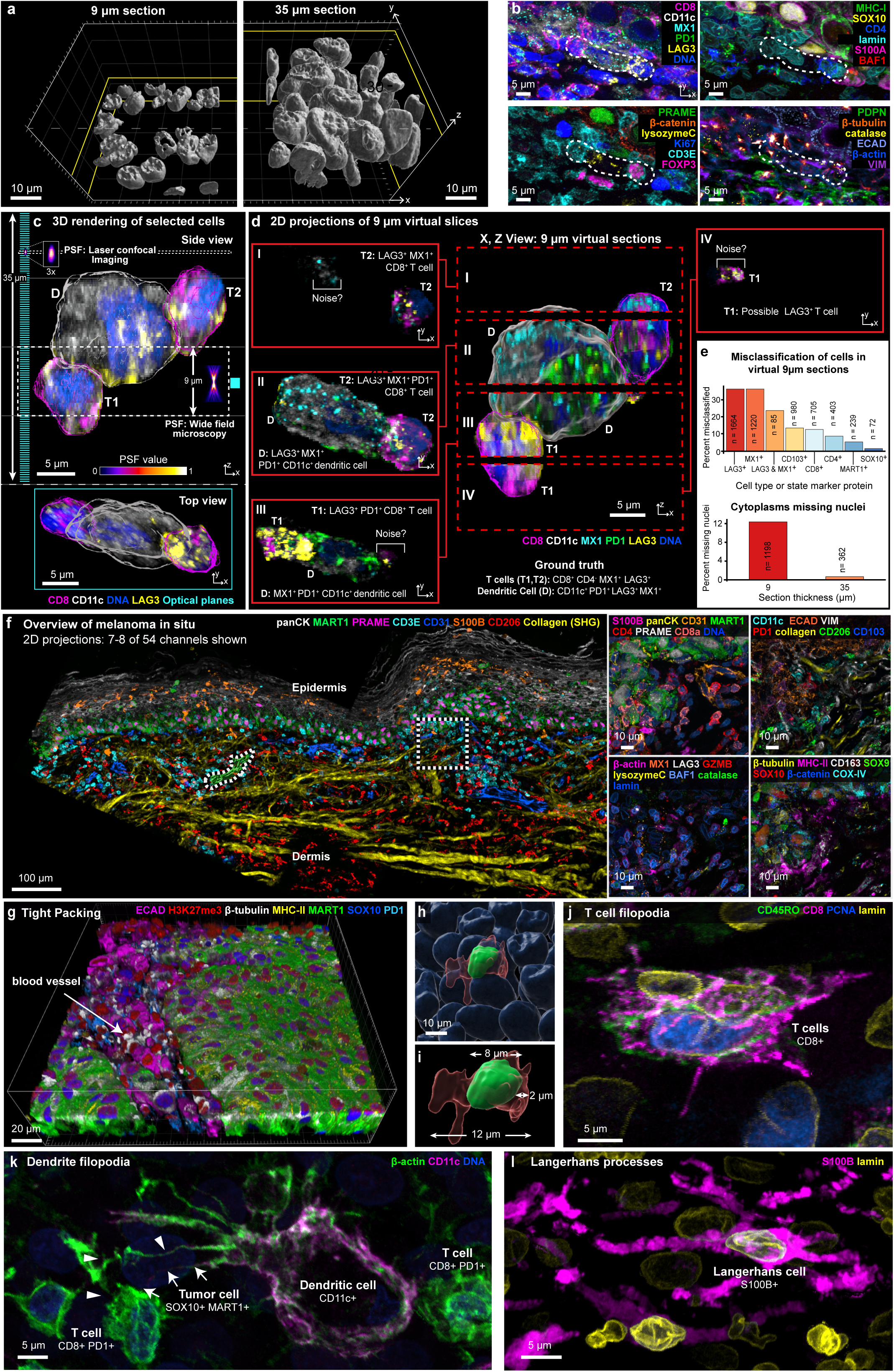
Demonstrating the need for thick tissue sections using 3D CyCIF. **a,** Surface rendering illustrates nuclear volumes in 9 µm (cut@5 µm, left) and 35 µm (cut@24 µm) tissue sections (right). Scale bars, 10=µm. **b,** Immunofluorescence images of 6-marker subsets illustrating the microenvironment of the cellular community from the VGP highlighted in **c-d** (dotted lines). **c,** 3D rendering of 3 selected cells from (**b**). Comparison of the point spread functions (PSF) and optical planes (cyan; 280 nm spacing) for laser scanning confocal and widefield microscopy performed with 40x/1.3NA objective. Upper: x,z (side) view. Lower: y-x (top) view. **d,** Computed (virtual) 9 µm sections generated from 3-dimensional data were used to generate x-z (centre) and x-y 2D projections (red boxes to left and right; labelled I-IV). **e**, Top: Percentage misclassified cells in a virtual 9 µm section when stained with polarized (LAG3, MX1) and diffuse (CD103, MART1) markers, compared to ground-truth data from 35 µm sections. Bottom: Quantification of the percent of cells missing nuclei from virtual 9-micron versus 35-micron tissue sections. A minimum size cut-off of 50 voxels was used to eliminate debris. **f,** Multi-modal image integrating 3D CyCIF with second harmonic generation (SHG) signal of collagen highlighting the melanoma in-situ region (MIS). Maximum intensity projection of selected channels at lower magnification (left), with additional marker subsets for the indicated ROI (right). Scale bars, 100=µm and 10=µm. **g-i,** FOVs capturing the boundary of a vertical growth phase tumour, highlighting densely packed cells at low-resolution (**g**) and in high-resolution renderings of an individual cell (**h-i**). **j-l,** Examples of cells with extended membrane processes in melanoma: **j**, Cluster of CD8 T cells in metastatic melanoma. **k,** Dendritic cell with filopodia extensions in metastatic melanoma. Two filopodia contacting a T cell and tumour cell, labelled with arrows or arrowheads, respectively. **l,** Langerhans cell in the MIS. Scale bars 5=µm.

The field of stereology is focused on identifying and mitigating biases arising from studying 3D objects using 2D images ^25,26^. Our findings suggest a way of extending stereology to high plex tissue imaging by generating simulated 2D datasets from ground truth 3D data (via virtual sections and downsampling the resolution) and then using this to develop solutions for 2D bias. We note, however, that the images in **Figure 1d** do not correctly represent what would be captured by 2D wide-field slide scanners, which use low NA objectives, resulting in low signal-to-noise ratios (**Extended Data Figure 1n)**. In contrast, confocal microscopes collect photons from a point source that is at least ∼3-fold smaller in X,Y and 5-fold smaller in Z than a slide scanner (assuming a 0.5 NA objective; ) and use a pinhole to reject out-of-focus light. This difference must be accounted for when comparing 2D and 3D high-plex images (e.g., using a mix of classic image processing methods and machine learning models).

### Tumour microarchitecture

Melanoma specimens in our dataset included a (i) a pre-invasive cutaneous melanoma in situ (MIS; **Fig. 1f**), (ii) an invasive vertical growth phase (VGP) primary melanoma from the same patient (**Extended Data Fig. 2a-b**), and (iii) a metastatic melanoma to the skin from a different patient. For simplicity, we focus our analysis on these specimens but data for other specimens is available in **Supplementary Figures 1-11**. We observed that cells in both tumours and the stroma were densely packed, except in areas where blood vessels or ECM filled the voids (**Fig. 1g; Supplementary Video 3**). In the MIS, for example, nuclei averaged 5.0 µm in diameter (mean 7.2 µm ± 2.3 µm SD for melanocytes and 4.9 µm ± 2.8 µm for immune cells). Cells averaged 13 µm ± 4.3 µm along the major axis, 6.1 µm ± 1.9 µm along the minor axis (consistent with recent and historical estimates)^27^. Thus, the depth of the cytoplasmic compartment, scored as the distance from the plasma membrane to the nuclear lamina, was often ∼1 to 6 µm and the membranes of neighbouring cells were ∼1 to 1.5 µm apart (**Fig. 1h,i)**.

Some cells had highly extended cell bodies and cytoplasmic processes. For example, in **Figure 1j**, three CD8^+^ T cells from metastatic melanoma exhibited multiple filipodia extending 5-10 µm from the cell body. A dendritic cell had 20-30 µm filopodia as well as membrane ruffles that contacted multiple CD8^+^ PD1^+^ effector T cells; these specialized filopodia enable a switch from antigen sampling to antigen presentation during T cell priming^28^ (**Fig. 1k, Extended Data Fig. 2c-d**). Langerhans cells, skin-resident macrophages that function similarly to dendritic cells^29^, also had many membrane extensions, with branches extending 30-50 µm across multiple Z planes (**Fig. 1l**). 3D imaging was essential to identify these morphologies and distinguish changes in cell shape from changes in orientation: 2D views of VGP melanoma suggested that round cells were more common in the tumour centre and elongated cells at the tumour margin. In 3D this was seen to be a difference in orientation: cells in the centre were more likely to be viewed end-on whereas those in the margin were rotated ∼90° (**Extended Data Fig. 2e-g).** Theoretical studies of cell migration have demonstrated a dependency on the geometry of cell packing^30^ and accurate 3D representations of tissues are likely to be useful in such studies. Tight cell packing, extended processes, and overlap along the Z-axis also explain why accurate single-cell segmentation of 2D images and spatial transcriptomic profiles^31^ is unlikely to be completely accurate even with optimized algorithms.

### Blood vessels and trans-endothelial migration

Thick section 3D imaging made it possible to dissect components of the tissue microarchitecture not generally visible in 2D, such as a 100 µm-long dermal blood vessel (venule) in the MIS (**Fig. 2a; Supplementary Video 2)**. In this vessel, ∼10 vimentin and beta-catenin-positive endothelial cells formed a tube enclosing erythrocytes and a neutrophil. At the distal end, a T_helper_ and dendritic cell were visible where the vessel appeared to branch. Most remarkable was a B cell with low sphericity (value of ∼0.35) flattened against the vessel wall, a morphology consistent with trans-endothelial migration (diapedesis) of immune cells from vessels into tissues^32^. These features were not evident in virtual 5 µm-thick sections (**Extended Data Fig. 3a**). Elsewhere in the dermis, another B cell had its nucleus traversing a vessel wall while its cell body remained inside the vessel, and in metastatic melanoma, a T cell was visible suspended within a vessel (**Extended Data Fig. 3b-c**). In the MIS, we found that B cells were the cell type most likely to associate with collagen fibres in the dermis^33,34^ (**Fig. 2b, c)** and were often (n = 11 of 14) stretched into irregular shapes. This was not a feature of all B cells; those found in the stroma and VGP were often round (**Extended Data Fig. 3d-e**). Functions have only recently been ascribed to B cells in the skin, and our images provide direct evidence of B cell recruitment from the vasculature into the dermis, followed by collagen binding, at densities consistent with other reports^35^.

**Figure 2:**
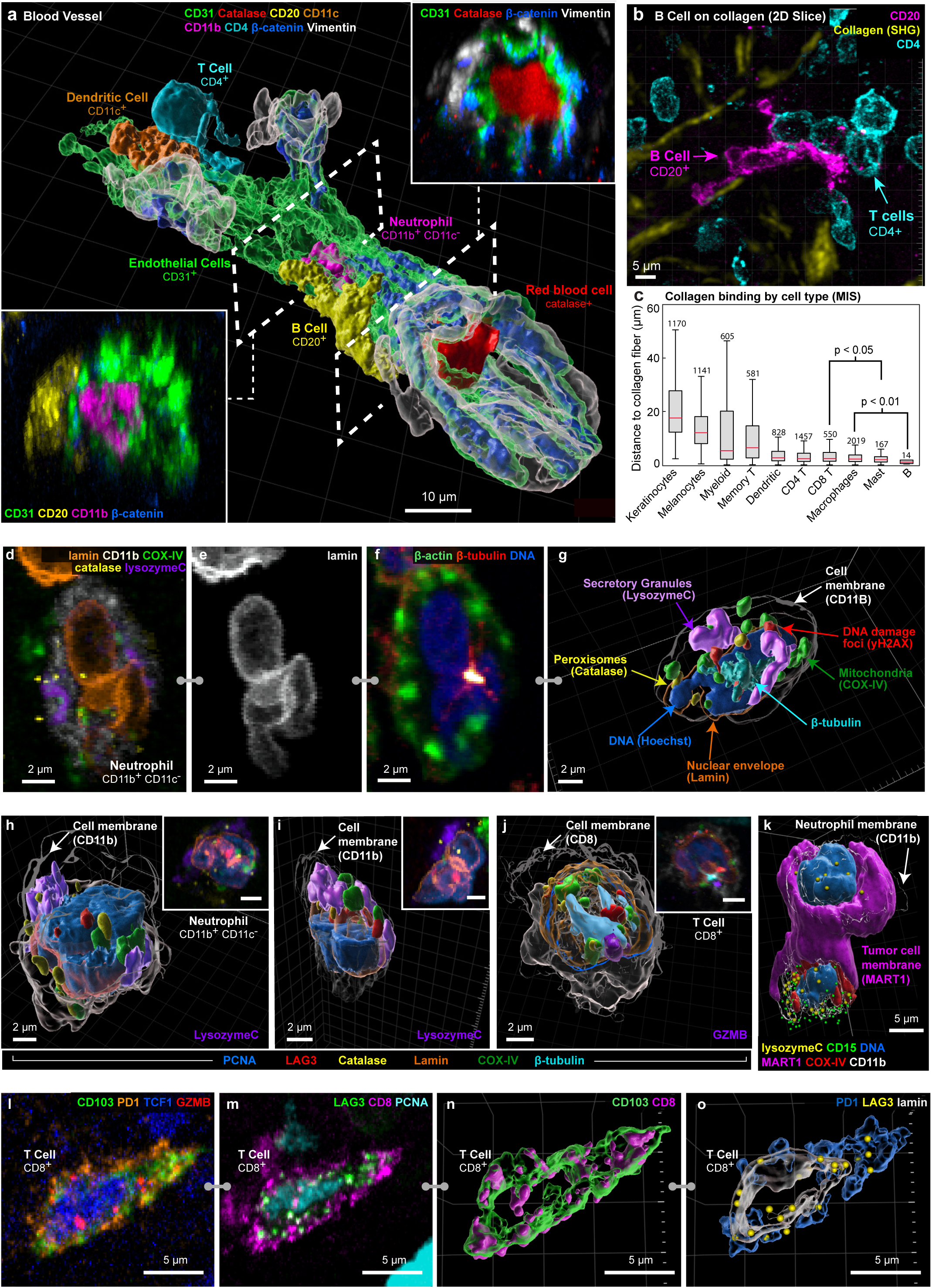
Visualizing tumour microarchitecture and complex organelle and cell-surface morphologies. **a,** Surface rendering of a segment of an intact blood vessel (delineated by CD31^+^ endothelial cells) within the MIS region. Dashed lines demarcate cutting planes for cross-sectional views (insets lower left and upper right) which reveal internal components of the blood vessel including: CD11c^+^ dendritic cell, a CD4^+^ T cell, CD11b^+^ CD11c^−^ neutrophil, catalase^+^ red blood cell and CD20^+^ B cell undergoing trans-endothelial migration. Scale bar, 10=µm. **b,** CD20^+^ B cell (magenta) in the MIS with elongated morphology interacting closely with collagen fibres (yellow), shown as a maximum intensity projection. Scale bars, 5=µm. **c,** Tukey box plot illustrating the distances between collagen fibres and different cell types in the MIS. Statistical significance was assessed using one sided unpaired student’s *t-test*. Centre line, median; box limits, upper and lower quartiles; whiskers, minimum and maximum after removing outliers. **d-i,** Selected channels of 54-plex 3D CyCIF images used for identifying cell types and organelles. All cells shown were from the dermis of the MIS (full marker assignment in **Supplementary Figure 13**)**. d-f,** Maximum projections of a neutrophil showing marker subsets for identifying specific organelles (**d**), a multi-lobed nucleus (**e**), the cytoskeleton (**f**). **g,** A 3D rendering of the cell shown in (**d-f**). **h-j,** 3D rendering and maximum intensity projection inset (upper right) for selected cells, including neutrophils in the MIS (**h** & **i**) and a T cell in metastatic melanoma (**j**). **k,** 3D rendering of a neutrophil (CD11b^+^CD11c^−^) interacting with a MART1^+^SOX10^+^ tumour cell (magenta) in the MIS. **l-o,** T cell (CD8^+^ CD103^+^ PD1^+^ TCF1^+^ LAG3^+^ GZMB^+^ PCNA^+^), showing distribution of intracellular (**l** & **n**) and membranous (**m** & **o**) markers as maximum projections (**l-m**) and surface rendering (**n-o**).

### Organelle and cell surface morphologies

High-plex imaging of whole immune and tumour cells in 3D revealed a wide variety of distinctive intracellular and plasma-membrane structures (**Fig. 2d),** including lineage-associated differences in nuclear lamina (e.g., a multi-lobed, hyper-segmented nucleus in neutrophils; **Fig. 2e**), microtubule organising centres (**Fig. 2f**), peroxisomes (based on catalase staining), secretory granules and/or ER (lysozyme C in neutrophils and granzyme B in T cells), DNA damage foci (γH2AX), mitochondria (COX IV), and biomolecular condensates (MX1^36^; **Fig. 2g-k, Extended Data Fig. 4a**). Some features were found in many cell types and others only in selected lineages (e.g., catalase foci in dendritic cells and γH2AX foci in keratinocytes and myeloid cells). These data suggest that cells in tissues can be characterized at a level of detail hitherto described only in cultured cells and some model organisms.

Proteins used for immune cell subtyping exhibited a wide range of intracellular distributions. Some proteins were found throughout the plasma membrane, for example the myeloid cell integrin CD11c, skin-homing T cell integrin CD103, and MHCII receptor (in tumour and antigen presenting cells). Other proteins were found in discontinuous islands (CD4 and CD8 in T cells) or puncta (the immune checkpoint protein LAG3; **Fig. 2l-o**). Some of these distributions have the potential to provide information on activity or cell state. For example, newly synthesized LAG3 localizes to endosomes but can rapidly translocate to the plasma membrane where it is activated by binding to MHC class II^37^ on the membranes of apposed cells. Across specimens, we found 1-20 LAG3 puncta per cell, both inside cells and at the plasma membrane (**Extended Data Fig. 4b**). Granzyme B (GZMB) staining was diffuse and globular in CD4 T cells and punctate in CD8 T cells, consistent with localization to cytoplasmic granules. GZMB mediates the cytotoxic activity of T and NK cells and globular GZMB can be used to identify activated memory CD4 T cells (**Extended Data Fig. 4c-d; Supplementary Video 4**)^38^.

### Functional tumour-immune interactions

Changes in the distributions of cell surface proteins also revealed functional interactions among cells. For example, the immune checkpoint receptor PD1 and its transmembrane ligand PD-L1 varied from a relatively uniform distribution in the membrane to punctate (**Fig. 3a-b; Extended Data Fig. 4e-g**), with the punctate morphology most evident when PD1^+^ T cells were in contact with PDL1 expressing cells (primarily dendritic cells)^39^. In some cases, many PD1 and PDL1 puncta were visible across an extended domain of membrane-membrane apposition (e.g., 13 foci over 40 µm^2^ in **Fig. 3b**), in an arrangement consistent with formation of multiple juxtacrine signalling complexes. Multiple distinctive membrane structures were often visible in a single cell; for example, a PD1^+^ CD8 T cell contacting a tumour cell with filopodia while also binding PDL1 from a neighbouring dendritic cell (**Fig. 3c-e**, **Extended Data Fig. 4h-j**). In a different multicellular community, filipodia from a CD8 T cell contacted a CD4 T helper cell (**Fig. 3f**; inset, **Extended Data Fig. 4k)**, which in turn contacted another CD8 T cell that was in contact with a tumour cell and dendritic cell. GZMB in the CD8 T cell was polarized toward the tumour cell **(Fig. 3g**; green arrowhead) even though PD1-PDL1 complexes had formed ∼1.5 μm away along the T cell membrane (blue arrowhead).

**Figure 3:**
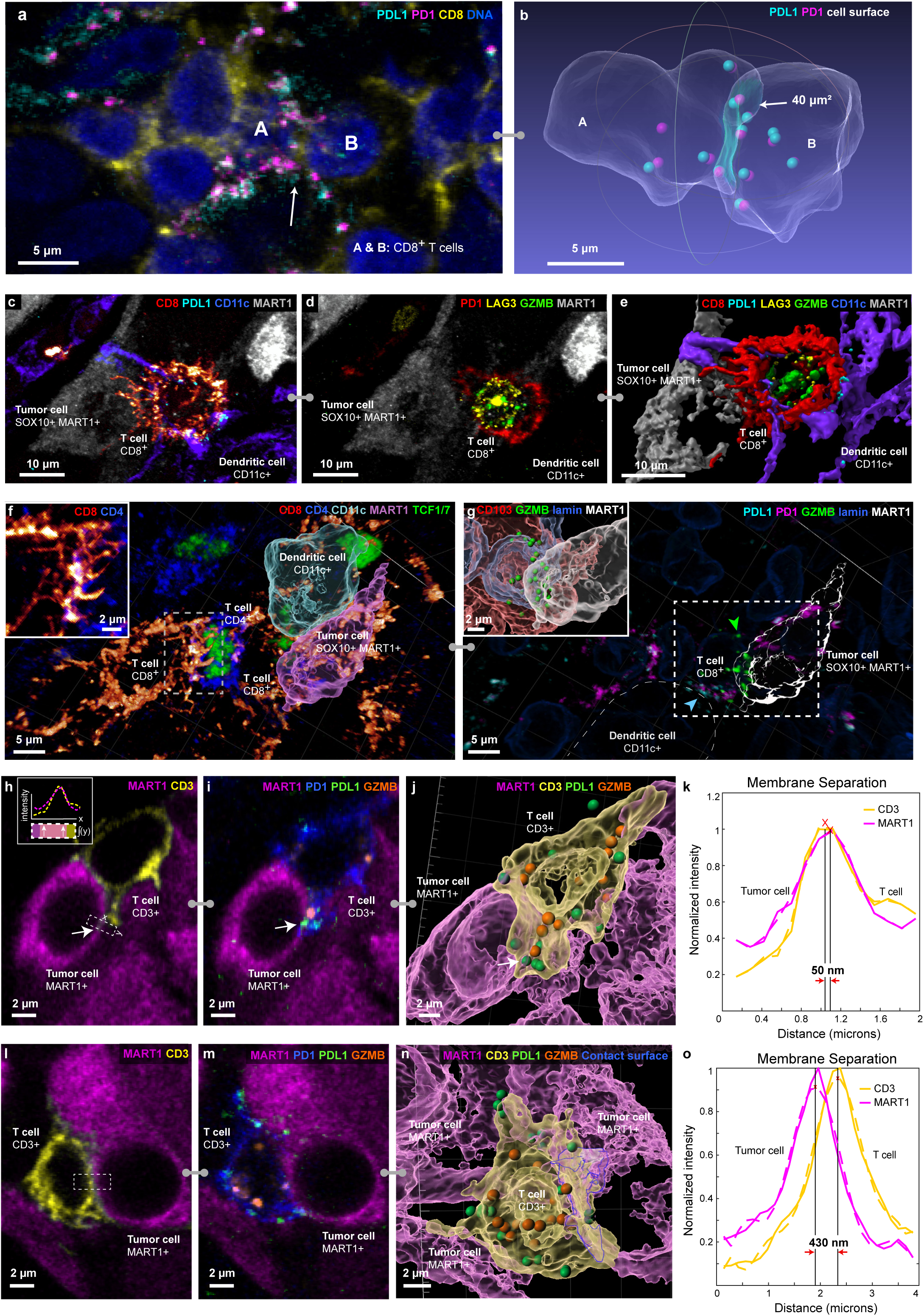
Visualizing functional tumour-immune interactions in native tissue. **a-b,** Two CD8^+^ T cells expressing PD1 and PDL1 interacting within the VGP. Shown as a maximum intensity projection (**a**) and transparent surface mesh showing contact area and colocalized PD1 and PDL1 as spheres (**b**). **c,** CD8^+^ T cell and dendritic cell interacting with a tumour cell with filopodia. **d,** Same cells as (**c**) with GZMB, PD1, and LAG3 shown; markers show that the T cell is activated and cytotoxic. **e,** surface rendering of interactions in (**c**) and (**d**), showing filopodia in greater detail. **f,** Multicellular interaction in metastatic melanoma. A dendritic cell interacting with a tumour cell (with surface rendering overlaid onto immunofluorescence) and a CD4 T_helper_ cell interacting with a CD8 T cell through filopodia (shown in inset and **Supplementary Video 6**). **g,** Same FOV in (**f**), showing PD1 and PDL1 colocalization between CD8 T cell and dendritic cell (blue arrow). In the CD8 cell, GZMB is polarized towards the tumour cell (green arrow and inset). Location of inset shown by box with dotted line. Scale bars as indicated. **h-j**, Interaction of a MART1^+^ tumour cell with CD3^+^ T cell, shown as single optical planes (**h-i**) and as a surface rendering (**j**). Inset in panel **h** depicts computation of a membrane intensity profile by integrating the fluorescence intensity parallel to the cell membrane (y) and plotting it as a function of distance perpendicular to the membrane (x). (**j**). Arrows indicate region of active PD1-PDL1 interaction. **k**, Membrane intensity profile of region indicated in (**h**), spanning a point of tumour-immune cell contact demonstrating a Type I interaction. **l-n**, Interaction of MART1^+^ tumour cell with CD3^+^ T cell, shown as single optical plane (**l-m**) and as a surface rendering (**n**) and highlighting PD1-PDL1 interactions and proximity to GZMB granules. **o**, Membrane intensity profile shown in (**l**) demonstrating a Type II interaction. For plots in **k** and **o**, solid lines are from raw data, dashed lines from polynomial curve fitting. Red ‘X’s mark the maximum intensity along the intensity profile which we defined as the midpoint of the cell membrane for each channel.

Regulation of TCR signalling is highly localized, so we looked for evidence of T cells experiencing simultaneous and potentially divergent regulatory or functional interactions with more than one neighbouring cell. These included CD8^+^ T cells with evidence of cytotoxicity (e.g., the presence GZMB granules) and cell membranes in close proximity to tumour cells (e.g., 50 nm separation – consistent with synapse formation; **Fig 3 h-j**), that also contained PD1-PDL1 complexes along the CD3-expressing membrane. The GZMB granules and PD1-PDL1 complexes were within a few microns of each other and the T cells containing them were also in contact (more distantly) with myeloid cells and potentially regulatory CD4 T cells. We found multiple examples of such communities, suggesting that a single T cell may be subjected to simultaneous negative and positive regulatory signals from interacting cells in a local niche (**Fig. 3h-o).**

### Tumour lineage plasticity

Mechanisms of melanoma initiation remain elusive^40^, although the genetics of later stage melanoma are well characterized^41^ and epigenetic changes (e.g. reductions in 5-Hydroxymethylcytosine (5hmc) levels)^42^ have been implicated. The presence of melanoma precursor fields^43^ and MIS is currently scored by changes in the morphologies, numbers, and positions of melanocytic cells in H&E and IHC images^44^. In the MIS region, most melanocytic cells were located at the dermal-epidermal junction (DEJ), interacted with keratinocytes, and retained a dendritic morphology (dendrites are involved transfer of UV-protective melanin; **Fig. 4a**). Also present were pagetoid melanocytic cells that lacked dendrites and had a rounded, ameboid morphology (cytological atypia), which was most obvious among cells that had migrated towards the top of the epidermis (**Fig. 4b-h)**. Pagetoid spread by single and small groups of cells is a hallmark of oncogenic transformation^45^. Nonetheless, the MIS and underlying dermis were not highly proliferative, with only 1% of cells (n = 110) positive for the Ki67^+^ proliferation marker. Among Ki67^+^ cells, 34% were T cells while the remainder consisted of monocytes (28%) and endothelial cells (2.7%); only a single melanocytic cell was Ki67^+^ (**Extended Data Fig. 5a**). By contrast, in the invasive VGP melanoma domain from the same specimen, 11% of all cells were Ki67^+^ with melanoma tumour cells the most proliferative (45% Ki67^+^), followed by monocytes (44%). Thus, the MIS had the hallmarks of early oncogenic transformation and migration, but limited cell division (proliferation is known to vary among MIS specimens)^46^.

**Figure 4:**
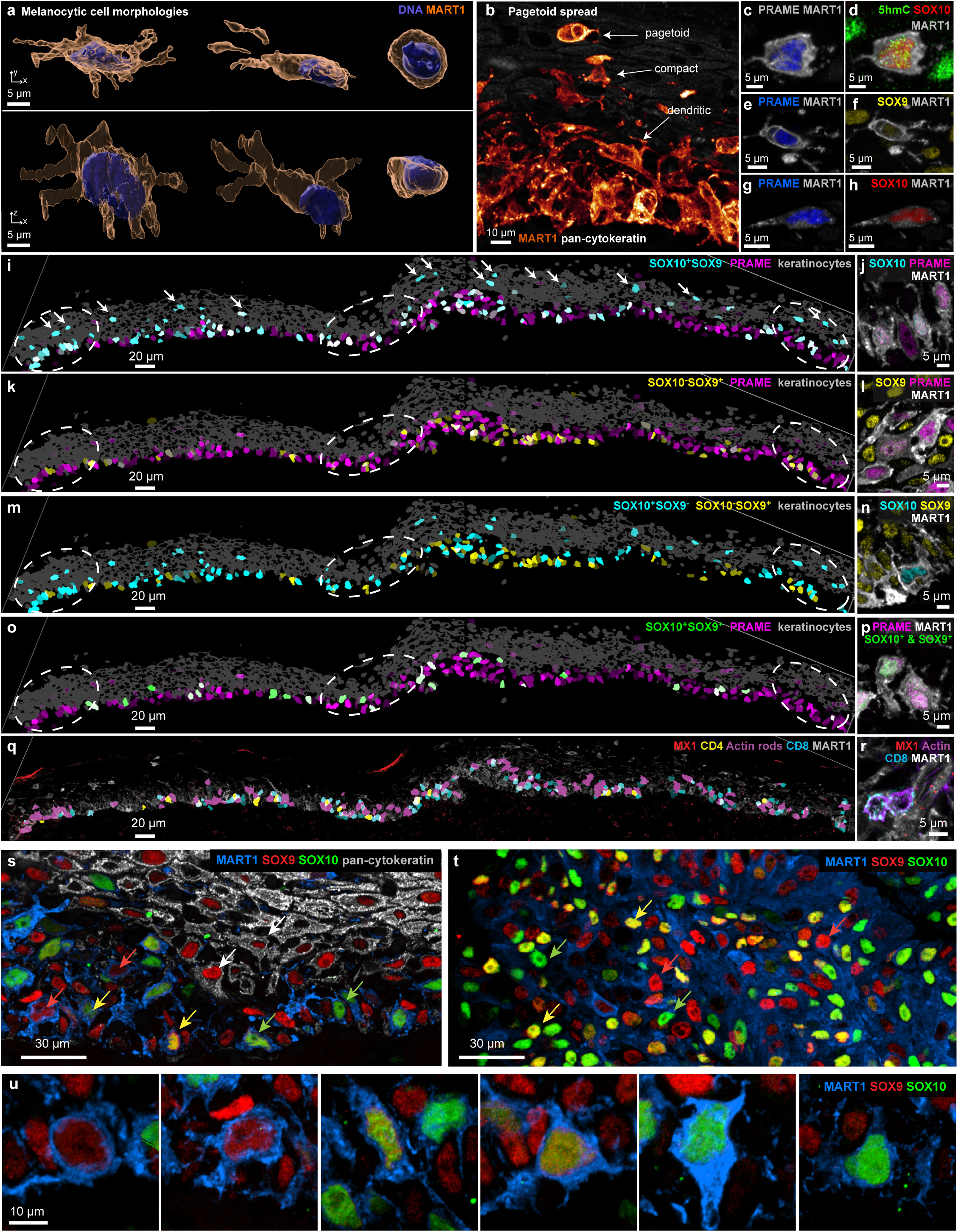
Melanocyte morphologies, lineage marker expression, and cellular interactions in the melanocytic intraepidermal compartment. **a,** Surface-rendered melanocytes within the MIS, illustrating variations in dendritic (normal) and rounded (transformed) morphologies. Scale bars, 5=µm. **b,** A representative FOV showcasing the transition of melanocyte morphology from dendritic-like at the DEJ to compact (bottom) and ultimately rounded during pagetoid spread within the epidermis (top). Scale bars, 10=µm. **c-h**, Representative examples of PRAME- and MART1-expressing pagetoid spread cells showing different expression levels of 5hMC, SOX9 and SOX10. Markers as indicated on each panel. Scale bars, 5 µm, **i-n**, Images of the MIS. Left column: segmentation masks coloured by marker intensity and brightness representing mean expression level. Masks in grey denote the positions of keratinocytes; dashed circles denote IFN-rich domains. Markers as indicated on each panel. Right column: high-resolution immunofluorescence images of the same markers per row. **o-p**, Images of the MIS as in **i-n**, but with colours indicating cells positive for PRAME (magenta) or cells dually positive for SOX9 and SOX10 (green). **q,r,** Images as in **l-p** but with magenta denoting cells containing nuclear actin rods. **s.** Maximum intensity projections of cells in the MIS showing gradations in SOX9 (red) and SOX10 (green) expression; red arrows denote differentiated MART1^+^ SOX10^+^ melanocytes; green arrows denote MART1^+^ SOX9^+^ melanocytic cells that have retained dendritic morphology; yellow arrows denote MART1^+^ melanocytic cells that co-express SOX9 and SOX10; white arrows denote keratinocytes (which are also SOX9^+^. Scale bar 30 μm. **t.** Similar data for tumour cells in the VGP. regions. Scale bar 30 μm. **u.** Magnified view of cells from panel s. Scale bar 10 μm.

We scored individual melanocytic cells (n=875; classified as MART1 or SOX10 positive, *see Methods*) in the MIS for expression of six markers of melanocyte lineage and transformation, including 5hmc, PRAME and MART1 (markers used clinically)^47^, SOX9, SOX10, and MITF (three transcription factors associated with melanocyte differentiation). 3D imaging made it possible to unambiguously score combinations of nuclear and cell surface markers at a single cell level. In melanocytic cells, the six markers we scored were present in nearly half of all possible cell type combinations (**Extended Data Fig. 5b**), without evidence of significant spatial correlation (**Fig. 4i-r**; see **Methods** for details on per channel gating). Cells undergoing pagetoid spread also expressed many different combinations of lineage markers (**Fig. 4c-h**) and transcription factors (**Fig. 4m,n**) but expression of NGFR (CD271), a marker of melanoma initiating “stem” cells^48^, was not detected. As a positive control, NGFR was detected in parallel thin sections of multiple metastatic melanomas.These data imply that melanocytic cells with features of early malignant transformation are subject to frequent changes in cell state (phenotypic plasticity) rather than progressive evolution from a single transformed or progenitor (stem-like) cell, as proposed for advanced invasive melanoma. The degree of plasticity may be greater than implied by **Extended Data Fig. 5b** since these data are binarized whereas images reveal graded transition between states (shown in **Fig. 4s-u** and **Extended Data Figure 5c** for SOX9 in red and SOX10 in green).

### Inflammatory neighbourhoods

The MIS contained multiple spatially distinct domains of inflammatory signalling, which were ∼50 to 100 µm in diameter and exhibited elevated levels or distributions of interferon (IFN) responsive proteins, such as IRF1, MX1, and MHC-I (**Fig. 5a-b, Extended Data Fig. 6a**). IFN domain size was confirmed on a distant serial section (**Extended Data Fig. 6b-c**). IRF1 is an IFN-responsive transcription factor that translocates from the cytoplasm to the nucleus and MX1 and MHC-I are downstream response genes; MX1 forms distinctive biomolecular condensates in the cytoplasm (often multiple condensates in a single cell)^36^ and MHC-I is found on the cell surface. Localized IFN-expressing niches have been previously described^49^, although the distribution of IFN within the tumour microenvironment is still a topic of active investigation^50^. Within these domains, melanocytic cells had started to pass through the DEJ and were in contact with immune cells (**Fig. 5b-c)**. Thus, our data provide direct evidence for restricted and recurrent spatial niches, defined by the simultaneous presence of an IFN response, melanocyte-immune cell contact, and melanocytes crossing the DEJ (the first step in invasion). These IFN-positive spatial niches were coincident with the lineage switching described above but without detectable spatial correlation, despite evidence that IFN can induce melanoma de-differentiation in cultured melanoma cells^51^.

**Figure 5:**
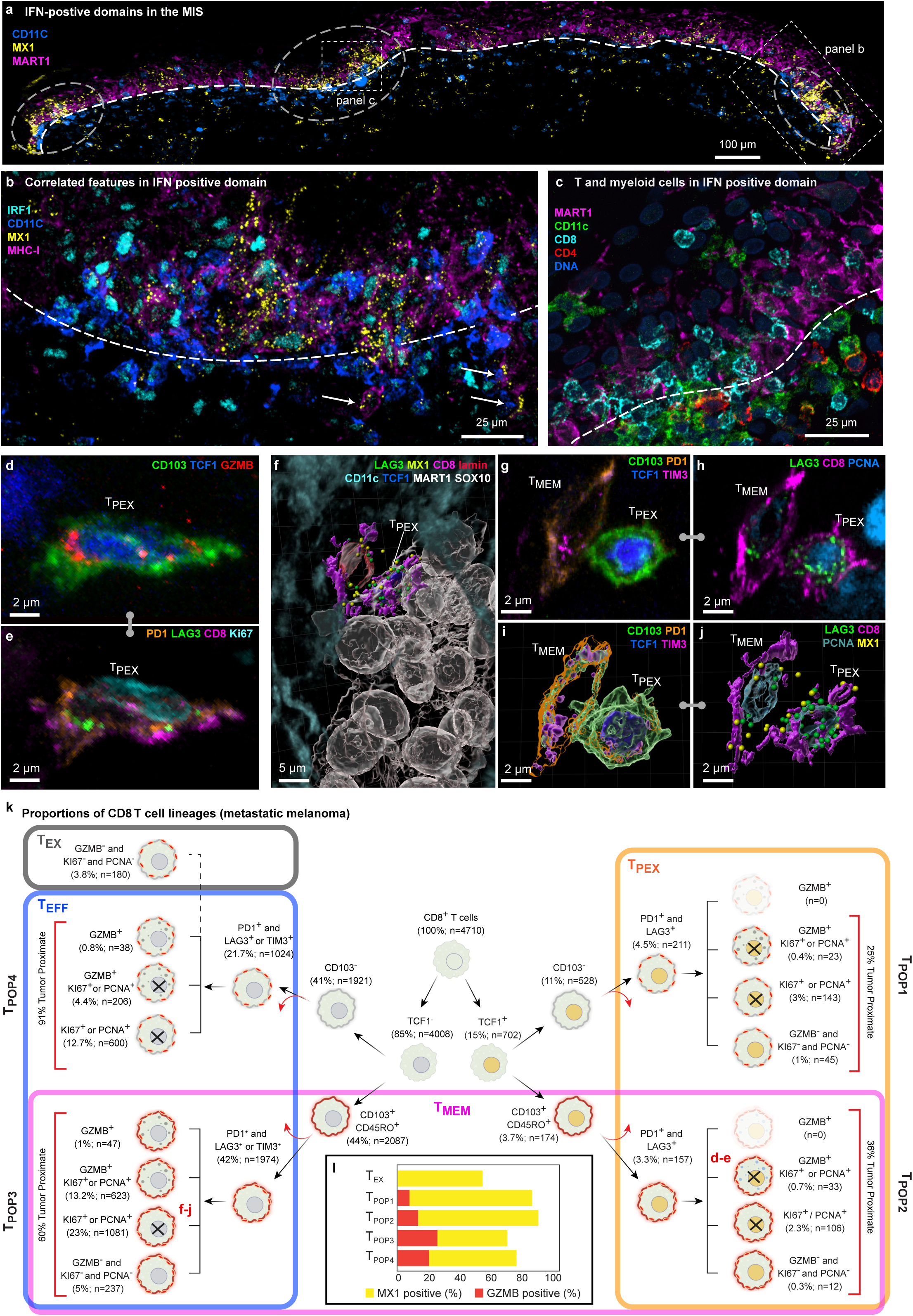
Spatial analysis of IFN-rich domains and distinct T cell lineages. **a,** Three selected channels of 54-plex 3D CyCIF image of the MIS in dataset 2 (LSP13625). A maximum intensity projection of 116 planes shown. IFN-rich domains denoted by dashed circles. DEJ denoted by white dashed line. Scale bar 100=µm. **b,** Magnified view of inset from panel **a** showing same cell nuclear localization of IRF1, expression of MX1, and MHC-1 upregulation. DEJ denoted by white dashed line. White arrowheads denote invasion of melanocytic cells into dermis. Scale bar, 25=µm. **c,** Enlarged inset from panel (**a**) showing diversity of immune cells crossing the DEJ. DEJ denoted by white dashed lines. **d-e,** Maximum intensity projection of an activated T_PEX_ cell, showing intracellular organelles like GZMB (**d**) and membranous proteins such as LAG3 and PD1 (**e**). **f,** 3D rendering of a T_MEM_ cell interacting with a T_PEX_ cell, which is in turn interacting with a cluster of metastatic melanoma tumour cells. Dendritic cells surround the neighbourhood. **g-j,** The T_MEM_ and T_PEX_ cells shown in (**f**), as a maximum projection (**g** & **h**) and 3D rendering (**I** & **j**). **k,** Hierarchical tree diagram showing proportions of CD8 T sub-lineages the metastatic melanoma specimen. T_MEM_ (magenta), T_PEX_ (orange), and T_EFF_ (blue) subtypes overlap, giving rise to four hybrid populations (T_POP1-4_) as denoted by vertical labels. See text for details. Red arrows denote additional cell subsets that are not shown on this tree. **l,** Percent of cells positive for GZMB (red) or MX1 (yellow) by population. See **Supplementary Figure 13** for a detailed diagram of which markers were used to define each cell type.

### Progenitor and effector T-cell subsets

The normal epidermis has an abundance of resident memory T cells (T_MEM_) as a consequence of prior encounters with non-tumour antigens (tissue-homing T_MEM_ cells are characterized in our data by expression of the lineage markers CD45RO and CD103)^52^. The presence of tumour leads to additional T-cell recruitment and activation. In-depth 3D immunoprofiling of metastatic melanoma using ten T cell lineage and state markers (n=4,710 CD8 and 2,820 CD4 cells) revealed a remarkable diversity of populations and states (**Fig. 5d-k, Extended Data Fig. 6d-e**). Among these, T_PEX_ cells^53–55^ (15% of CD8 cells; orange box; **Fig. 5k**) are of particular interest because they can be re-activated by immune checkpoint inhibitors (ICIs) and their presence is associated with improved patient outcomes^56^. These cells are commonly defined as CD8^+^ CD3^+^ T cells that co-express the master transcriptional regulator T Cell Factor 1 (TCF1)^57^ and checkpoint proteins (exhaustion markers) PD-1 and LAG3. In our data, T_PEX_ cells could be divided into two subpopulations based on expression of CD45RO and CD103 (T_MEM_; magenta box). Thus, T_PEX_ and T_MEM_ populations overlapped (giving rise to the hybrid populations T_POP1_ and T_POP2_; **Fig. 5k).** T_MEM_ cells also overlapped effector T cells (T_EFF_; blue box; defined as CD8^+^ TCF1^−^ PD1^+^ [LAG3 or TIM3]^+^ [Ki67, PCNA, and/or GZMB]^+^) and gave rise to hybrid populations T_POP3_ and T_POP4_. Terminally exhausted cells (T_EX_; grey box; defined as CD8^+^ PD1^+^ [LAG3 or TIM3]^+^ [Ki67^−^, PCNA^−^, and GZMB^−^]) were distinct from the four hybrid populations. Interestingly, we observed LAG3 puncta in all hybrid populations with T_MEM_ and CD103+T_PEX_ having the most (**Extended Data Fig. 4b**). Thus, in-depth phenotyping made possible by 3D imaging shows T_PEX_, T_MEM_, and T_EFF_ CD8 T cell populations overlap and possibly contain transitional states, consistent with a growing body of scRNAseq data (from chronic infections, for example)^53^. Our data show that it is possible to localize cells with these states in tumours and study their properties and interactions (**Extended Data Fig. 6f**).

Cytotoxic T cells were distinguished in our data by the presence of 1-20 GZMB puncta per cell (**Extended Data Fig. 4c-d**). Unexpectedly, GZMB^+^ T cells (5% of all CD8 T cells) were found to be both positive or negative for TCF1, CD45RO, and CD103 (**Fig. 5l** red bars). TCF1^−^ cytotoxic T cells were more likely to be GZMB^+^ and in contact with tumour cells than any other subtype, but visual review confirmed that all four populations included cells with GZMB polarized toward closely apposed tumour cells, implying active cell killing. Additionally, 40-80% of T_PEX_, T_MEM_, and T_EFF_ cells were PCNA or Ki67 positive, consistent with recent proliferation.^58,59^. Moreover, greater than 60% of all CD8 T cells contained multiple MX1 puncta, indicating an active response to IFN (**Fig. 5l**; yellow bars). These data are consistent with the known effects of IFN on T cell proliferation and suggest that this extends to all major T cell subtypes.

Spatial analysis showed that T_PEX_ cells were significantly closer to T_MEM_ and T_EFF_ cells than to T_EX_ or other T_PEX_ cells and that T_MEM_ cells were closer to tumour cells (**Fig. 5k, Extended Data Fig. 6g-i**). These data are consistent with evolution of T_PEX_ cells from a TCF1^+^ to a TCF1^−^ effector status in both the memory-related (T_POP2_ CD45RO^+^ CD103^+^) and classical (T_POP1_) populations (**Extended Data Fig. 6j-k**). Thus, the T_PEX_ cells in the tumours profiled here have properties consistent with three distinct populations representing (i) self-renewal, (ii) formation of TCF1^+^ cytotoxic cells, and (iii) differentiation into classic T_EFF_ cells.

### Membrane-membrane interactions

Existing approaches to proximity analysis use nuclear positions to identify cell-cell interactions^39^ but high-resolution imaging makes it possible to directly study more biologically-relevant membrane-membrane interactions. We observed three arrangements based on the degree of separation and extent of membrane-membrane proximity: (i) direct binding (Type I interaction), (ii) membrane apposition (Type II), and (iii) neighbourhood clustering (Type III). Direct binding involved pixel-level overlap in membrane proteins from neighbouring cells, and thus, co-localization at the resolution limit of the microscope. These Type I interactions were most obvious in the case of a CD8^+^ PD1^+^ T cells interacting with tumour and myeloid cells; **Figure 6a** shows this for a rare PDL1^+^ melanocytic cell. A membrane intensity profile spanning the a cell-cell junction on two apposed membranes followed by polynomial curve fitting, revealed a membrane-to-membrane spacing of ∼70 nm (**Fig. 6b**) vs. an average intermembrane spacing of ∼1.5 µm among all cells (see online methods ethods for a discussion of the impact of signal-to-noise ratio on these values). PDL1 expressing dendritic cells bound to CD8^+^ PD1^+^ T cells had a similar membrane spacing (**Fig. 6c,d**), as did a T cell interacting with a PDL1 negative melanocytic cell in the VGP (**Fig. 6e,f, Extended Data Fig. 7a**). Given the resolution limits optical microscopes, these Type I spacings are consistent with juxtacrine signalling and immunological synapses involving integrin-stabilised cell-cell contacts, which EM shows to have ∼30 nm membrane separation^60^.

**Figure 6:**
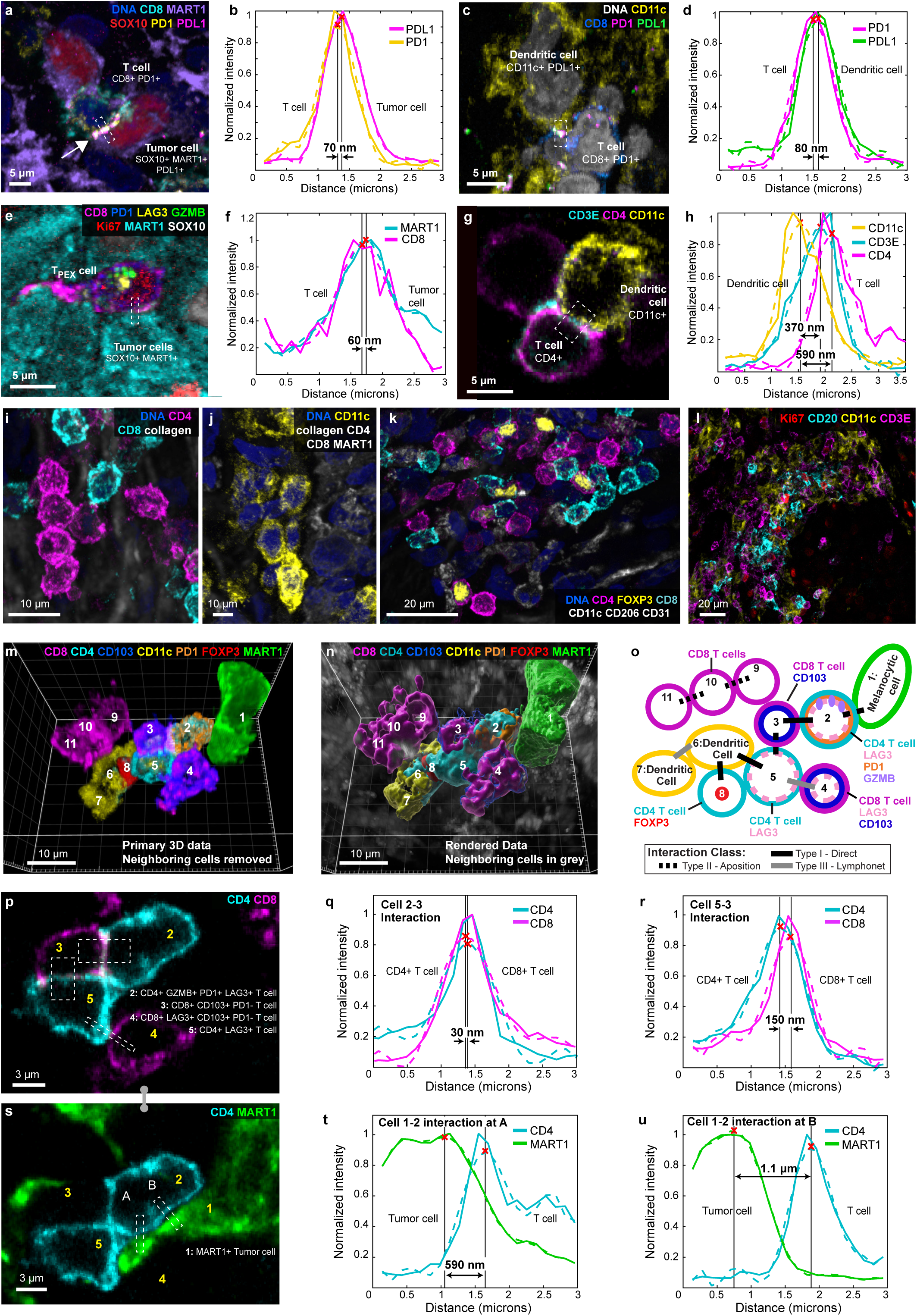
Cell-cell interactions and multivalent immune cell niches. **a-h,** Selected fields of view and intensity profiles for selected membrane-membrane interactions in dataset 1 (LSP13626). Scale bars 5=µm. **a,** A single image plane of a PD1^+^ CD8^+^ T cell interacting with two MART1^+^ tumour cells. White arrow denotes juxtacrine PD1-PDL1 interaction. **b**, Membrane intensity profile perpendicular to axis of interaction of membranes from cells 1 and 3 in **a** and demonstrating a Type I interaction. **c-d,** A dendritic cell interacting with a CD8^+^ T cell as a maximum projection (**c**) and membrane intensity profile (Type I interaction) (**d**). Box indicates region of line membrane intensity profile shown in **d**. **e,f,** Representations of a T_PEX_ cell from the invasive margin as a maximum projection (**e**) and membrane intensity profile (Type I interaction) (**f**), for region marked by white dashed box in I. **g,h,** A CD4 helper T cell (magenta) interacting with a dendritic cell (yellow), as a maximum intensity projection (**g**). Membrane intensity profiles are representative of Type II interactions (**h**). **i-k,** Examples of Type III interactions (lymphonets), involving CD4^+^ CD8^+^ dendritic cells in the MIS. These networks are characterised by loosely packed cells with cell-cell interactions via relatively small membrane domains. **l,** Stroma in the vicinity of the VGP melanoma showing neighbourhoods rich in CD20^+^ B cells, CD11C^+^ dendritic cells, CD3E^+^ T cells but without the clusters of proliferating Ki67^+^ cells that are characteristic of mature germinal centres. **m-o,** 11 cells from the MIS lying in proximity to the DEJ, shown as primary data (**m**), 3D surface renderings (**n**), and as a schematic representation of three Type I, four Type II, and two Type III interactions inferred from membrane intensity profiles in single-plane images (**o**). Community in (**m-n**) also shown in **Supplementary Video 5**. Scale bars,10=µm. **p,** A cross-sectional slice of two CD8^+^ and two CD4^+^ T cells depicting the direct engagement of cell membranes (see **Supplementary Video 4**). Scale bar 3=µm. **q,r,** A membrane intensity profile depicting CD4 and CD8 average expression within the bottom boxed region of **p** (**q**) or right boxed region of **r** (**s**). **s-u,** A Type II interaction between a MART1^+^ tumour cell and CD4^+^ T cells, as a representative image (**s**) and membrane intensity profiles for the interaction between cells 1-2 at region A (**t**) and B (**u**). Scale bars, 3=µm. For membrane intensity profiles, solid lines are from raw data, dashed lines from polynomial curve fitting. Red ‘X’s mark the maximum intensity along membrane intensity profile and thus, the centroid of the cell membrane for each channel. See **Supplementary Figure 13** for a detailed diagram of which markers were used to define each cell type.

Type II cell-cell interactions were characterized by neighbouring cells with extensive membrane apposition but without evidence of pixel-level overlap in protein staining; in this case a membrane-membrane spacing of 300-600 nm was typical (**Fig. 6g,h**). Type II interactions between CD4 and CD8 T cells were common across the MIS. In the conventional mode, APCs present antigens to both CD4 and CD8 T cells, with the CD4 helper cells enhancing the cytotoxicity of CD8 cells via cytokine production. In our data, CD4 and CD8 often exhibit Type II interactions without evidence of a nearby antigen presenting cell (e.g., dendritic cell, perhaps because such an interaction occurred earlier in time). Speculatively, Type II interactions may facilitate paracrine signalling.

Type III cell-cell interactions involved an intermembrane spacing of ∼500 nm but only along a small area of the membrane (1-2 µm^2^); such interactions may correspond to the tightly packed (jammed) arrangement of cells widely studied in biophysical models of tissue^61^. In some cases, 100 or more immune cells were observed to make Type III interactions, generating an arrangement we have previously described as lymphocyte networks (lymphonets)^62^. We observed lymphonets comprised primarily of CD4 T cells or CD8 T cells, dendritic cells, and mixtures thereof (**Fig. 6i-l, Extended Data Fig. 7b**). Lymphonets did not contain CD4^+^ FOXP3^+^ regulatory T cells, which were most commonly involved in Type I interactions (**Fig. 6k**), or tissue-resident macrophages, which were uniformly distributed across the dermis. Type III interactions among T, B, and dendritic cells were also observed in VGP melanoma (**Fig. 6l**) and may represent nascent tertiary lymphoid structures (TLS), which play role in responsiveness to immunotherapy^63^.

These three classes of membrane-membrane interaction frequently co-occurred. **Figure 6m-n** shows a complex set of cell-cell interactions involving a melanocytic cell at the DEJ (cell 1) and 10 immune cells (cells 2-11; **Fig. 6o, Supplementary Video 5**). In this network, a CD4^+^ GZMB^+^ memory T cell (cell 2) formed a Type I contact with a CD8^+^ LAG3^+^ CD103^+^ PD1^−^ memory T cell (cell 3; **Fig. 6p**) with an estimated membrane-membrane spacing of 30 nm over a 20-30 um^2^ area (**Fig. 6q)**. Cell 3 made an extended Type II contact with a CD4^+^ LAG3^+^ T cell (cell 5; 150 nm spacing **Fig. 6r**). Cell 4 and 5 engaged in a spatially restricted Type III interaction (640 nm spacing; **Extended Data Fig. 7c**). Cell 2 (CD4 T cell) also engaged in Type II contact with a melanocytic cell (1; 690 nm spacing; **Fig. 6s,t**). The two cells were proximate over a much larger area, but we judged the 1.1 µm spacing to be a consequence of tissue packing rather than interaction (**Fig. 6u**). Elsewhere in the network, Type I interactions were observed between CD4 T cell 5 and a dendritic cell (cell 6) and cell 6 and a CD4 T_REG_ cell (cell 8); finally, a CD8^+^ network (cells 9-11) extended in an epidermal direction (**Fig. 6o**). A caveat to this analysis is that distinctions among types of membrane interaction were not unambiguous and, given the dynamic nature of cell-cell communication, different interaction classes may represent different points in the time of evolution of a common structure. Regardless, our data show that immune cells form complex simultaneous associations with various cell types, including ones that send both positive and negative signals.

## DISCUSSION

Our data show that 3D high-plex imaging data of tissues and tumours substantially impacts how we understand the physical organization of tissues, assign precise single-cell phenotypes, and study multi-cellular communities (as compared to conventional 2D spatial proteomics). In particular, visualising the precise shapes of whole cells makes it possible to link form to function and study cell-cell interactions from the perspective of juxtaposed membrane-membrane contacts rather than mere proximity of cell (nuclear) centroids. High-resolution 3D imaging also overcomes errors in 2D image segmentation and corrects for misassignment of marker expression state due to protein polarization or overlap of cells along the imaging axis. To make high-plex 3D tissue imaging broadly available, online protocols (see Methods) provide detailed instructions for 3D tissue sectioning and imaging using a range of instrumentation, reagents, and public domain CyCIF methods. The data we provide also serves as a test bed for development of new 3D computational and visualisation tools (e.g. the recent mixed reality visualization of data from **Fig 1f** and **2a**^64^) and comparison with other methods in development^65^.

One striking result from imaging sections of varying thicknesses is that conventional 5 µm tissue sections contain few if any intact cells (or even nuclei) and this substantially interferes with detailed phenotyping of tightly packed tissues. However, by increasing section thickness only 4-5-fold (to a hydrated thickness of 30-40 µm), it is possible to overcome this inaccuracy and describe precise cell morphology, enumerate intracellular organelles, and classify cell types and states at a level of detail normally associated with cultured cells. For example, we find that at the earliest stages of melanocyte transformation, when cells lose their dendritic morphology, they express many combinations of differentiation and epigenetic markers, implying a high degree of plasticity in morphology and cell state. Plasticity is well-described in late-stage melanoma and cell lines^66^, but our data suggest it is also a feature of early stage disease. However, consistent with our prior observations^39^, primary melanomas generally lacked an NGFR-high state, which is often present in melanoma metastases and described as neural crest (stem-like) and tumour-initiating^48^. Thus, high cell plasticity may precede the appearance of NGFR-high, stem-like cells, rather than derive from them.

T cells in tissues are also revealed by 3D imaging to have a wide range of morphologies and states, with overlap between precursor, effector, and memory subtypes. This is also consistent with phenotypic plasticity and branching developmental trajectories. Immune cells are often components of complex communities of cells with tightly apposed membranes that appear to be involved in opposing regulatory signals: for example, a single cytotoxic GZMB^+^ CD8 T cell can be polarized toward a tumour cell, enveloped by filipodia from a CD4 helper cell, and repressed by a PDL1-expressing myeloid cell. Plasticity in tumour and immune compartments overlap (but does not obviously correlate) with spatially restricted inflammatory domains, which are often a few cell diameters wide. Spatially restricted cytokine signalling is predicted by mathematical models of cytokine production and uptake under conditions of high consumption^49^ and accurate membrane-level descriptions of tumour architecture should substantially assist in the development of these and other computational models of migration, signalling, and oncogenesis^67,68^. 3D thick section tissue imaging will also make it possible to study intracellular organelles, condensates, cytoskeletal structures, receptor-ligand complexes (including targets of therapeutic drugs), networks of filipodia and dendrites, changes in cell adhesion, and precise protein distributions in diverse cell types and tissues.

Understanding tissues in 3D must account for an ∼10^6^ range of length scales from intracellular organelles to entire organs; this will require integration of methods ranging from super resolution optical microscopy to microscale computed tomography (μCT). In general, as length scales increase, practically achievable resolution falls, as does the number of simultaneously addressable markers. The current work is specifically focused on the first part of the 3D continuum, namely high-plex subcellular imaging of intact cells, their organelles, and their neighbourhoods. Multiplexed imaging of ∼30 µm thick tissues has been described previously, primarily using flash frozen murine samples^5^ (rather than the FFPE human tissues that are available in clinical archives) but the impact of 3D imaging on the accuracy of downstream analysis has not previously been assessed. Next steps in the evolution of multiplexed 3D tissue imaging methods will include extending them to light sheet fluorescence microscopes able to image specimens up to a millimetre thick^17,69^ (currently in 3-4 channels), same-specimen μCT^24^ and 3D transcript profiling. These types of studies would make it possible to interrogate broader 3D architectures, for instance, whether the apparently separate interferon domains visible in Figure 5 are interconnected in 3-dimensional space (as has been shown for mucin pools in colorectal cancer^70^). It will also be essential to relate cell morphologies and interactions visible in high-resolution fixed cell images to functional states, most likely by parallel imaging of cultured cells, organoids, and mouse models subjected to genetic and immunological perturbation.

3D tissue imaging is not a replacement for existing 2D methods because it is harder to perform, generates substantially larger datasets, and is not as sparing of tissue (an important consideration with clinical samples). Instead, 3D data is likely to be most useful in detailed analysis of structures or cell states of particular interest (e.g. precise localization of potential drug targets), study of multiway immune cell communities, and generation of ground truth for training computational models able to discriminate otherwise ambiguous states in 2D images. As in stereology^25^, even a limited number of accurate 3D datasets on specific tissues and tumours will make it possible to correct for limitations in 2D images and identify those errors that confound accurate biological interpretation.

## Acknowledgements

We thank G. Guimaraes, T. Desai, and S. Fore from Carl Zeiss Inc. for providing access to the LSM980 Airyscan 2 microscope used to collect data in this proposal. We also thank T. Kupper, D. Liu, and J. Agudo for scientific advice, A. Chen, S. Chan, J. Muhlich, and J, Hoffer for help with data analysis, N. Ghelenborg and E. Moerth for access to the 3D Vitesse data viewer, J Lian for model training, and J Appelt for tissue integrity studies and the MicRoN core facility at HMS for assistance with microscopy.

## Author Contributions

CY, PKS, and AJN developed the concept for the study. CY, AW, ZM and PML collected image data. YL, ZS, FZ and GD developed software and performed data analysis in collaboration with CY, AJN, and JBT. SS GFM and CGL provided specimens and pathology expertise. CY, PKS, and JBT wrote the manuscript. All authors provided edits and approved the final manuscript. GD, GFM, CGL, SS and PKS provided supervision.

## Data Availability (At time of publication)

All primary images and derived data (∼5 TB) will available via AWS transfer at the time of publication. Instructions for accessing the primary and derived data is available via a data index page on Zenodo (doi.org/10.5281/zenodo.10055593). These images can be viewed using the free Imaris viewer (https://imaris.oxinst.com/imaris-viewer). 2D maximum projections of each dataset can be viewed in the MINERVA viewer (no download required), please see **Supplementary Table 1** for links. A subset of data will be available for 3D interactive viewing within the browser-based tool Vitessce (http://vitessce.io/)^71^. This effort is a work in progress and will be available in the future.

## Code Availability

Original code associated with this paper is available on GitHub (github.com/labsyspharm/mel-3d-mis) and Zenodo (doi.org/10.5281/zenodo.10055593).

## Extended Data Figure Legends

**Extended Data Figure 1: Imaging of thick tissue sections using 3D CyCIF**. **a-b,** Two methods for 3D imaging of thick tissue sections using high NA (high-resolution) oil immersion objectives. **a,** The standard arrangement in which a stained specimen is mounted to a glass slide and overlaid with a coverslip in 70% glycerol mounting medium. The coverslip is removed after each imaging cycle for fluorophore inactivation and another round of staining.^7^ This approach is often satisfactory but can result in damage to some tissues as cycle number increases. All data shown in the main figures of this paper was collected using this method. **b,** An alternative arrangement developed for imaging tissues prone to damage. The specimen is mounted to the coverslip and overlaid with Matrigel and a polyethylene mesh to hold the tissue in place, the assembly is fitted into a 3D printer holder and covered with a second coverslip to enable imaging with standard microscope slides. Use of this approach is demonstrated for a 35-micron section of colorectal cancer across 6 CyCIF cycles in **Supplementary Fig. 12a** and a melanoma precancer sample across 3 cycles in **Supplementary Fig. 12b**. **c,** Rendering of the holder used for specimen-on-coverslip imaging shown in **b**. **d,** UMAP rendering of all cell types analysed in Dataset 1 (LSP13626) as generated using 3D image segmentation algorithms. See **Supplementary Figure 13** for flow chart of cell type classifications. **e**, Difference in FFPE section thickness before and after hydration during antigen retrieval, as measured by confocal microscopy. Each datapoint represents measurement from 3 random positions. Error bars represent standard deviation (SD; n=3 per condition). **f**, Thickness of tissue sectioned in a frozen state, when fully-hydrated, after dehydration in ethanol plus xylene, and following rehydration. Thickness was measured at 3 random positions and error bars represent standard deviation. Statistical significance based on paired t-test. **g**. Height vs length ratio for computed bounding boxes covering ellipsoidal tumour cell nuclei in the VGP (having major and minor axes *l_1_* and *l_2_*) as viewed in the Z,X plane. As depicted in the inset, top-down views (along the Z axis) would produce a distribution of apparent lengths representing *l_1_* and *l_2_*; the fact that these are nearly equivalent in number in the data suggests no significant distortion along the imaging axis. **h**. Same analysis as **j** but viewed from the Z,Y plane. **i.** Percentage of cells from 35-micron Dataset (LSP13626) that would be incomplete in 9-micron virtual sections (positioned along the optical axis) compared to sections 18-35 µm thick. n represents the number of cells analysed. **j**, Percentage of cell volume missing when Dataset 1 was sectioned into virtual 9 µm thick sections for all cells, CD3^+^ T cells, and MART1^+^ tumour cells. Error bars represent the interquartile range. n value is the number of cells analysed. Note that tumour cell nuclei in this specimen are not much larger than immune cell nuclei; truncation of tumour cells would be more severe with large, pleomorphic nuclei. **k**, Average number of observed cell interactions between nearby cells as thickness of virtual section increases. **l**, Fraction of observed cell interactions with increasing virtual section thickness normalized to known number of cell interactions identified from full tissue thickness. **m**. Maximum intensity projection of Dataset 1 showing the cropped region used for analysis in **k** and **l**. **n,** Images of the same cut@5μm tissue section acquired with traditional widefield microscopy (left) and laser scanning confocal microscopy (right) with the same objective lens and microscope (40x/1.3NA, LSM980). Note that existing slide scanners operate at lower NA than the image shown here (typically 0.5 to 0.95 non-oil immersion objectives), resulting in higher signal-to-noise ratio in the widefield image. These data are quantified in **Supplementary Figure 22** to facilitate conversion of 3D confocal data into 2D representations.

**Extended Data Figure 2: Visualizing elongated immune and tumour cells in dense region from vertical growth phase. a,** 2D CyCIF whole slide image of adjacent section of the primary melanoma sample HTA7_1. White squares indicate the regions of melanoma in-situ (MIS) and the invasive vertical growth phase (VGP) melanoma where high-resolution 3D CyCIF was performed. Marker colours as indicated. Scale bar 1 mm. **b,** Maximum projection of the region of the invasive margin imaged with 3D CyCIF of Dataset 1 (LSP13626), showing a subset (6) of the total 54 markers. Image corresponds to right ROI indicated in (**a**). Scale bar 100 μm. **c-d,** Dendritic cell from **Figure 1j** with additional markers highlighting T cell subtypes (**c**) and the dense neighbourhood of tumour cells (**d**). See **Supplementary Figure 13** for detailed T cell subtype calling. **e,** Colour-coded segmentation masks of tumour cells from vertical growth phase of melanoma region. Colour encodes for orientation of tumour cells (spherical cells in yellow; elongated cells in magenta and cyan). Scale bar, 100=µm. **f, g**, Surface renderings of selected cells from (**e**). These cells differ in orientation in the tissue rather than aspect ratio.

**Extended Data Figure 3: Visualizing multi-cellular structures, cell shape, and motility in native tissue. a,** Surface rendering of 5-micron virtual section of the blood vessel from **Figure 2a**, viewed from above, showing B cell undergoing trans-endothelial migration (diapedesis). Scale bar 10 μm. **b,** Surface rendering of a different B cell (yellow) elsewhere in dermis of melanoma in-situ migrating through vessel wall (green) Scale bar 10 μm. **c,** T cell inside vessel, shown as a surface rendering (left) and as volumetric rendering (right). **d**, B cells in the VGP dermis with a characteristic round morphology. **e,** Sphericity (left) and volume (right) of B cells in the MIS (n=15) and VGP (n=352) regions. Error bars represent standard deviation.

**Extended Data Figure 4: Visualizing complex organelle and cell-surface morphologies. a,** 3D rendering of a neutrophil with organelles labelled. **b,** Left: Histogram displaying the frequency distribution of LAG3 spots per cell. The x-axis represents the number of LAG3 spots identified within individual cells, and the y-axis indicates the frequency of cells corresponding to each LAG3 spot count. Right: Distribution of LAG3 puncta by cell type. Shown as a boxen plot, where each box represents a quantile range, progressively detailing the distribution from the median outwards to the extremes. Open circles indicate outliers. **c,** Surface rendering of five interacting immune cells in the MIS, including three CD4^+^ helper T cells (magenta) and two CD8^+^ T cells (cyan). Left: MX1 biomolecular condensates (green), globular GZMB^+^ (yellow) in CD4^+^ T cells. Spacing between opposed membranes is <1.5 µm and contact area is ∼20 µm^2^. Scale bars 5=µm. Right: Reversed opacity of left image, showing punctate LAG3 (yellow) on the membranes of CD8^+^ T cells. **d,** Comparison of GZMB morphology (arrows) within a CD8 and a CD4 T cell in the MIS. **e-g,** Volumetric renderings showing PD1 and PDL1 can manifest as different morphologies within the same sample (Dataset 3 - LSP22409). **e**, Diffuse PDL1 on dendritic and CD8^+^ T cells. **f**, Diffuse PD1 and punctate PDL1 within the same dendritic cell. **g**, colocalization of punctate PD1 and PDL1 within the same dendritic cells. **h,** Activated LAG3^+^ GZMB^+^ CD103^+^ PD1^+^ CD8^+^ T cell with long filopodia (red) and dendritic cell (purple) interacting with a tumour cell. Note: This is a different cell community from that in Figure 2p-r. **i,** Same cells as (**h**) with GZMB and LAG3 shown, highlighting that the T cell is activated and cytotoxic. **j,** surface rendering of interactions in (**h**) and (**i**), showing the filopodia in greater detail. **k,** 3D rendering of cells highlighted in the multicellular interactions in **Figure 3f-g** and **Supplementary Video 6**.

**Extended Data Figure 5: Distribution of KI67 cells by phenotype in melanoma tissues. a,** Quantification of the Ki67+ cells within the MIS and VGP regions. Sample size is indicated above each bar. **b,** Bar graph showing the number of melanocytic cells in the MIS positive for each combination of six proteins (63 possible combinations): MART1 (green), PRAME (yellow), MITF (orange), SOX10 (red), SOX9 (blue), and 5hmc (violet). n=857. All categories with fewer than 15 cells were verified by manual inspection. Note different y-axes.

**Extended Data Figure 6: Spatial analysis of IFN-rich domains and distinct T cell lineages. a,** 2D CyCIF image of the MIS showing the correlation between pockets of MX1 (yellow) and IRF1 (cyan) along the dermal epidermal junction (dark blue). CD11c^+^ Dendritic cells shown in purple. Scale bar 50 μm. **b,** Quantification of MHC-1 expression in Gray Level Units (GLU) <1.5 μm (proximal) or >1.5 μm (distant) from an MX1 punctum in the DEJ of an independent dataset (dataset 1; LSP13626). Error bar indicates STD. **c,** Five selected channels from 42-plex CyCIF image of MIS in dataset 2. DEJ denoted by white dashed lines. White dashed rectangle is enlarged in **Figure 5c** and exemplifies CD45+ immune cells (green) breaking through the DEJ into the epidermis. Scale bar 30 μm. **d-e,** Two examples of maximum projections of TCF1^+^ T_PEX_ cells, showing non-proliferation (**d**) and proliferation via PCNA staining (**e**). Scale bar 5 μm. **f,** Hierarchical tree diagram showing proportions of CD4+ T cell sub-lineages in the metastatic melanoma sample. Red values indicate the percentage of cells in the associated state that are next to tumour cells. **g-k** Boxen plots of cell proximity in metastatic melanoma. Each box in a boxen plot represents a quantile range, progressively detailing the distribution from the median outwards to the extremes. Red boxes indicate cells positive for the given marker, blue indicate cells negative for the given marker, white indicate markers that were not used for selection. **g-h,** The distribution of the shortest distance between CD103^+^ (**g**) and CD103^−^ (**h**) T_PEX_ cells with other T cell subpopulations. **i,** The distribution of the shortest distance between tumour cells and T cell subpopulations. **j-k,** The distribution of the shortest distance between CD103^+^ (**j**) and CD103^−^ (**k**) T_PEX_ cells with subclasses of T_MEM_ populations, with activity states indicated by the marker patterns on the left.

**Extended Data Figure 7: Cell-cell interactions and multivalent immune cell niches. a,** Surface rendering of **Fig. 6e. b,** Maximum projection of tertiary lymphoid structures in the VGP tumour region. Dashed lines demarcate colonies of CD20+ B cells (cyan) mixed with CD11c+ dendritic cells (yellow). White box indicates zoom in version found in **Figure 6l**. Scale bar 100 μm. **c,** A l membrane intensity profile depicting CD4 (cyan) and CD8 (magenta) average expression across membranes of cells 4 (CD8 T cell) and 5 (CD4 T cell) from **Figure 6p**. Solid lines represent raw data and dashed lines polynomial curve fitting. Red ‘X’s mark the maximum intensity along membrane intensity profiles and denote the midpoint of the cell membrane for each channel.

**Extended Data Figure 8. Recurrent cellular neighbourhoods, identified by applying spatial latent Dirichlet allocation (LDA) to 2D projections of 3D data. a,** Scatter plot showing the MIS region. Cells are coloured based on their recurrent cellular neighbourhoods (RCN1–7). **b,** Scatter plot highlighting the distribution of each RCN in the MIS region of tissue. **c,** Scatter plot showing the invasive region of vertical growth phase melanoma. Cells are coloured based on the RCN to which they belong. **d,** Scatter plot highlighting the distribution of each RCN in the invasive region of tissue. **e,** Bar plot depicting the proportion of different cell-types within each RCN.

## Supplementary Videos

**Supplementary Video 1:** Surface rendering of Hoechst stained 5 and 35-micron thick serial sections from primary melanoma. 3D rendering from Bitplane Imaris 10.0.

**Supplementary Video 2:** Surface rendering of an intact blood vessel (green) from the MIS region with B cell (yellow), red blood cell (red), and neutrophil (pink). 3D rendering from Bitplane Imaris 10.0.

**Supplementary Video 3:** Volumetric rendering of packed tumour cells from the invasive margin. Rendering from Bitplane Imaris 10.0. Marker colours as indicated.

**Supplementary Video 4:** Surface rendering of CD4 (magenta) and CD8 (blue) positive T cells with LAG3 (yellow spheres), MX1 (green sphere), and GZMB (yellow blobs). Rendering from Bitplane Imaris 10.0.

**Supplementary Video 5:** Surface rendering of 1 melanocytic cell (green) and 10 interacting immune cells comprising of CD4 T cells (blue), CD8 T cells (magenta), and dendritic cells (yellow) from the MIS region. Rendering from Bitplane Imaris 10.0.

**Supplementary Video 6:** Mixed volume and surface rendering of cell community (from **Figure 3f-g**) in metastatic melanoma showing CD4 and CD8 T cells interacting with a tumour cell and dendritic cell. PD1 and PDL1 interaction can be observed in addition to GZMB in proximity to tumour cell. Marker colours as indicated. Rendering from Bitplane Imaris 10.0.

## Supplemental Tables

**Supplementary Table 1:** Sample metadata and related identifiers.

**Supplementary Table 2:** Definitions of all protein abbreviations.

**Supplementary Table 3:** CyCIF antibody panel used for LSP13626 MIS and VGP datasets, the first serial section from a patient with cutaneous melanoma.

**Supplementary Table 4:** CyCIF antibody panel used for LSP13625 MIS and VGP datasets, the first serial section from a patient with cutaneous melanoma.

**Supplementary Table 5:** CyCIF antibody panel used for metastatic melanoma (LSP22409 / WD-100476).

**Supplementary Table 6:** CyCIF antibody panel used for metastatic melanoma in lung (LSP22408).

**Supplementary Table 7:** CyCIF antibody panel used for glioblastoma (LSP17378).

**Supplementary Table 8:** CyCIF antibody panel used for serous tubal intraepithelial carcinoma (LSP18251).

**Supplementary Table 9:** CyCIF antibody panel used for tonsil (LSP13357).

**Supplementary Table 10:** CyCIF antibody panel used for fragile melanoma in-situ (LSP27564).

## Supplementary Discussion

### Approaches to Imaging 3D Tissue Specimens

This manuscript describes one way to acquire high-plex high resolution 3D images of thick tissue sections, but other methods are possible. One obvious approach is to collect a series of images from serial cut@5μm sections^24,70,72^, and then assemble 3D volumes. However, when the serial images are collected using standard slide-scanning microscopes, the resolution is relatively poor: ∼500 nm laterally and roughly the thickness of each section axially (e.g. ∼ 5μm). Each cutting plane also has the potential to introduce distortions. Thus, despite the potential advantages of using serial section reconstruction for multi-omic studies (e.g., alternating protein imaging and spatial transcriptomics across a 3D stack), such approaches are not truly 3D.

Multiple technologies exist for true 3D high-plex imaging. Selecting an approach involves balancing the number of fluorescence channels per cycle and the feasibility of performing cyclic staining with technical criteria like acquisition time per channel, contrast, and sensitivity (with good cameras, these parameters are determined in large part by the intrinsic efficiency of photon collection and the extent of rejection of out-of-focus light that would otherwise reduce contrast). As in any optical imaging system, resolution depends on wavelength, numerical aperture of the objective lens, axial sampling step size, and – in the case of a confocal microscope – pinhole diameter.

Shallow depth of field and effective rejection of out of out-of-focus light makes confocal microscopes an ideal way to perform optical sectioning on thick sections, and thereby, generate image stacks for 3D reconstruction. As an alternative, widefield microscopy can be supplemented with deconvolution methods to reassign out-of-focus signal back to the focal plane (based on knowledge of the point spread function) and improve the signal to noise ratio (SNR) and effective resolution. We have found deconvolution microscopy to be satisfactory with standard cut@5μm samples^39,73^ but less effective for thick sections, likely due to light scatter. Total internal reflection fluorescence (TIRF) microscopy^74,75^ can also be used produce 3D images in conjunction with techniques like DNA paint, but such methods are limited to a depth of ∼100 nm from the coverslip. Confocal microscopy can be performed using both spinning disk^76^ and laser scanning^77,78^ microscopes. We have tried both. The latter is slower due to the need to scan the image with a raster, but it removes pinhole crosstalk, provides finer control over the pinhole diameter, and thus, the resolution of optical sectioning (as a result, it also has higher contrast due to superior rejection of out-of-focus light).

In specialized research settings, two-photon excitation (TPE) microscopy^79^ uses pulsed mode-locked lasers to excite a femto-litre sized volume via the simultaneous absorption of two photons (typically in the near-infrared range). Two-photon lasers can also image certain collagen types that emit a label-free second harmonic signal^80–82^; the resulting signal has a wavelength that is precisely half of the incident laser wavelength. Since near-infrared wavelengths experience less attenuation and contribute to less phototoxicity, TPE is a preferred method for tissue and intravital imaging. Furthermore, multiple fluorophores can share similar two-photon excitation spectra, and this creates a throughput advantage in thick specimens. Recent research has shown that 3-photon excitation (3PE) microscopy enables even deeper imaging than TPE and is also compatible with live cell imaging^83^. However, TPE and 3PE systems are less common than confocal microscopes because they are significantly costlier and require specialist knowledge to operate and maintain. We exploit TPE in the current manuscript to image the collagen fibres that are major constituents of the extracellular matrix. We anticipate that other uses of TPE will emerge in tissue profiling but not as a general-purpose means of performing cyclic tissue imaging.

Overall, we conclude that confocal microscopy is the most promising approach to high-resolution imaging of tissue specimens up to ∼50 μm thick. However, confocal microscopes are relatively inefficient at collecting emitted light; orders of magnitude fewer photons reach the detector in a confocal than a widefield microcope^84^. Additionally, the full sample thickness (at any specific point in X,Y) is fully illuminated along the Z axis regardless of which focal plane is being imaged; this significantly contributes to photobleaching. These issues are known to be problematic with live-cell microscopy, but a key conclusion of our research is that they are not major issues in the performance of high-plex imaging of fixed tissue using antibodies and fluorophores.

Another promising imaging approach that emerged for our research is Light Sheet Fluorescence Microscopy (LSFM) and tissue clearing^69,85–88^. LSFM is an ideal way to image tissue sections as thick as several mm and the most recent systems are capable of cellular and subcellular resolution (ca. 0.7-2 μm), but existing clearing methods are generally incompatible with FFPE tissue^89^. Moreover the total number of fluorophores that can currently be imaged is typically 3-5 depending on the optics^90,91^, however, there are methods that demonstrate higher-plex 3D high-resolution (sub-micrometre) imaging in thick animal tissues^92,93^ or thin 5 μm sections^94^. We are currently developing an LSFM CyCIF protocol for millimetre-thick FFPE clinical samples and strongly believe that this will complement rather than replace the thick-section confocal methods described here.

## ONLINE METHODS

Detailed protocols for performing 3D thick section tissue imaging are available at dx.doi.org/10.17504/protocols.io.261ge59m7g47/v1 and will be updated as the methods improve.

### Specimen collection

See **Supplementary Table 1** for clinical metadata for all specimens. Specimens for melanoma (MIS and VGP), glioblastoma, lung metastasis, and tonsil were retrieved from the archives of the Department of Pathology at Brigham and Women’s Hospital and collected under Institutional Review Board approval (FWA00007071, Protocol IRB18-1363) under a waiver of consent. Serous Tubal Intraepithelial Carcinoma (STIC) samples were obtained from University of Pennsylvania after Institutional Review Board approval. Three datasets were used for the studies described in the body of the text: two 35 µm serial sections of melanoma (referred to as Dataset 1 (LSP13626) and Dataset 2 (LSP13625)) and a 35 µm section of metastatic melanoma obtained from the NIH Cooperative Human Tissue Network (CHTN) (referred to as metastatic melanoma or dataset 3; LSP22409 / WD-100476). Quantifications are based on Dataset 1. Deep immune cell phenotyping was based on features computed from Dataset 3. The histopathological regions of interest for each specimen were annotated as described previously^39^ by a board-certified pathologist using standard melanoma diagnostic criteria.

### 3D Cyclic immunofluorescence (CyCIF)

The procedure for 3D CyCIF is modified from the standard CyCIF^7^ protocol, with additional care taken during staining steps (see below). Staining plans containing lists of antibodies used with different specimens can be found in **Supplementary Tables 3-10**. Antigen retrieval, staining, and bleaching was performed as described previously^39^. Due to the fragile nature of thicker samples, extra care was taken during washes, bleaching and removing coverslips. Antibodies were diluted in 400 µl of blocking buffer and each section stained for 8-10 hours at room temperature to encourage penetration of antibodies but allow for same-day imaging. See **Supplementary Figures 1-11** for the whole slide images of the full dataset for all samples.

We found that most tissues held up well to these procedures, but that a subset of melanoma samples disintegrated during antigen retrieval. We have observed this previously with standard section skin and primary melanoma specimens but the reason why some specimens are more fragile than others remains unknown (such “pre-analytical variables” are common in histology). However, 3D imaging of several specimens (e.g. **Supplementary Figure 16**) revealed that the tissue had not fully adhered to the slide, instead exhibiting a series of corrugations that touched the slide in only some locations. Further research will be required to overcome this “corrugation” problem.

### Preparation of fragile samples

For fragile tissue specimens that adhere poorly to the microscope slides following dewaxing and antigen retrieval, we developed alternative approaches that did not require removing the coverslip between cycles, which we identified as one contributor to tissue degradation (**Extended Data Fig. 1a**). In this procedure, FFPE tissue sections were laid directly onto No. 1.5H grade glass coverslips (**Extended Data Fig. 1b, c**) and then stained, bleached, and imaged. This is ideal for the use of high numerical aperture objective lenses on inverted microscopes since such lenses are sensitive to coverslip thickness. We found that coating coverslips with poly-l-lysine overnight significantly reduced tissues from lifting off during dewaxing. We then glued a 1-mm thick spacer (cat. No.: IS003, SUNJin lab) around the tissue using acrylic glue thereby forming a well for the antibody solution or mounting media. Antibody incubation, imaging, bleaching, and washing were performed using the standard thick tissue CyCIF approach. During imaging, the well spacer was filled with 70% glycerol and covered with a second coverslip to reduce evaporation. We also explored the use of overlaying 400-500 µl volume of Matrigel (mixed with PBS at 1:1 volume ratio) or a black poly-ethylene mesh (with 7 square holes per inch) on the tissue within the spacer. Both helped to protect the specimen while not interfering with antibody staining. The entire assembly can either be inserted into standard microscope stage inserts or fitted into 3D-printed slide-shaped holders for more convenient handling. Although this second approach to sample preparation requires some optimization and user training, with the use of pre-printed components, the additional setup time is insignificant compared to the per cycle 3D image acquisition time.

### Optimizing sample thickness

To determine an ideal tissue thickness for CyCIF imaging we used tonsil tissue. Based on the maximum working distance of most water and oil-immersion lenses (∼200 µm) and the thickness of a no. 1.5H coverslip (170 µm), we selected tonsil sections that were cut@10 µm, 20 µm, 30 µm, 35 µm, and 40 µm thick (**Supplementary Fig. 14).** These were stained with Hoechst and gamma-tubulin conjugated in Alexafluor 555. Gamma-tubulin is punctate and serves as a useful stain for assessing antibody penetration and image aberrations. Z-stacks were acquired from each stained tissue sampled at 103 nm laterally and 230 nm axially using a 40x/1.2W C-Apochromat water immersion objective lens on a Zeiss LSM980 confocal microscope. We observed punctate gamma-tubulin in all thicknesses up to 35 µm tissue thickness, with uniform intensity along the axial axis (**Supplementary Fig. 14a-d**). However, at 40 µm thickness, gamma-tubulin intensity significantly diminished along the axial axis (**Supplementary Figure 14e**) and contrast (even along the top surface) was poorer than with thinner samples. We speculate that standard dewaxing and antigen retrieval protocols were not working well at tissue thicknesses greater than cut@35 µm. Moreover, we also observed signal attenuation in the Hoechst channel in cut@40 µm specimens. In this case, poor penetration of short wavelength light is likely an issue. Based on these considerations, we concluded that confocal imaging of thick sections is likely to be most effective with samples thinner than cut@30-35 µm.

### Variables in antibody staining

We sought to identify the shortest antibody incubation time needed to homogenously stain a thickness of 35 µm. Unlike 2D widefield imaging where overnight incubation times are often conveniently used, 3D imaging takes significantly longer and we want to accelerate the overall process **(Supplementary Figure 15).** Four 35 µm human colorectal cancer tissue specimens were stained with a cocktail of primary conjugated antibodies for 1, 2, 4, and 8 hours at room temperature. Z-stacks were acquired with voxel size (400nm x 400nm x 290nm) to reduce file sizes, and the depth of antibody penetration was assessed using orthogonal views in Imaris software. After 8 hr of staining, we observed that E-cadherin, CD11c, CD3E and MX1 had homogenously penetrated the entire thickness of the tissue. E-cadherin staining was complete within 2 hours. However, vimentin staining was limited to the top and bottom surfaces of the tissue and penetration did not improve with longer incubation times. We suspect this could have to do with the relative distribution of protein across the tissue thickness. It is expected that vimentin (which is expressed by many diverse cell types) would be present in high concentrations compared to other markers (e.g., MX1, CD3E). A high-degree of antibody binding at the outer surfaces could lower the effective concentration of antibody within the centre of the tissue, as has been described elsewhere^89^.

We also evaluated whether certain fluorophores impacted antibody penetration. This is important for CyCIF where the ability to choose different antibody fluorophore combinations is essential. We obtained a primary melanoma and co-stained MART1 conjugated to Alexafluor 647 with other secondary antibodies (Alexafluor 488, Alexafluor 555, Alexafluor 750) (**Supplementary Fig. 16a**) for 8-10 hours at room temperature. We bleached MART1-647 and restained with Alexafluor 647 in a subsequent cycle. **Supplementary Fig. 16b** shows that the MART1 primary conjugate (magenta) penetrated the full thickness of the tissue, as judged by Hoechst staining (turquoise). **Supplementary Fig. 16c-f** shows that all secondary antibodies (magenta) penetrated equally well and showed a similar staining pattern to the MART1 primary conjugate. This demonstrates the ability for secondary antibodies to be used for thick tissue CyCIF. We noted that Alexafluor 750 had lower contrast, which can be attributed to the lower sensitivity of detectors in the near infrared spectrum.

While testing multiple primary conjugated antibodies, we observed antibody penetration issues with some antibody conjugates. Although many immune markers (PD1, CD11c, CD8a, MHC-1, MHC-II; green) exhibited full depth staining, several tumour and stromal markers (αSMA, PCNA, SOX10; red) only stained the top layer of tissue (**Supplementary Fig. 16**). To determine whether the fluorophore played a role in this, we repeated staining with the same PCNA clone conjugated to Alexafluor 488 or Alexafluor 750. We noticed there that was a difference in staining pattern; the Alexafluor 488 conjugate-stained fewer cells (**Supplementary Fig. 18a**) but showed improved staining penetration (**Supplementary Fig. 18b**). For αSMA, we tried a similar strategy, but using a different fluorophore required a different antibody clone. Unlike PCNA, we did not see an improvement in staining penetration of a blood vessel (**Supplementary Fig. 19**). From these data we concluded that antibody penetration is not uniquely dependent on fluorophore or clone but is influenced by multiple factors and that each antibody must therefore be evaluated for its ability to stain a thick section using Z-stacks.

### 3D image acquisition

All image data was collected on a LSM980 Airyscan 2 (Carl Zeiss) equipped with a 405nm, 488nm, 561nm, 647nm, and 750nm laser lines, and 5x/0.16NA air, 10x/0.45NA air and 40x/1.3NA oil immersion objective lenses. Microscope slides were secured in a slide holder fitted with a spring-loaded clamp, which correspondingly was secured onto the microscope stage in a plateholder. In ZEN 3.9, a 2D overview scan of Hoechst using the 5x objective lens was used to identify regions of interest for higher resolution imaging at 40x in 3D. Images were sampled at 16-bit at 0.14 microns per pixel in X and Y, and 0.28 microns per pixel in Z for approximately 170 or more optical planes. The pinhole size was set to 35 microns. A focus surface was used to maintain focus. To increase throughput, bidirectional and fast frame scanning was used. Channels were separated into two tracks: track 1 - Hoechst, Alexafluor 555, and Alexafluor 750 (if present). track 2 - Alexafluor 488 and Alexafluor 647. The emission range for Hoechst, Alexafluor 488, Alexafluor 555, Alexafluor 647, and Alexafluor 750 were 380nm-489nm, 499nm-544nm, 579nm-640nm, 660nm-705nm, and 755nm-900nm respectively. We note that the current work does not fully exploit the spectral unmixing capabilities of the LSM980^95–97^ due to a requirement for additional panel optimization. However, better spectral unmixing in the future is expected to reduce the number of cycles required to collect high-plex data in the future.

Type I and II collagen were imaged using Second Harmonic Generation (SHG) in a Stellaris 8 DIVE coupled to an Insight X3 multiphoton laser and running LasX. Images were acquired with a 20x/0.75NA multi-immersion lens and sampled at 0.36 microns laterally and 0.95 microns axially. SHG signal was detected using 4Tune Spectra non-descanned HyD detectors and separated from that of Hoechst 33342 using Fluorescence Lifetime Imaging Microscopy (FLIM).

### 3D image processing and registration

To improve signal-to-noise, all data acquired on the Zeiss LSM980 were processed using Zeiss ZEN LSM Plus Processing. Channels were background subtracted by removing a fixed constant grey-level from the background. The first cycle was stitched in ZEN using the Hoechst channel as a reference, and all subsequent cycles were registered to this first stitched cycle. Single-field and stitched 3D datasets were imported using Bioformats in MATLAB (Mathworks). First, the X and Y translations were obtained using max projections of the Hoechst nuclei channel. Following this transformation, subsequent cycles were registered in Z. We found that separating the lateral from axial transformations was more accurate than registering X, Y, and Z in one optimization step. We then performed histogram equalization with MATLAB’s *histeq()* function and fine-tuned image alignment with elastic deformations using MATLAB’s *imregdemons()* function. Lastly, all transformations for each cycle were applied to their corresponding channels. Each channel was saved and appended to a TIFF file and visualized in Meshlab, ChimeraX or Imaris 10.0 (Bitplane) as .ims files. We regard these as interim methods that will benefit from additional automation and refinement in the future.

### Single-Cell Phenotyping

Manual gating was performed for each marker to differentiate background from true signal. All antibodies had been validated in our laboratory; true signal was determined by comparing signal from a positive control tissue. The gates identified for each marker were subsequently used to normalize the single-cell data within a range of 0 to 1, wherein values above 0.5 indicated cells expressing the marker. The scaled data was subsequently used for phenotyping the cells based on known lineage markers as described previously using the SCIMAP Python package (scimap.xyz).^39^ See **Supplementary Figure 13** for the detailed marker combinations used to define cell types.

### RCN Analysis to Identify Microenvironmental Communities

The Latent Dirichlet allocation (LDA) based recurrent cellular neighbourhood (RCN) was performed using SCIMAP (scimap.xyz)^39^ using a k value of 10 (**Extended Data Figure 8)**. The clusters were manually organized into meta-clusters (7 clusters), based on the cellular composition of the clusters. The meta-clusters were also overlaid on the H&E and CyCIF images to validate their characteristics. For instance, RCN1 typically aligned with areas known to be tumour domains, while RCN2 was more closely associated with the epidermis, thereby highlighting the structural elements within the dataset.

### Statistical Tests

All statistical tests were performed using MATLAB’s *ttest2* implementation of the two-sample t-test without assuming equal variances and significance value of p<0.05.

### Cell type calling in virtual thin sections

Three sections of varying thicknesses (cut@5, 10, 20, and 35 microns) were cut from a FFPE block, processed, stained with Hoechst and imaged as described above. The mean thickness of random regions of interest were measured using orthogonal views in Imaris. We observed that if a FFPE section was cut at 5 microns, it will inflate to 9 microns after rehydration. Therefore, from our datasets and corresponding cell masks, we created virtual 9-micron thick serial sections and compared them with the entire 35-micron thick section. To enumerate the percentage of incomplete cells in these thinner sections, we used MATLAB’s regionprops3 boundingbox to find cells that had top or bottom faces coinciding with the top or bottom of the virtual section respectively. The volume property was used to compare the average cell volume that would be missing from each virtual section from its corresponding whole volume in the 35-micron section. To determine degree of cell type miscalling in thin sections, we performed spot counts of LAG3, GranzymeB, and MX1 and compared to the corresponding thick section. A cut-off of >2 spots was used to identify positivity in both thin and thick sections. We calculated the percentage of positive cells that were misclassified as negative cells due to spot exclusion in a virtual thin section. Graphs were plotted in R showing mean and interquartile range.

### Cell interaction analysis in virtual thin sections

A densely packed volume of 56 x 56 x 35 microns was selected and cropped from the epidermis region of Dataset 1-melanoma in-situ for cell interaction analysis and 3D segmentation. From each cell, we measured the cell centroid and the distance to the nearest unobstructed neighbouring cell edge/membrane to identify neighbours. We further used a minimum distance cut-off of 2 microns to identify neighbours that involved membrane contact. The number of missing neighbours in all virtual sections was normalized to the number of neighbours in the full tissue thickness, which was taken as ground truth ([number of neighbouring cells in 2D] / [number of neighbouring cells in 3D]).

To test the ability of detecting neighbour cells in 2D optical planes, we extracted single Z-planes spaced 3 planes apart. We filtered out cells with an area of less than 2 µm^2^ (25 pixels). For thicker sections, we started from the middle Z-plane and systematically increased the thickness by 12 planes. We filtered out cells with a volume less than 2.7 µm^3^ (125 voxels). Finally, we also tested a maximum projection of a 9-micron virtual section to simulate widefield imaging of a traditional 5-micron section (adjusted for hydration). Graphs were plotted in MATLAB (Mathworks).

### B cell sphericity analysis

B cells in the melanoma in-situ were segmented in 3D in Imaris using the Surfaces module with a smoothing of 0.14 microns/per pixel. Non-cellular bright objects were manually removed. Since B cells were more compact in the vertical growth phase, we used Arivis’ Cellpose Cyto2 model implementation based on the membrane marker MHC-1 and a cell diameter of 8 microns. B cells were manually selected by having a CD20 mean intensity above 750 gray level units and a minimum volume of 25 micron^2^. In both software, the calculation for sphericity is the same and was based on each cell’s mask instead of voxels.

### Calibration Curve of FFPE tissue thickness

FFPE mouse thymus tissue was sectioned at different thicknesses of 5 µm, 10 µm, 20 µm, and 35 µm. We cut three sections for each thickness and stained each with DAPI and LDS751 (ThermoFisher Scientific) overnight in PBST (0.1%). A 3D stack of each tissue section was imaged in 70% glycerol (n=3). The thickness was measured in the XZ orthogonal view in ImageJ with the merged channels. The regression analysis was performed using measured thicknesses as the dependent variable and set nominal thicknesses on the microtome as the independent variable. Mean values were calculated for each nominal thickness, with standard deviation used to construct error bars on the calibration plot. The analysis was carried out using Python (version 3.9.16) and the scikit-learn package (version 1.4.1.post1), and the calibration plot was visualized using the matplotlib package (version 3.4.3).

### Segmentation of individual 3D cells with Cellpose

Individual 3D cells were segmented from the dense tissue volumes using a new modification of Cellpose designed for 3D data^21^. The original 2D Cellpose model (https://github.com/MouseLand/cellpose),^98^ is a custom gradient tracking approach that aggregates x-y, y-z, x-z 2D slice cell probability and gradient maps predicted by pretrained 2D segmentation models. The full Cellpose segmentation framework, suitable for a wide range of 3D cell imaging data along with in-depth validation and determination of method applicability specific to this project, is described below.

#### Image preprocessing for Cellpose

The 3D volumes were acquired at voxel resolution of 140 x 140 x 280 nm. For each 3D channel image, we resized the x-y slices by half to obtain isotropic voxels. The raw image intensity, 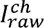 was then corrected for uneven illumination, 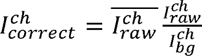 where 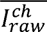 is the mean image intensity and 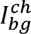 an estimation of the background illumination obtained by downsampling the image by a factor of 8, Gaussian smoothing with sigma = 5 and resizing back to the original image dimensions. 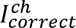 was then contrast-stretched to a range of 0-1, clipping any intensities less than the 2^nd^ percentile to 0 and any greater than the 99.8^th^ percentile to 1. Cellpose uses a single channel cytoplasmic and nuclear signal for two-color based cell segmentation. The mean of the intensity-normalized, background-corrected HLA-AB, CD3E, CD11b, and β-actin channels was used as the cytoplasmic signal. DAPI was used as the nucleus signal. Both cytoplasmic and nucleus signals underwent a further round of background correction and contrast stretching as described above before being concatenated to form the input RGB volume image.

#### Initial Cellpose 2D segmentation

The RGB volume was input slice-by-slice to Cellpose 2D in three different orientations: x-y, x-z, y-z to obtain three stacks of cell probability and 2D gradients. The performance of Cellpose depends on appropriate setting of the diameter parameter which relates to the size of the cells to be segmented. As the appearance of the cells may vary depending on orientation, we conduct a parameter screen with diameter = [10,100] at increments of 5 using the mid-slice for each orientation. At each diameter we compute the ‘sharpness’ of the predicted gradient map as the mean of the image variance evaluated over a local 5×5 pixel window in both ‘x’ and ‘y’ gradient directions. The diameter maximizing the variance after a moving average smoothing with window size of 3 was used to run Cellpose 2D on the remaining slices in the orientation. The raw cell probability output, *P* from Cellpose are the inputs to a sigmoid centered at zero, 1/(1+*e*^−*P*^). This means the probabilities vary predominantly linearly in the range −6 to +6 and this reduces the distinction between foreground and background. Thus, we clip the probabilities to the range [-88.72, 88.72] (to prevent overflow or underflow in float32) and convert back to a normalized probability value in the range 0-1 by evaluating the sigmoid, 1/(1+*e*^−*P*^). The probabilities from all 3 orientations are combined into one by averaging. Similarly, the 2D gradients are Gaussian smoothed with sigma=1 voxel and combined into a single 3D gradient map. Gradients are then normalized to be unit length. Lastly, we perform 3-level Otsu thresholding on the combined probability map and use the lower threshold to define the foreground binary voxels for gradient tracking.

#### Aggregating Cellpose 2D predictions

The volume was divided into subvolumes of (256, 512, 512) with 25% overlap. Within each subvolume we run gradient descent with momentum for 200 iterations, momenta, *µ* = 0.98, step size *δ* = 1 to propagate the position of foreground pixels towards its final attractor in the 3D gradient map.

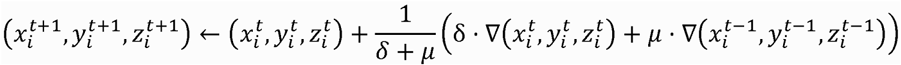

Here 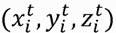 denotes the coordinate of foreground voxel *i* at iteration number *t*, *µ* the momentum ranging from 0-1, δ the step size and ∇ is the gradient map. Nearest neighbor interpolation is used, thus 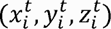 is always integer valued. Gradient tracking of all subvolumes are conducted in parallel using multiprocessing. The final coordinate positions from all subvolumes are compiled. We then build a volume count map where voxels mapping to the same final coordinate adds +1 to the count. The count map is Gaussian smoothed with sigma=1 and binarized using the mean value as the threshold. Connected component analysis identifies the unique cell as clusters where foreground voxels have been mapped to the same cell. Transferring this labelling to initial voxel positions 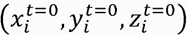 generates the individual 3D cell segmentations.

#### Postprocessing 3D cell segmentations

Small individual cell masks (<1000 voxels^3^ ≈ 20µm^3^) were first removed because they corresponded to debris. We also removed all cell masks that do not agree with the Cellpose predicted 3D gradient map. This is done by computing the 3D heat diffusion gradient map given the computed 3D cell segmentations and computing the mean squared error (MSE) with the input combined Cellpose 3D gradient map for each cell. Cells with MSE > 0.8 were discarded. Cells that are implausibly large, with volume greater than the mean volume ± 5 standard deviations were also discarded.

For the remainder cells, we run a label propagation^99^ to enforce that each segmented cell mask comprises only a single connected component and to denoise the masks. This is done for each cell mask, *M_i_*, by cropping a subvolume, *V_i_*, the size of its bounding box padded isotropically by 25 voxels. Each unique cell region is represented as a positive integer label. Every label in *M_i_* is encoded using a one-hot encoding scheme to create a binary column for each unique label. This generates a label matrix, 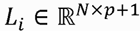 for *V_i_* where *N* is the total number of voxels and *p* the number of unique labels in *V_i_* and one additional label for background. We then construct the affinity matrix, as a weighted sum (α = 0.25) of an affinity matrix based on the intensity difference in the cytoplasmic signal between 8-connected voxel neighbors, *A_intensity_*, and one based on the connectivity alone, 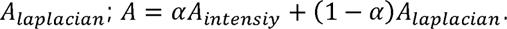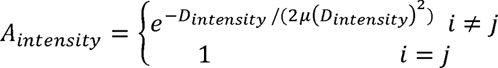. where *D_intensity_* is the pairwise absolute difference matrix between two neighboring voxels *i* and *j*. 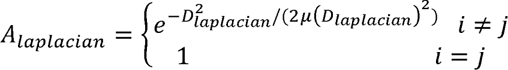 where *D_laplacian_* is the graph Laplacian with a value of 1 if a voxel *i* is a neighbor of voxel *j*, and 0 otherwise. *µ*(*D*) denotes the mean value of the entries of matrix *D*. The iterative label propagation is

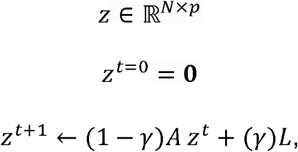

where *t* is the interation number, 0, denotes the empty vector and γ is a ‘clamping’ factor that controls the extent the original labeling is preserved. We set γ = 0.01. We run the propagation for 25 iterations. The final *z* is normalized using the softmax operation and argmax is used to obtain the final labels. The refined cell mask, 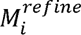 is defined by all voxels where *z*, has the same cell label *i.* All postprocessing steps were implemented using parallel multiprocessing iterating over individual cells.

### Comparison of tissue thickness before and after hydration, dehydration and rehydration

#### Tissue Preparation and 3D image acquisition

Kaede mouse thymus tissue was fixed in 4% paraformaldehyde (PFA) and stored in PBS at 4°C. The tissue was embedded in 8% agarose for vibratome sectioning (Leica VT1000 S) at a thickness of 35 µm and mounted on frosted glass slides. The section was stained with Hoechst 33342 and imaged in the hydrated state. The tissue was then mounted with 70% glycerol and imaged with a high-precision coverslip No. 1.5H (ThorLabs) on a Zeiss LSM980 confocal microscope with an EC Plan-Neofluar 40x/1.30 oil DIC M27 objective.

For tissue dehydration, the same sections were dehydrated through a series of ethanol solutions (50%, 70%, 95%, two changes of 100%) and xylene (two changes). The dehydrated sections were then mounted in a toluene based mounting media (Permount, Fisher Scientific) and imaged again. Subsequently, the sections were decoverslipped in xylene for 2 hours and then rehydrated through two changes of xylene and graded ethanol back to PBS (as previously described). The rehydrated section was imaged in the same condition as the hydrated state in 70% glycerol as previously described.

#### Image Processing, Tissue Thickness Measurements, and Statistical Analysis

Z-stacks were observed in ImageJ (version 1.54f) as XZ orthogonal views to measure tissue thickness. We selected an orthogonal plane from each of the three conditions (hydrated, dehydrated, rehydrated) and measured the tissue top and bottom in three locations. These were selected to be approximately at the same locations between the tissue sections for consistency.

Tissue thickness measurements were compared across three hydration states of the same sample: hydrated, dehydrated, and rehydrated. Paired t-tests was conducted using the scipy.stats package (version 1.12.0) to assess the effect of hydration state on tissue thickness. To account for multiple comparisons, a Bonferroni correction was applied, adjusting the significance level to α = 0.0167. The paired t-tests revealed that hydrated tissues had significantly greater thickness than dehydrated tissues (t = 8.194, P = 0.015), while there was no significant difference between hydrated and rehydrated tissues (t = −0.507, P = 0.663). Additionally, dehydrated tissues had significantly lower thickness compared to rehydrated tissues (t = −7.124, P = 0.019). Error bars in the accompanying figure represent the standard deviation (s.d.) for each treatment group. The analysis was visualized using the matplotlib package (version 3.7.1), and data manipulation was performed using the pandas package (version 2.2.1). The entire analysis was carried out in an environment running NumPy (version 1.24.3) in Python (version 3.9.16).

### Membrane-Membrane interaction analysis

To estimate the distance between membranes of adjacent cells, we computed membrane intensity profiles by integrating the fluorescence intensity in each channel parallel to the cell membrane (dimension y) over a distance of 5-10 pixels (0.7 to 1.4 μm) and then plotting it as a function of distance perpendicular to the membrane (dimension x) over a distance of 20-40 pixels (2.8 to 5.6 μm). This approach increased the signal to noise ratio relative to a simple line integral. The position and distance over which membrane intensity profiles were computed is depicted by rectangles in **Figure 3** & **6** and **Supplementary Fig. 20** & **22**. Both the position and dimension of the rectangles were determined visually based on the extent of cell-cell contact with the goal of characterizing local contacts at reasonable SNR. We then performed polynomial curve fitting on the resulting profiles to estimate the distance between the peaks of membrane staining. The first and second derivatives were obtained from the fitted curves to determine the roots, and the root at the maximum peak of the fitted curve was used to localise the cell membrane more precisely. As described previously, sub-pixel precision can be obtained using this approach^100^. To determine the robustness of these estimates to limitations in the imaging data we performed a range of studies on prototypical membrane-membrane interactions, as follows

#### Effect of spatial sampling rate (pixel size)

We imaged panCK (green) and CD11c (magenta) at lateral pixel sizes optimal for Nyquist sampling and denoising (60 nm) (**Supplementary Fig. 20a**) and then systematically reduced the sampling rate to 600 nm. We found that membrane midpoints could still be identified using polynomial curve fitting. We conclude that our spatial sampling rate (280 nm for most datasets) is sufficient for the distances we seek to measure.

#### Effect of magnification and numerical aperture in confocal imaging

We compared a high numerical aperture oil immersion lens (40x/1.3NA) to an air objective lens (20x/0.8NA). Since the sensitivity of an objective lens is dependent on numerical aperture and magnification, the 20x air objective lens is 3 times dimmer than the 40x/1.3NA. As a result, the overall signal and contrast is lower. Despite using a lower sampling rate (due to the lower numerical aperture), the membrane interactions were still detectable by curve fitting of line profiles. We surmise that this arises because out-of-focus light is still rejected by the confocal pinhole (**Supplementary Fig. 20b**), which was set to 1 Airy unit.

#### Effect of averaging over multiple line profiles

When the signal across a membrane is discontinuous or noisy, averaging several parallel line profiles across membranes can smoothen noisy data. This is more readily implementable across straight membranes than round membranes as in the case of cells from **Supplementary Fig. 22b** (right) imaged with a 40x/1.3NA objective lens with a 1 Airy Unit pinhole. However, depending on the severity of the noise, peaks are still identifiable via curve fitting of a single line profile (**Supplementary Fig. 21a**).

#### Effect of averaging over multiple Z-planes

We compared the use of a single line profile averaged across multiple Z-planes from the pair of cells in **Supplementary Fig. 20b** (right) imaged with a 40x/1.3NA objective lens. Unexpectedly, we found that precision was reduced when multiple Z-planes were used. We surmised that this arose because not all planes fully included the interaction and out-of-focus signals were increased. This can be observed by a widening of both membrane intensity curves (**Supplementary Fig. 21b**), which we observed beyond a thickness of 2.8 microns (10 planes spaced at 280 nm each). Despite this, curve fitting was still able to robustly identify peaks.

#### Effect of poor antibody staining resulting in low signal-to-noise ratio

We sought to assess the effect of lateral and axial resolution and signal-to-noise ratio (SNR) on the ability to study interacting membranes. Poor SNR can result from poor antibody staining.

**Supplementary Fig. 22a** demonstrates a PD1 and PDL1 interaction on two cells (PD1 conjugated to Alexafluor 647 - magenta; PDL1 conjugated to Alexafluor 647 in a different cycle - yellow). When anti-PD1 antibody was instead conjugated to phycoerythrin (green), a less photostable fluorophore, the overall signal intensity was significantly lower causing the line profile to be noisier. This is evidenced by the multiple peaks along the trace despite performing denoising operations. The peak-to-peak distance increased from 70 nm to 230 nm suggesting that the membranes were incorrectly identified as lying further apart. We conclude that analysis of membrane-membrane interaction requires “good” antibody staining; in the current manuscript we judged this visually, but in future work it should be possible to come develop a more objective metric.

#### Effect of out-of-focus rejection

The effect of low signal-to-background due to out-of-focus signal rejection was compared at 20x/0.8NA air and 40x/1.3NA oil immersion lens. For both magnifications, the out-of-focus signal from different optical planes masked the signal from the in-focus plane containing the cell membrane interactions (**Supplementary Fig. 22b**). This was evidenced by the high background signal level. Curve fitting was not able to find the correct peaks in cases where a tight interaction exists based on confocal images with a higher signal-to-background ratio. From this we conclude that membrane-membrane interaction requires 3D high-resolution data of the type that can be acquired by confocal but not conventional wide field microscopy.

In summary, we observed that contrast has the most significant impact on assessing cell membrane interactions. This can be influenced by methods to reduce background such as optical sectioning techniques to remove out-of-focus blur. Contrast is also dependent on the quality of antibody staining (bright fluorophores, specific antibodies) and sensitivity of the microscope detectors, which we did not directly compare. In the current work we opted to measure membrane intensity profiles along one or a small number of optical planes as this was observed to improve peak height as compared to using entire cell volumes. More specifically, we found that as the thickness of each optical plane decreases, the precision of detecting a membrane interaction increases (presumably being limited in super-resolution by the number of fluorophores in the field of view). However, through the use of curve fitting of membrane intensity profiles, the sampling rate and pixel size can be relatively coarse (in the interest of throughput) for membrane-membrane interaction analysis, but at the cost of less accurate characterization of cell morphology. Overall, we judge that the conditions used this paper represent a reasonable starting point for the development of automated proximity detection algorithms based on membrane contact rather than nuclear centroids.

## References

1. Kapałczyńska, M. et al. 2D and 3D cell cultures - a comparison of different types of cancer cell cultures. Arch Med Sci 14, 910–919 (2018).

2. Slaoui, M. & Fiette, L. Histopathology procedures: from tissue sampling to histopathological evaluation. Methods Mol Biol 691, 69–82 (2011).

3. Fischer, R. S., Wu, Y., Kanchanawong, P., Shroff, H. & Waterman, C. M. Microscopy in 3D: a biologist’s toolbox. Trends Cell Biol 21, 682–691 (2011).

4. Method of the Year 2024: spatial proteomics. Nat Methods 21, 2195–2196 (2024).

5. Radtke, A. J. et al. IBEX: an iterative immunolabeling and chemical bleaching method for high-content imaging of diverse tissues. Nat Protoc 17, 378–401 (2022).

6. Radtke, A. J. et al. IBEX: A versatile multiplex optical imaging approach for deep phenotyping and spatial analysis of cells in complex tissues. Proc. Natl. Acad. Sci. U.S.A. 117, 33455–33465 (2020).

7. Lin, J.-R. et al. Highly multiplexed immunofluorescence imaging of human tissues and tumors using t-CyCIF and conventional optical microscopes. eLife 7, e31657 (2018).

8. Bonnett, S. A. et al. Ultra High-plex Spatial Proteogenomic Investigation of Giant Cell Glioblastoma Multiforme Immune Infiltrates Reveals Distinct Protein and RNA Expression Profiles. Cancer Res Commun 3, 763–779 (2023).

9. Rao, A., Barkley, D., França, G. S. & Yanai, I. Exploring tissue architecture using spatial transcriptomics. Nature 596, 211–220 (2021).

10. Herman, B. & Lemasters, J. J. Optical Microscopy: Emerging Methods and Applications. (Elsevier, 2012).

11. Hickey, J. W. et al. Spatial mapping of protein composition and tissue organization: a primer for multiplexed antibody-based imaging. Nat Methods 19, 284–295 (2022).

12. Amin, M. B. et al. The Eighth Edition AJCC Cancer Staging Manual: Continuing to build a bridge from a population-based to a more ‘personalized’ approach to cancer staging. CA Cancer J Clin 67, 93–99 (2017).

13. Scudamore, C. L. A Practical Guide to the Histology of the Mouse. (John Wiley & Sons, 2014).

14. Matenaers, C., Popper, B., Rieger, A., Wanke, R. & Blutke, A. Practicable methods for histological section thickness measurement in quantitative stereological analyses. PLoS One 13, e0192879 (2018).

15. Masuda, S. et al. Tissue Thickness Interferes With the Estimation of the Immunohistochemical Intensity: Introduction of a Control System for Managing Tissue Thickness. Applied Immunohistochemistry & Molecular Morphology 29, 118 (2021).

16. Wu, X. & Hammer, J. A. ZEISS Airyscan: Optimizing usage for fast, gentle, super-resolution imaging. Methods Mol Biol 2304, 111–130 (2021).

17. Stelzer, E. H. K. et al. Light sheet fluorescence microscopy. Nat Rev Methods Primers 1, 1–25 (2021).

18. Creech, M. K., Wang, J., Nan, X. & Gibbs, S. L. Superresolution Imaging of Clinical Formalin Fixed Paraffin Embedded Breast Cancer with Single Molecule Localization Microscopy. Sci Rep 7, 40766 (2017).

19. Hoffer, J. et al. Minerva: a light-weight, narrative image browser for multiplexed tissue images. J Open Source Softw 5, 2579 (2020).

20. Shi, S.-R., Shi, Y. & Taylor, C. R. Antigen Retrieval Immunohistochemistry. J Histochem Cytochem 59, 13–32 (2011).

21. Zhou, F. Y. et al. A general algorithm for consensus 3D cell segmentation from 2D segmented stacks. bioRxiv 2024.05.03.592249 (2024) doi:10.1101/2024.05.03.592249.

22. Tran, T. et al. Correcting the Shrinkage Effects of Formalin Fixation and Tissue Processing for Renal Tumors: toward Standardization of Pathological Reporting of Tumor Size. J Cancer 6, 759–766 (2015).

23. Caicedo, J. C. et al. Nucleus segmentation across imaging experiments: the 2018 Data Science Bowl. Nat Methods 16, 1247–1253 (2019).

24. Ghose, S. et al. 3D reconstruction of skin and spatial mapping of immune cell density, vascular distance and effects of sun exposure and aging. Commun Biol 6, 718 (2023).

25. Peterson, D. A. Quantitative histology using confocal microscopy: implementation of unbiased stereology procedures. Methods 18, 493–507 (1999).

26. Schmitz, C. & Hof, P. R. Design-based stereology in neuroscience. Neuroscience 130, 813–831 (2005).

27. Wu, Y., Pegoraro, A. F., Weitz, D. A., Janmey, P. & Sun, S. X. The correlation between cell and nucleus size is explained by an eukaryotic cell growth model. PLoS Comput Biol 18, e1009400 (2022).

28. Benvenuti, F. et al. Requirement of Rac1 and Rac2 Expression by Mature Dendritic Cells for T Cell Priming. Science 305, 1150–1153 (2004).

29. Guilliams, M. et al. Dendritic cells, monocytes and macrophages: a unified nomenclature based on ontogeny. Nat Rev Immunol 14, 571–578 (2014).

30. Blauth, E., Kubitschke, H., Gottheil, P., Grosser, S. & Käs, J. A. Jamming in Embryogenesis and Cancer Progression. Frontiers in Physics 9, (2021).

31. Chen, H., Li, D. & Bar-Joseph, Z. SCS: cell segmentation for high-resolution spatial transcriptomics. Nat Methods 20, 1237–1243 (2023).

32. Muller, W. A. Getting Leukocytes to the Site of Inflammation. Vet Pathol 50, 7–22 (2013).

33. Van Goethem, E., Poincloux, R., Gauffre, F., Maridonneau-Parini, I. & Le Cabec, V. Matrix Architecture Dictates Three-Dimensional Migration Modes of Human Macrophages: Differential Involvement of Proteases and Podosome-Like Structures. The Journal of Immunology 184, 1049–1061 (2010).

34. Kuczek, D. E. et al. Collagen density regulates the activity of tumor-infiltrating T cells. j. immunotherapy cancer 7, 68 (2019).

35. Willsmore, Z. N. et al. B Cells in Patients With Melanoma: Implications for Treatment With Checkpoint Inhibitor Antibodies. Front Immunol 11, 622442 (2021).

36. Sehgal, P. B. et al. Murine GFP-Mx1 forms nuclear condensates and associates with cytoplasmic intermediate filaments: Novel antiviral activity against VSV. J Biol Chem 295, 18023–18035 (2021).

37. Woo, S.-R. et al. Differential subcellular localization of the regulatory T-cell protein LAG-3 and the coreceptor CD4. Eur J Immunol 40, 1768–1777 (2010).

38. Lin, L. et al. Granzyme B secretion by human memory CD4 T cells is less strictly regulated compared to memory CD8 T cells. BMC Immunol 15, 36 (2014).

39. Nirmal, A. J. et al. The Spatial Landscape of Progression and Immunoediting in Primary Melanoma at Single-Cell Resolution. Cancer Discovery 12, 1518–1541 (2022).

40. Jackett, L. A. & Scolyer, R. A. A Review of Key Biological and Molecular Events Underpinning Transformation of Melanocytes to Primary and Metastatic Melanoma. Cancers (Basel) 11, 2041 (2019).

41. Hodis, E. et al. A Landscape of Driver Mutations in Melanoma. Cell 150, 251–263 (2012).

42. Lian, C. G. et al. Loss of 5-hydroxymethylcytosine is an epigenetic hallmark of melanoma. Cell 150, 1135–1146 (2012).

43. Clark, W. H. et al. A study of tumor progression: The precursor lesions of superficial spreading and nodular melanoma. Human Pathology 15, 1147–1165 (1984).

44. Smoller, B. R. Histologic criteria for diagnosing primary cutaneous malignant melanoma. Mod Pathol 19, S34–S40 (2006).

45. Chudnovsky, Y., Khavari, P. A. & Adams, A. E. Melanoma genetics and the development of rational therapeutics. J Clin Invest 115, 813–824 (2005).

46. Kaufmann, C. et al. The role of cyclin D1 and Ki-67 in the development and prognostication of thin melanoma. Histopathology 77, 460–470 (2020).

47. Fetsch, P. A. et al. Melanoma-associated antigen recognized by T cells (MART-1): the advent of a preferred immunocytochemical antibody for the diagnosis of metastatic malignant melanoma with fine-needle aspiration. Cancer 87, 37–42 (1999).

48. Boiko, A. D. et al. Human melanoma-initiating cells express neural crest nerve growth factor receptor CD271. Nature 466, 133–137 (2010).

49. Oyler-Yaniv, A. et al. A Tunable Diffusion-Consumption Mechanism of Cytokine Propagation Enables Plasticity in Cell-to-Cell Communication in the Immune System. Immunity 46, 609–620 (2017).

50. Thibaut, R. et al. Bystander IFN-γ activity promotes widespread and sustained cytokine signaling altering the tumor microenvironment. Nat Cancer 1, 302–314 (2020).

51. Kim, Y. J. et al. Melanoma dedifferentiation induced by IFN-γ epigenetic remodeling in response to anti-PD-1 therapy. J Clin Invest 131, e145859, 145859 (2021).

52. Watanabe, R. et al. Human skin is protected by four functionally and phenotypically discrete populations of resident and recirculating memory T cells. Sci Transl Med 7, 279ra39 (2015).

53. Utzschneider, D. T. et al. Early precursor T cells establish and propagate T cell exhaustion in chronic infection. Nat Immunol 21, 1256–1266 (2020).

54. Burger, M. L. et al. Antigen dominance hierarchies shape TCF1+ progenitor CD8 T cell phenotypes in tumors. Cell 184, 4996–5014.e26 (2021).

55. Miller, B. C. et al. Subsets of exhausted CD8+ T cells differentially mediate tumor control and respond to checkpoint blockade. Nat Immunol 20, 326–336 (2019).

56. Kallies, A., Zehn, D. & Utzschneider, D. T. Precursor exhausted T cells: key to successful immunotherapy? Nat Rev Immunol 20, 128–136 (2020).

57. Escobar, G., Mangani, D. & Anderson, A. C. T cell factor 1 (Tcf1): a master regulator of the T cell response in disease. Sci Immunol 5, eabb9726 (2020).

58. Zhou, P. et al. Single-cell CRISPR screens in vivo map T cell fate regulomes in cancer. Nature 624, 154–163 (2023).

59. Rahim, M. K. et al. Dynamic CD8+ T cell responses to cancer immunotherapy in human regional lymph nodes are disrupted in metastatic lymph nodes. Cell 186, 1127–1143.e18 (2023).

60. Dustin, M. L. The immunological synapse. Cancer Immunol Res 2, 1023–1033 (2014).

61. Grosser, S. et al. Cell and Nucleus Shape as an Indicator of Tissue Fluidity in Carcinoma. Phys. Rev. X 11, 011033 (2021).

62. Gaglia, G. et al. Lymphocyte networks are dynamic cellular communities in the immunoregulatory landscape of lung adenocarcinoma. Cancer Cell 41, 871–886.e10 (2023).

63. Cabrita, R. et al. Tertiary lymphoid structures improve immunotherapy and survival in melanoma. Nature 577, 561–565 (2020).

64. Mörth, E., et al. A Mixed Reality and 2D Display Hybrid Approach for Visual Analysis of 3D Tissue Maps. Preprint at 10.31219/osf.io/zka2j (2024).

65. Arroyo, A. et al. Ce3D-IBEX: Achieving Multiplex 3-dimensional Imaging for Deep Phenotyping of Cells in Tissues. The Journal of Immunology 212, 1506_4978 (2024).

66. Huang, F., Santinon, F., Flores González, R. E. & del Rincón, S. V. Melanoma Plasticity: Promoter of Metastasis and Resistance to Therapy. Front Oncol 11, 756001 (2021).

67. Knutsdottir, H., Condeelis, J. S. & Palsson, E. 3-D individual cell based computational modeling of tumor cell–macrophage paracrine signaling mediated by EGF and CSF-1 gradients. Integr Biol (Camb) 8, 104–119 (2016).

68. Zangooei, M. H., Margolis, R. & Hoyt, K. Multiscale computational modeling of cancer growth using features derived from microCT images. Sci Rep 11, 18524 (2021).

69. Li, W., Germain, R. N. & Gerner, M. Y. Multiplex, quantitative cellular analysis in large tissue volumes with clearing-enhanced 3D microscopy (Ce3D). Proc Natl Acad Sci U S A 114, E7321–E7330 (2017).

70. Lin, J.-R. et al. Multiplexed 3D atlas of state transitions and immune interaction in colorectal cancer. Cell 186, 363–381.e19 (2023).

71. Keller, M. S. et al. Vitessce: integrative visualization of multimodal and spatially resolved single-cell data. Nat Methods (2024) doi:10.1038/s41592-024-02436-x.

72. Kuett, L. et al. Three-dimensional imaging mass cytometry for highly multiplexed molecular and cellular mapping of tissues and the tumor microenvironment. Nat Cancer 3, 122–133 (2022).

73. Wallace, W., Schaefer, L. H. & Swedlow, J. R. A Workingperson’s Guide to Deconvolution in Light Microscopy. BioTechniques 31, 1076–1097 (2001).

74. Fish, K. N. Total Internal Reflection Fluorescence (TIRF) Microscopy. CP Cytometry 50, (2009).

75. Axelrod, D., Thompson, N. L. & Burghardt, T. P. Total internal reflection fluorescent microscopy. Journal of Microscopy 129, 19–28 (1983).

76. Egger, M. D. & Petráň, M. New Reflected-Light Microscope for Viewing Unstained Brain and Ganglion Cells. Science 157, 305–307 (1967).

77. Wilson, T. Resolution and optical sectioning in the confocal microscope: Journal of Microscopy 244, 113–121 (2011).

78. Wilson, T. Optical sectioning in confocal fluorescent microscopes. Journal of Microscopy 154, 143–156 (1989).

79. Denk, W., Strickler, J. H. & Webb, W. W. Two-Photon Laser Scanning Fluorescence Microscopy. Science 248, 73–76 (1990).

80. Campagnola, P. J. et al. Three-Dimensional High-Resolution Second-Harmonic Generation Imaging of Endogenous Structural Proteins in Biological Tissues. Biophysical Journal 82, 493–508 (2002).

81. Campagnola, P. J., Wei, M., Lewis, A. & Loew, L. M. High-Resolution Nonlinear Optical Imaging of Live Cells by Second Harmonic Generation. Biophysical Journal 77, 3341–3349 (1999).

82. Zipfel, W. R. et al. Live tissue intrinsic emission microscopy using multiphoton-excited native fluorescence and second harmonic generation. Proc. Natl. Acad. Sci. U.S.A. 100, 7075–7080 (2003).

83. Bakker, G.-J. et al. Intravital deep-tumor single-beam 3-photon, 4-photon, and harmonic microscopy. Elife 11, e63776 (2022).

84. Murray, J. M., Appleton, P. L., Swedlow, J. R. & Waters, J. C. Evaluating performance in three-dimensional fluorescence microscopy. J Microsc 228, 390–405 (2007).

85. Renier, N. et al. iDISCO: A Simple, Rapid Method to Immunolabel Large Tissue Samples for Volume Imaging. Cell 159, 896–910 (2014).

86. Tomer, R., Ye, L., Hsueh, B. & Deisseroth, K. Advanced CLARITY for rapid and high-resolution imaging of intact tissues. Nat Protoc 9, 1682–1697 (2014).

87. Tanaka, N. et al. Whole-tissue biopsy phenotyping of three-dimensional tumours reveals patterns of cancer heterogeneity. Nat Biomed Eng 1, 796–806 (2017).

88. Li, W., Germain, R. N. & Gerner, M. Y. High-dimensional cell-level analysis of tissues with Ce3D multiplex volume imaging. Nat Protoc 14, 1708–1733 (2019).

89. Murray, E. et al. Simple, Scalable Proteomic Imaging for High-Dimensional Profiling of Intact Systems. Cell 163, 1500–1514 (2015).

90. Chen, B.-C. et al. Lattice light-sheet microscopy: Imaging molecules to embryos at high spatiotemporal resolution. Science 346, 1257998 (2014).

91. Dean, K. M., Roudot, P., Welf, E. S., Danuser, G. & Fiolka, R. Deconvolution-free Subcellular Imaging with Axially Swept Light Sheet Microscopy. Biophysical Journal 108, 2807–2815 (2015).

92. Ku, T. et al. Multiplexed and scalable super-resolution imaging of three-dimensional protein localization in size-adjustable tissues. Nat Biotechnol 34, 973–981 (2016).

93. Park, J. et al. Epitope-preserving magnified analysis of proteome (eMAP). Sci Adv 7, eabf6589 (2021).

94. Saka, S. K. et al. Immuno-SABER enables highly multiplexed and amplified protein imaging in tissues. Nat Biotechnol 37, 1080–1090 (2019).

95. Scholaert, M. et al. 3D deconvolution of human skin immune architecture with Multiplex Annotated Tissue Imaging System. Sci Adv 9, eadf9491 (2023).

96. Van Ineveld, R. L. et al. Revealing the spatio-phenotypic patterning of cells in healthy and tumor tissues with mLSR-3D and STAPL-3D. Nat Biotechnol 39, 1239–1245 (2021).

97. Shi, L., Wei, M. & Min, W. Highly-Multiplexed Tissue Imaging with Raman Dyes. JoVE 63547 (2022) doi:10.3791/63547.

98. Stringer, C., Wang, T., Michaelos, M. & Pachitariu, M. Cellpose: a generalist algorithm for cellular segmentation. Nat Methods 18, 100–106 (2021).

99. Zhou, D., Bousquet, O., Lal, T., Weston, J. & Schölkopf, B. Learning with Local and Global Consistency. in Advances in Neural Information Processing Systems (eds. Thrun, S., Saul, L. & Schölkopf, B.) vol. 16 (MIT Press, 2003).

100. Niederhuber, M. J., Lambert, T. J., Yapp, C., Silver, P. A. & Polka, J. K. Superresolution microscopy of the β-carboxysome reveals a homogeneous matrix. Mol Biol Cell 28, 2734–2745 (2017).

