## Supplementary Figures for "Highly Multiplexed 3D Profiling of Cell States and Immune Niches in Human Tumours"

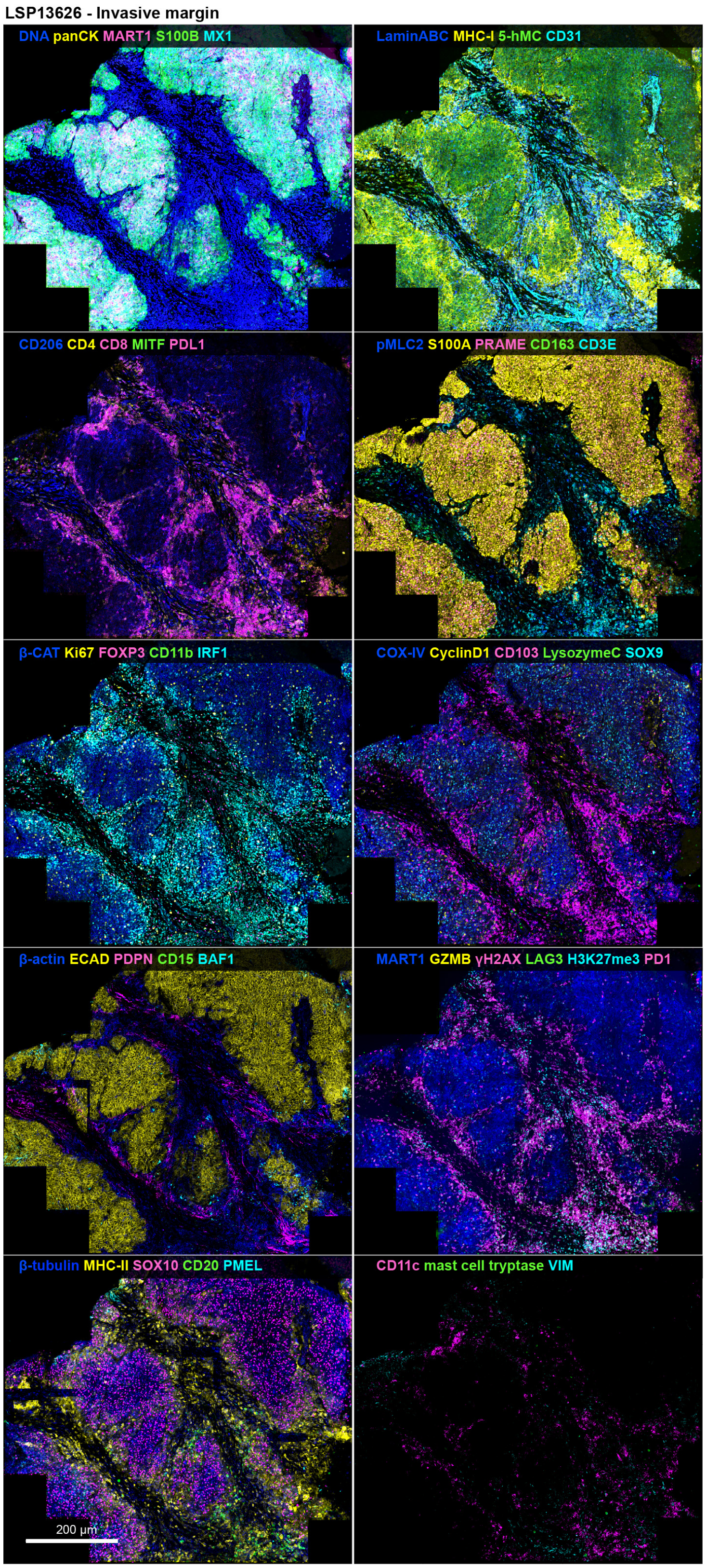


**Supplementary Figure 1. Z-projection of full dataset for invasive melanoma (vertical growth phase; VGP) region for tissue section LSP13626.** Imaged with 40x/1.3NA oil immersion objective lens on Zeiss LSM980 confocal microscope; sampled at 140nm (x,y) and 280 (z) over a 35-micron thick tissue specimen. See **Supplementary Table 1** for patient info and Minerva story. See **Supplementary Table 3** for marker panel.

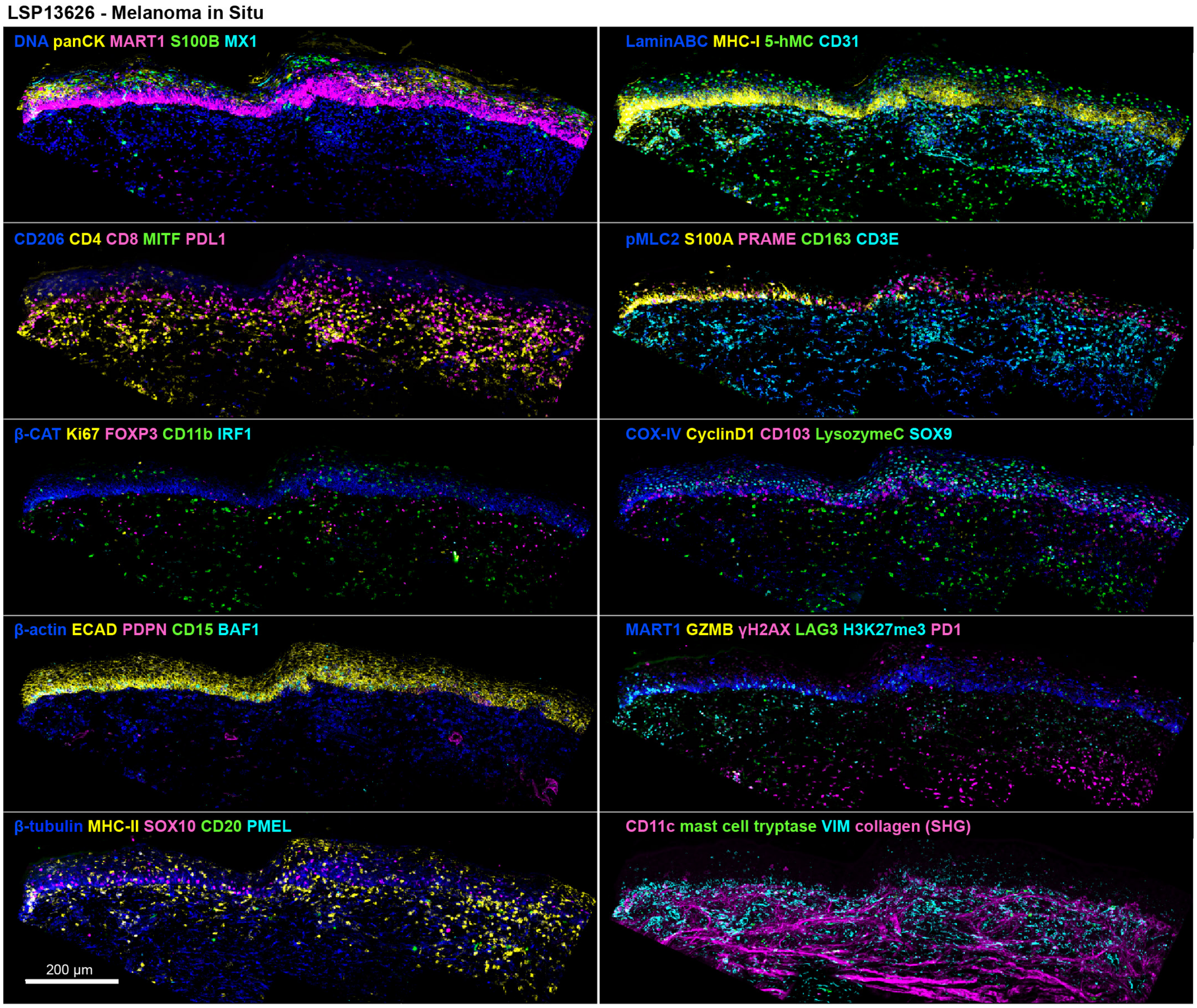


**Supplementary Figure 2. Z-projection of full dataset for melanoma in situ (MIS) region for tissue section LSP13626.** Imaged with 40x/1.3NA oil immersion objective lens on Zeiss LSM980 confocal microscope; sampled at 140nm (x,y) and 280 (z) over a 35-micron thick tissue specimen. See **Supplementary Table 1** for patient info and Minerva story. See **Supplementary Table 3** for marker panel.

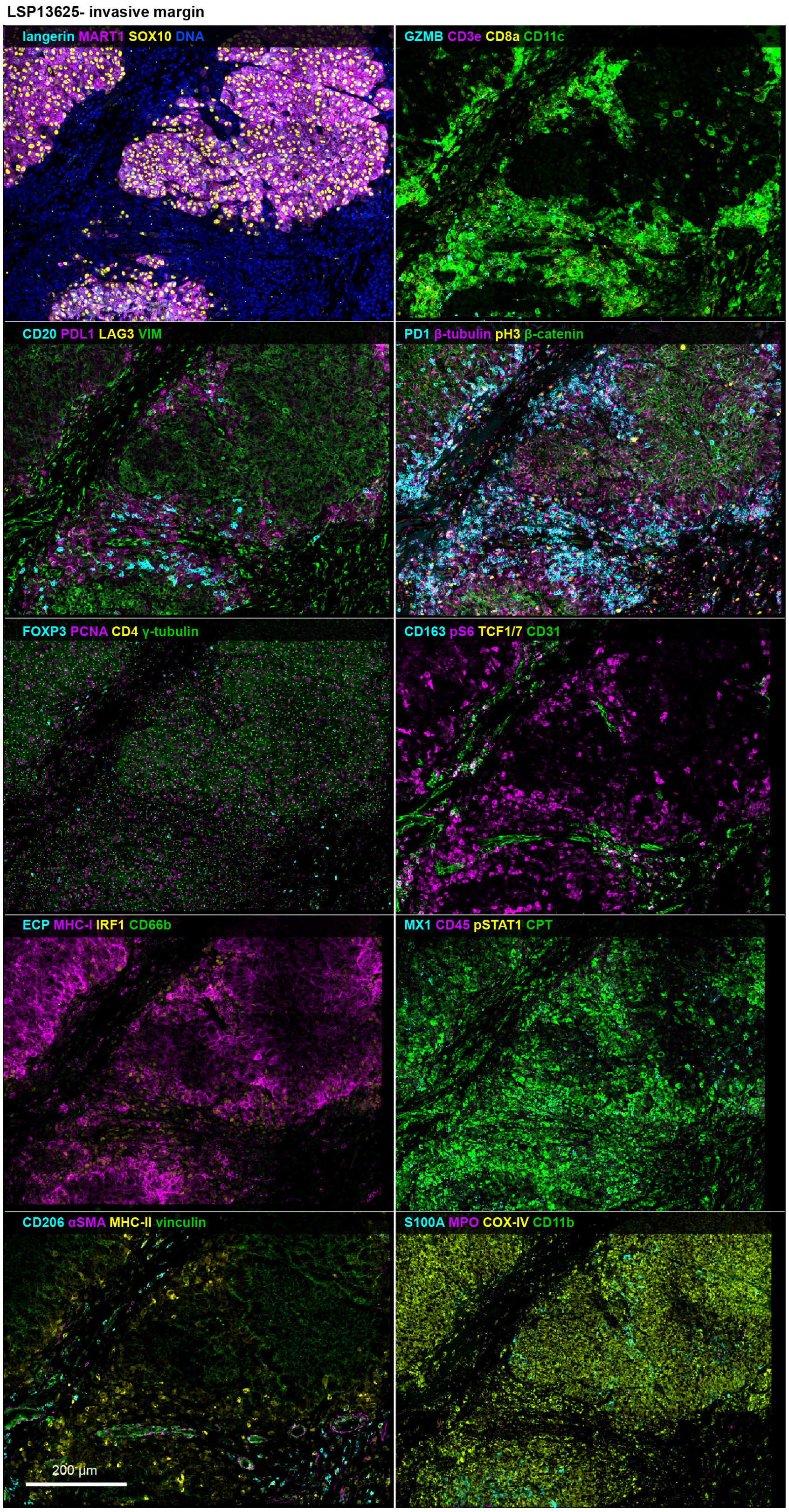


**Supplementary Figure 3.** Z-projection of full dataset for invasive melanoma (vertical growth phase; VGP) region for tissue section LSP13625. Imaged with 40x/1.3NA oil immersion objective lens on Zeiss LSM980 confocal microscope; sampled at 140nm (x,y) and 280 (z) over a 35-micron thick tissue specimen. See **Supplementary Table 1** for patient info and Minerva story. See **Supplementary Table 4** for marker panel.

LSP13625- Melanoma in Situ

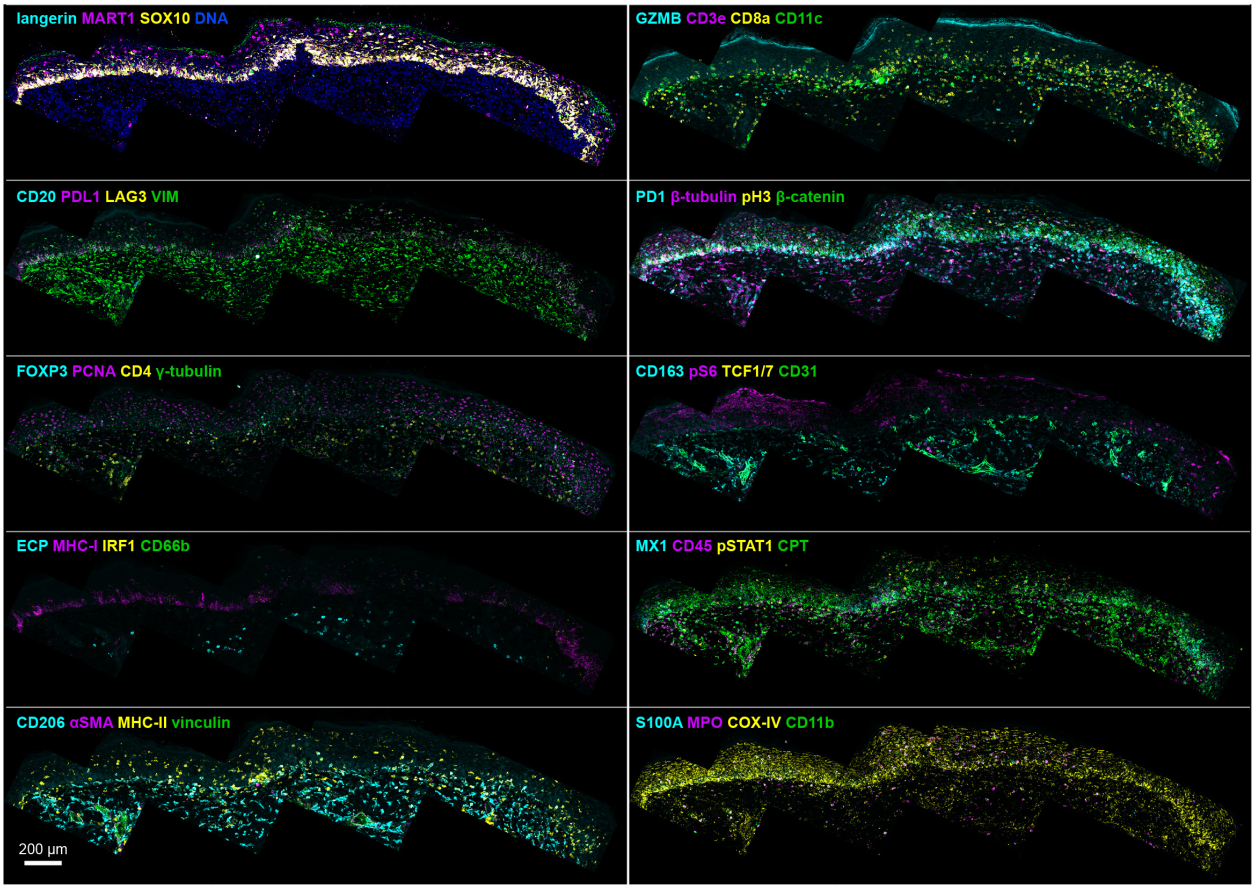


**Supplementary Figure 4. Z-projection of full dataset for melanoma in situ (MIS) region for tissue section LSP13625.** Imaged with 40x/1.3NA oil immersion objective lens on Zeiss LSM980 confocal microscope; sampled at 140nm (x,y) and 280 (z) over a 35-micron thick tissue specimen. See **Supplementary Table 1** for patient info and Minerva story. See **Supplementary Table 4** for marker panel.

LSP22409 - metastatic melanoma

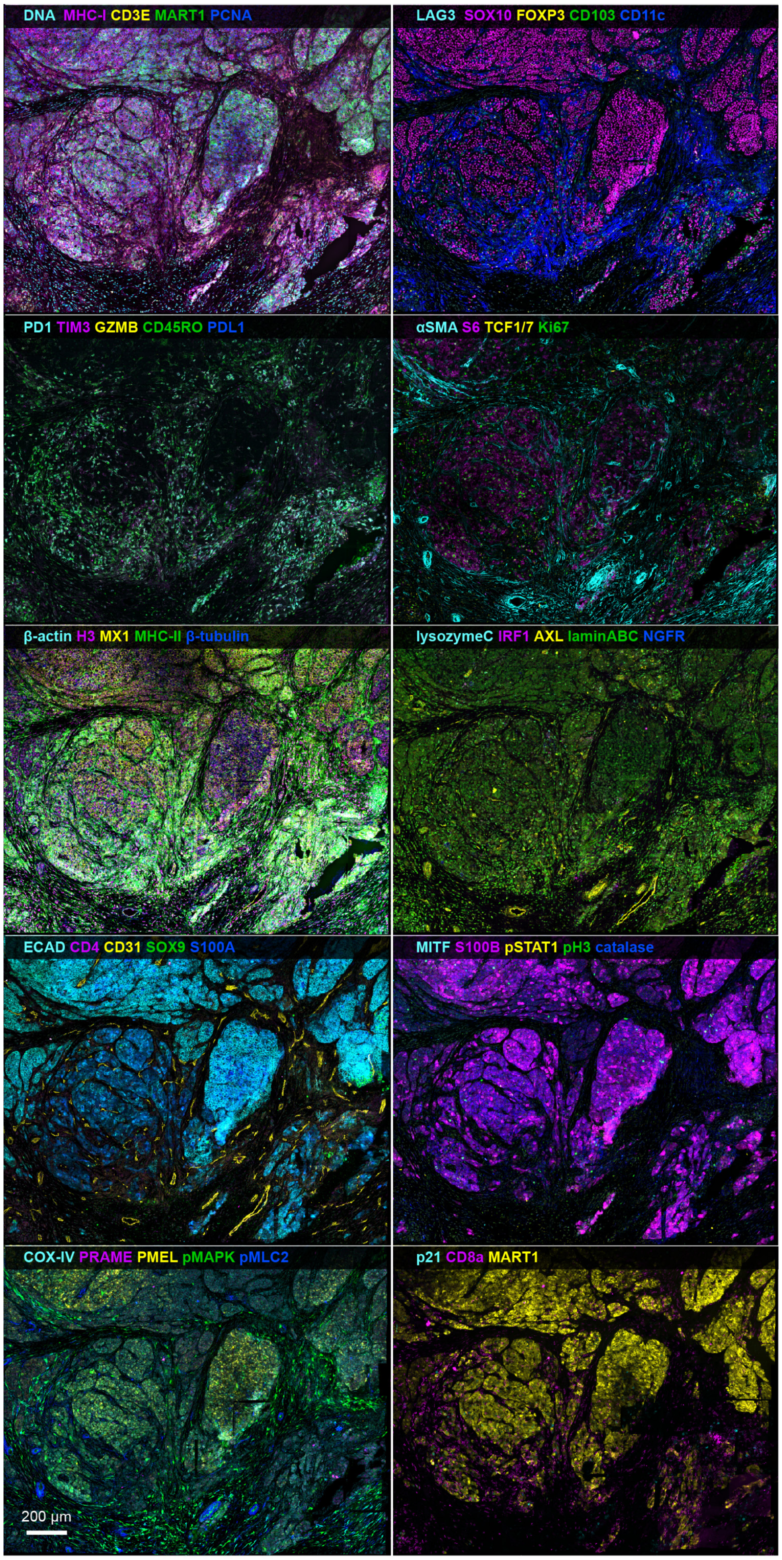


**Supplementary Figure 5. Z-projection of full dataset for metastatic melanoma (tissue section LSP22409).** Imaged with 40x/1.3NA oil immersion objective lens on Zeiss LSM980 confocal microscope; sampled at 140nm (x,y) and 280 (z) over a 25-micron thick tissue specimen. See **Supplementary Table 1** for patient info and Minerva story. See **Supplementary Table 5** for marker panel.

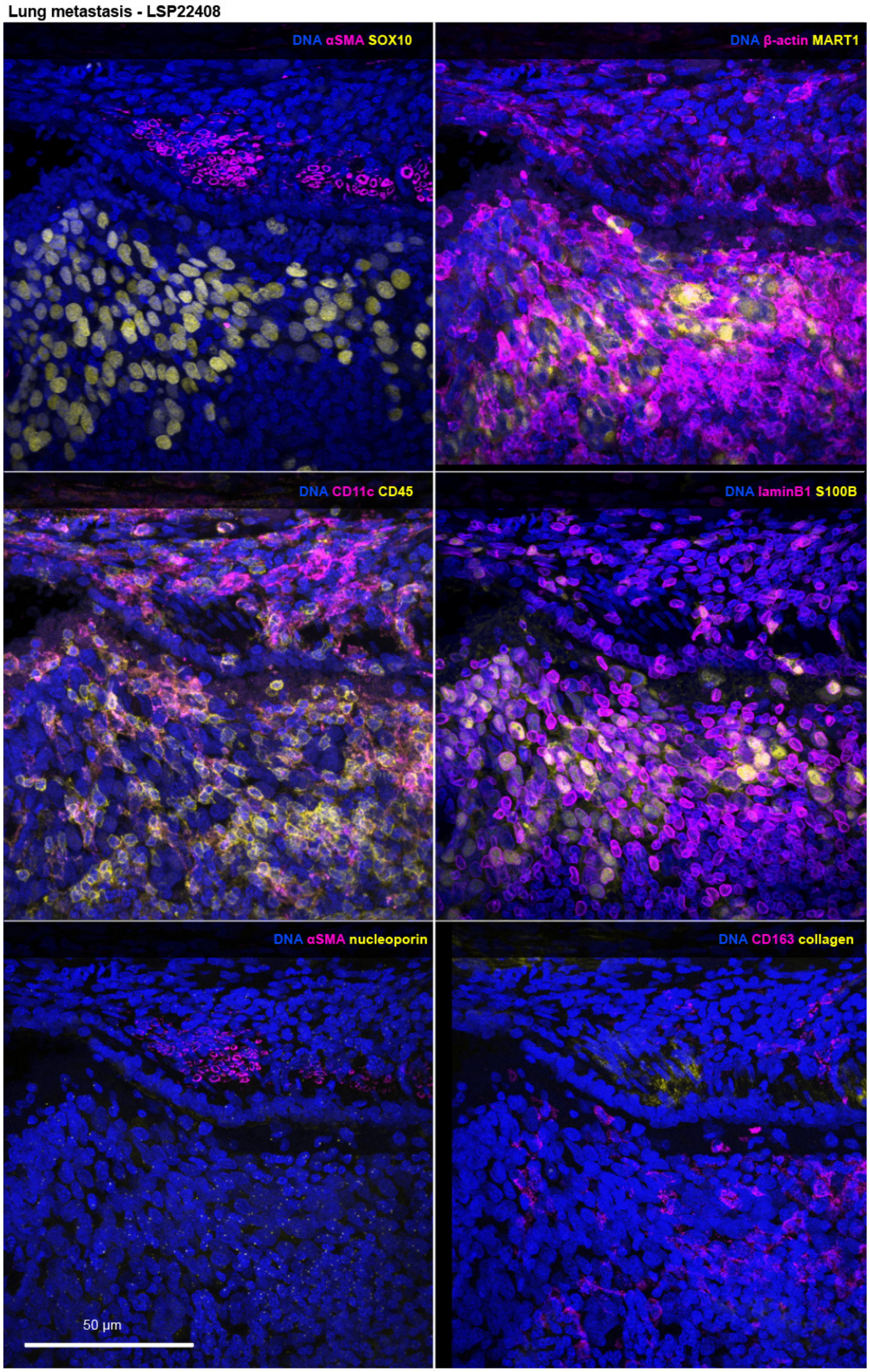


**Supplementary Figure 6. Z-projection of full dataset for lung metastasis (tissue section LSP22408).** Imaged with 40x/1.25NA silicone oil objective lens on Olympus FV1200 confocal microscope; sampled at 310nm (x,y) and 540 (z) over a 40-micron thick tissue specimen. See **Supplementary Table 1** for patient info and Minerva story. See **Supplementary Table 6** for marker panel.

**Glioblastoma - LSP17378**

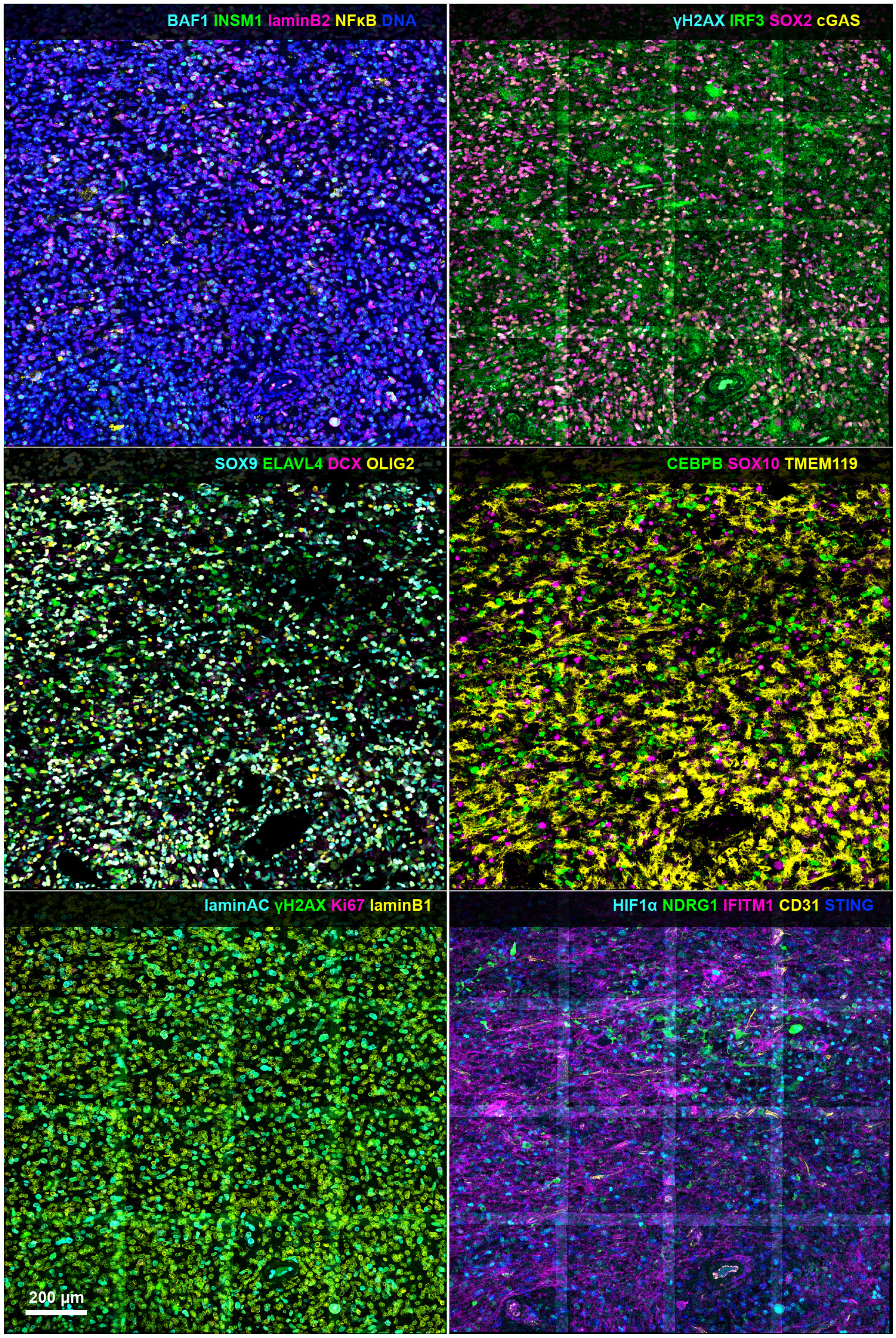


**Supplementary Figure 7. Z-projection of full dataset for glioblastoma (tissue section LSP17378).** Imaged with 40x/1.3NA objective lens on Zeiss LSM980 confocal microscope; sampled at 140nm (x,y) and 280 (z) over a 20-micron thick tissue specimen. See **Supplementary Table 1** for patient info and Minerva story. See **Supplementary Table 7** for marker panel.

Serous Tubal Intraepithelial Carcinoma (STIC) - region TR3 - LSP18251

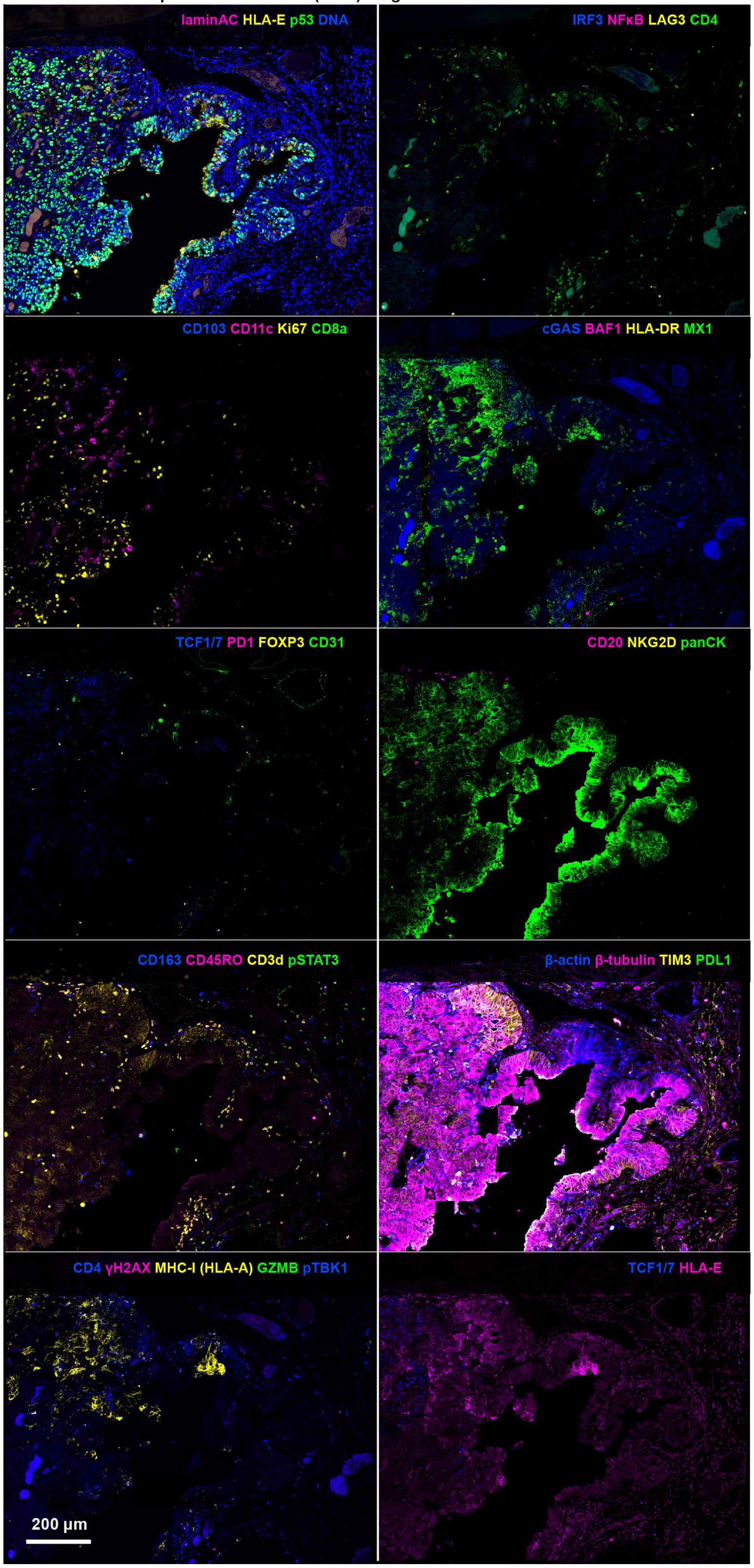


**Supplementary Figure 8. Z-projection of full dataset for serous tubal intraepithelial carcinoma (STIC), region TR3 (tissue section LSP18251).** Imaged with 40x/1.3NA oil immersion objective lens on Zeiss LSM980 confocal microscope; sampled at 140nm (x,y) and 280 (z) over a 20-micron thick tissue specimen. See **Supplementary Table 1** for patient info and Minerva story. See **Supplementary Table 8** for marker panel.

Serous Tubal Intraepithelial Carcinoma (STIC) - region TR4 - LSP18251

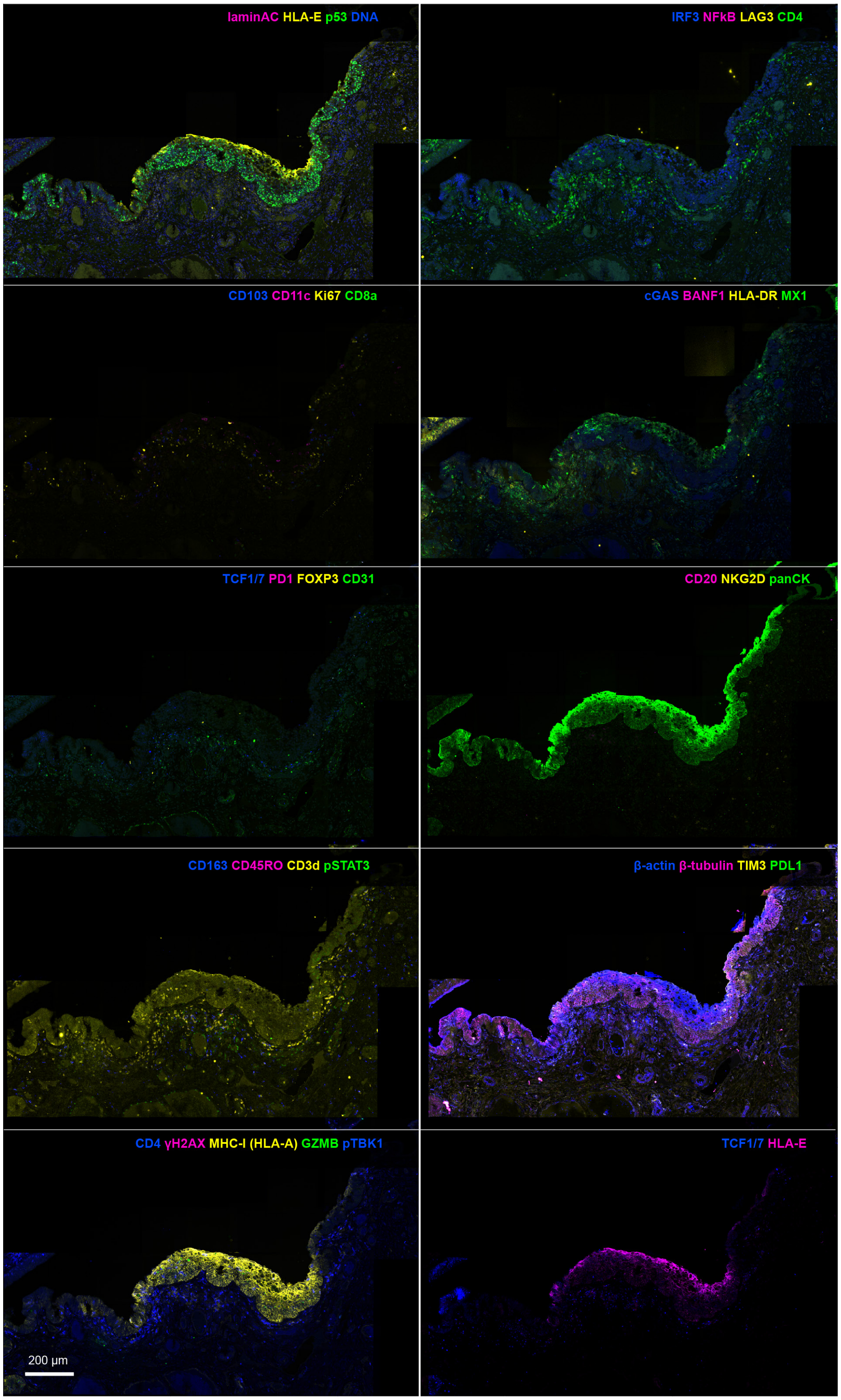


**Supplementary Figure 9. Z-projection of full dataset for serous tubal intraepithelial carcinoma (STIC), region TR4 (tissue section LSP18251).** Imaged with 40x/1.3NA oil immersion objective lens on Zeiss LSM980 confocal microscope; sampled at 140nm (x,y) and 280 (z) over a 20-micron thick tissue specimen. See **Supplementary Table 1** for patient info and Minerva story. See **Supplementary Table 8** for marker panel.

Serous Tubal Intraepithelial Carcinoma (STIC) - region TR5 - LSP18251

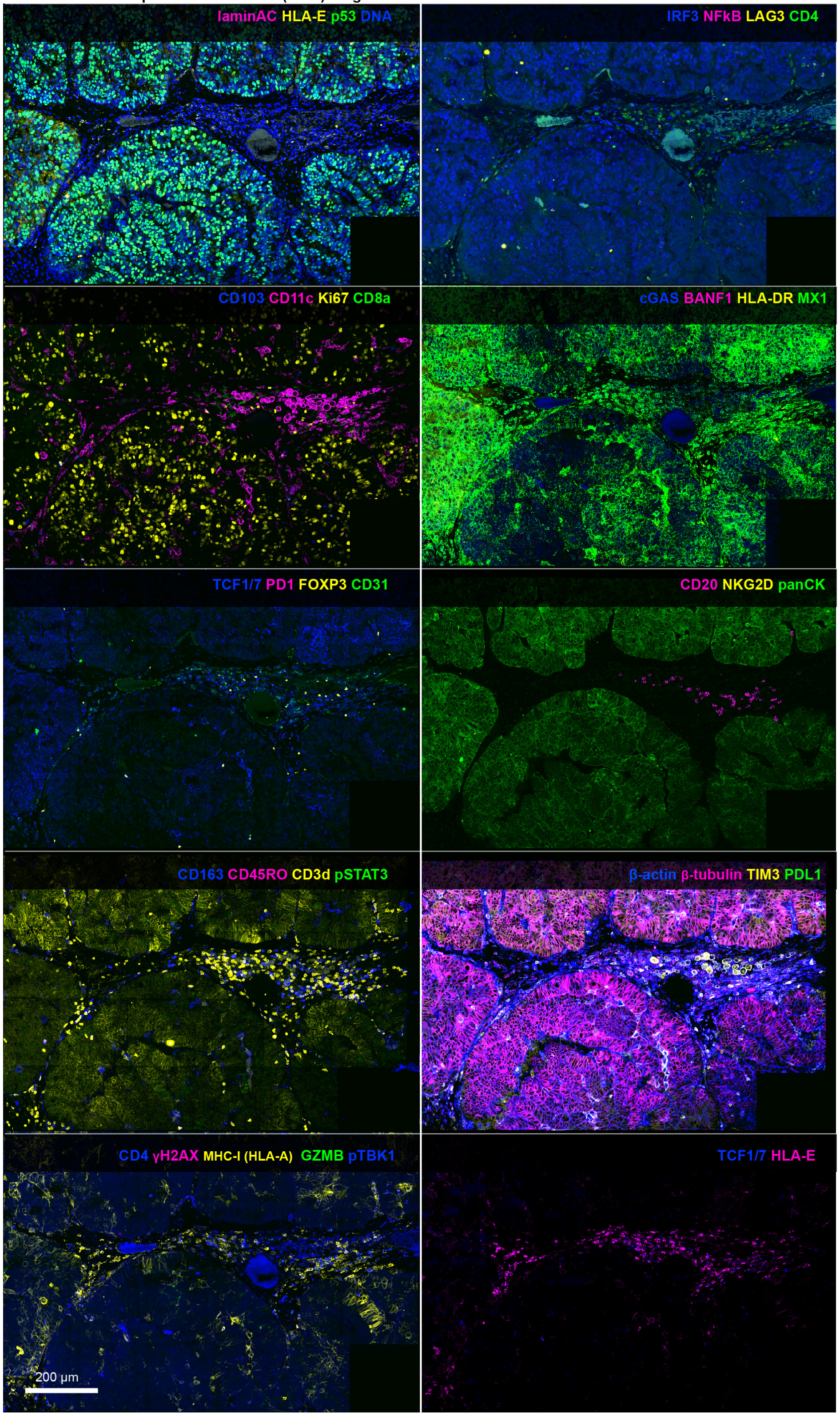


**Supplementary Figure 10. Z-projection of full dataset for serous tubal intraepithelial carcinoma (STIC), region TR5 (tissue section LSP18251).** Imaged with 40x/1.3NA oil immersion objective lens on Zeiss LSM980 confocal microscope; sampled at 140nm (x,y) and 280 (z) over a 20-micron thick tissue specimen. See **Supplementary Table 1** for patient info and Minerva story. See **Supplementary Table 8** for marker panel.

Tonsil - LSP13357

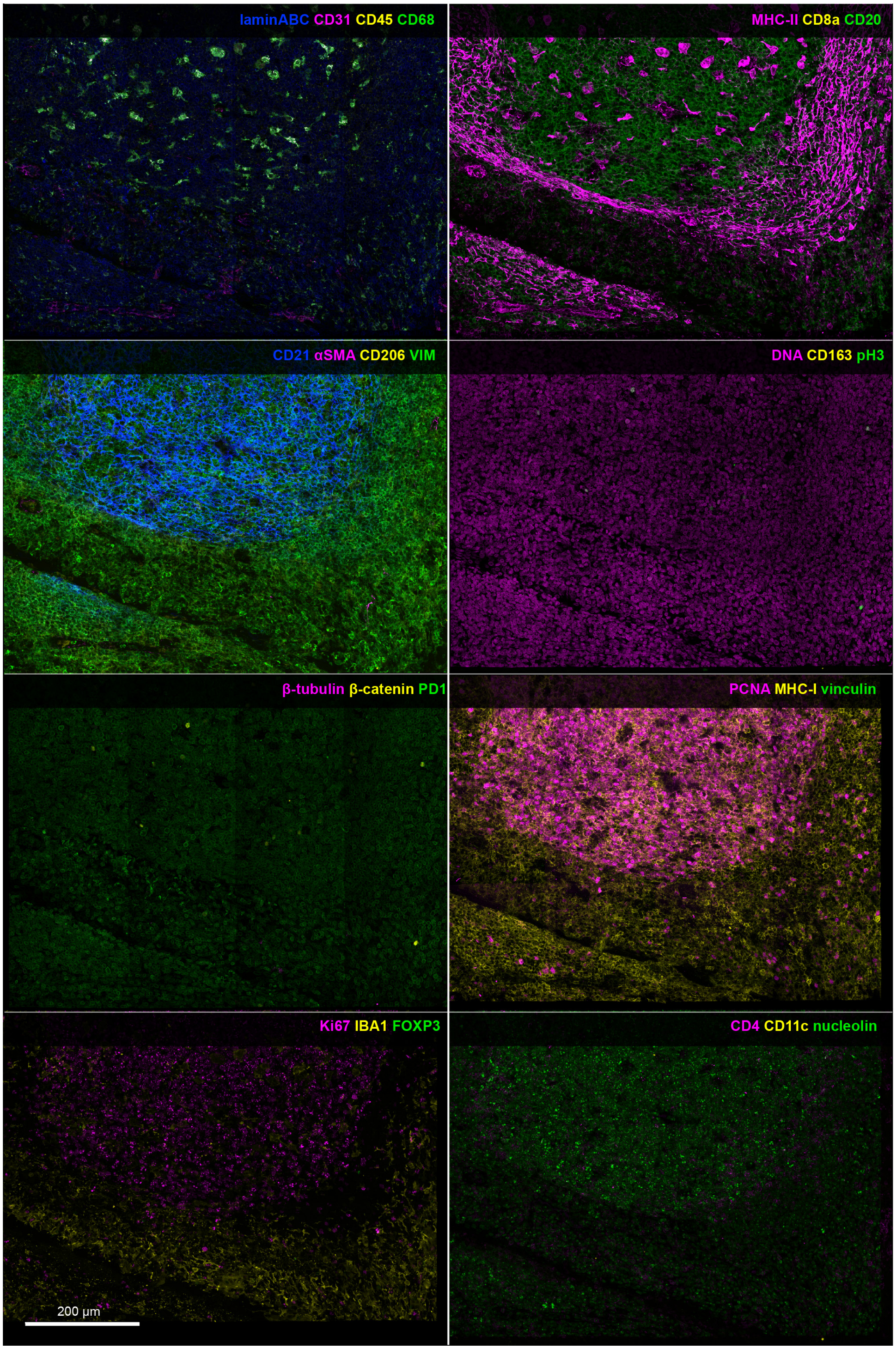


**Supplementary Figure 11. Z-projection of full dataset of tonsil (tissue section LSP13357).** Imaged with 40x/1.2NA water immersion objective lens on Zeiss LSM980 confocal microscope; sampled at 140nm (x,y) and 280 (z) over a 20-micron thick tissue specimen. See **Supplementary Table 1** for patient info and Minerva story. See **Supplementary Table 9** for marker panel.

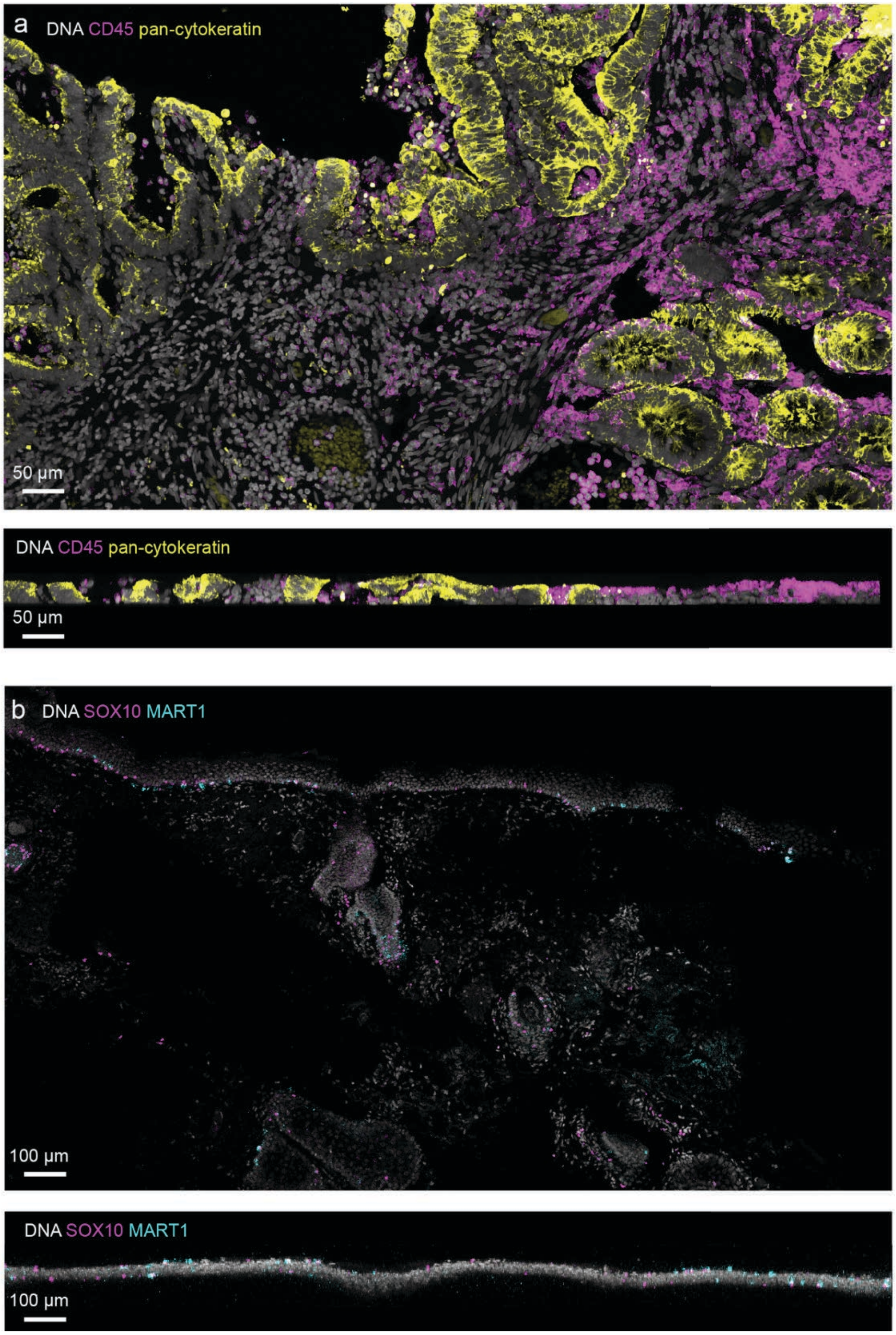


**Supplementary Figure 12. Tissue processing strategies to improve tissue adherence for fragile samples. a,** XY (top) and XZ (bottom) projection views of a 35-micron thick colorectal cancer sample that survived more than 6 cycles of CyCIF with the set up from **Extended Data Figure 1b** and with Matrigel coated over the sample. XZ projection shows that the tissue tolerated multiple rounds of bleaching and adhered to the coverslip along the length of the tissue. **b,** XY (top) and XZ (bottom) projection views of a fragile melanoma precursor sample (LSP27564 – see **Supplementary Table 10** for marker panel) with the set up from **Extended Data Figure 1b** but with a mesh over the sample. Bottom XZ projection shows that the tissue exhibited significantly poorer adhesion to the coverslip shown by the wavy pattern. Yet the tissue survived 3 cycles of CyCIF as a result of overlaying a protective mesh over the tissue. In contrast, setup from **Extended Data Figure 1a** caused the sample to immediately detach after initial bleaching.

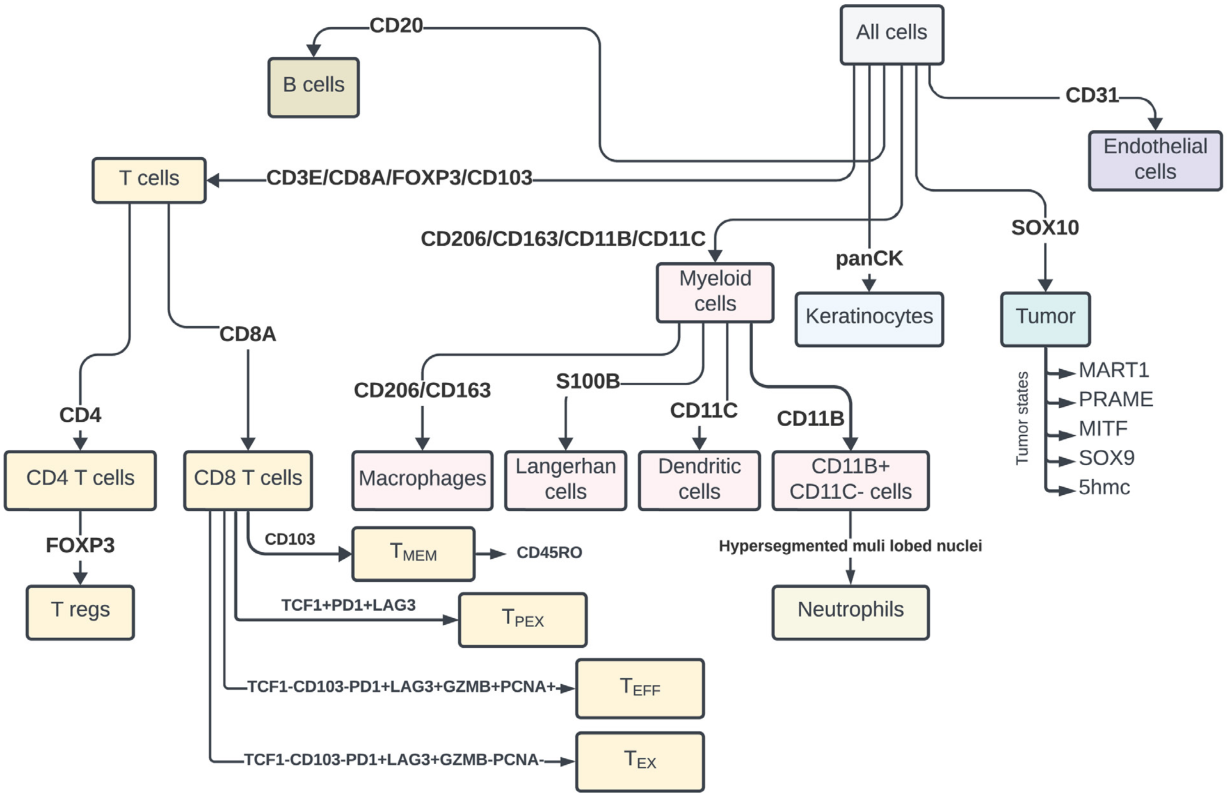


**Additional markers that were used to determine cell states and morphologies:**

| | | | | |
|---|---|---|---|---|
| COX-IV | CyclinD1 | MX1 | pMLC2 | β-actin |
| γH2AX | KI67 | IRF1 | VIM | β-catenin |
| | PCNA | pSTAT1 | Vinculin | β-tubulin |

**Supplementary Figure 13. Flowchart used for cell type calling in melanoma.** Lines and arrows represent cells that stain positive for the indicated markers. Under tumour cells, specific tumour cell states are shown. Cell states that are not related to a specific marker are shown in the lower box.

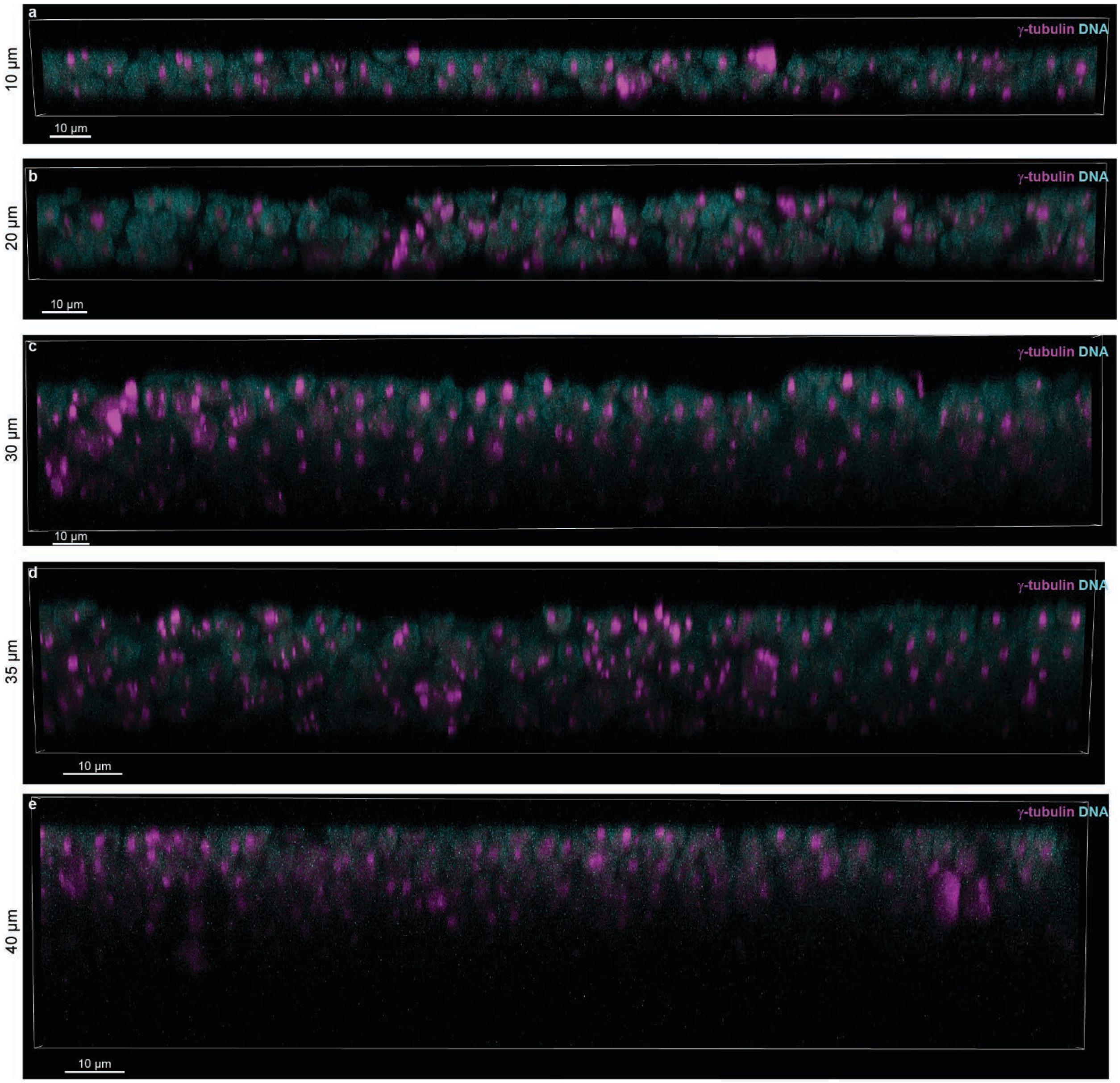


**Supplementary Figure 14. Antibody penetration in tissues of different thicknesses. a-e,** Orthogonal views of γ-tubulin-Alexafluor 555 staining (magenta) tonsil sections of 10 µm (**a**), 20 µm (**b**), 30 µm (**c**), 35 µm (**d**), and 40 µm (**e**) thicknesses, respectively. γ-tubulin staining can be observed from the top to bottom of the tissue up until 35-micron thick tissue sections. In the 40-micron thick sample, γ-tubulin appears dimmer and significantly distorted at the far side of the sample due to lack of antibody penetration or light attenuation. All tonsil section prepared on glass slides, stained simultaneously at 4 degrees overnight, washed in PBS, and mounted with 70% glycerol. Imaged with 40x/1.2NA water objective lens on Zeiss LSM980 laser scanning confocal microscope sampled at 104nm (x,y) and 230nm (z). DNA, stained with Hoechst, shown in cyan. Scalebar 10 µm.

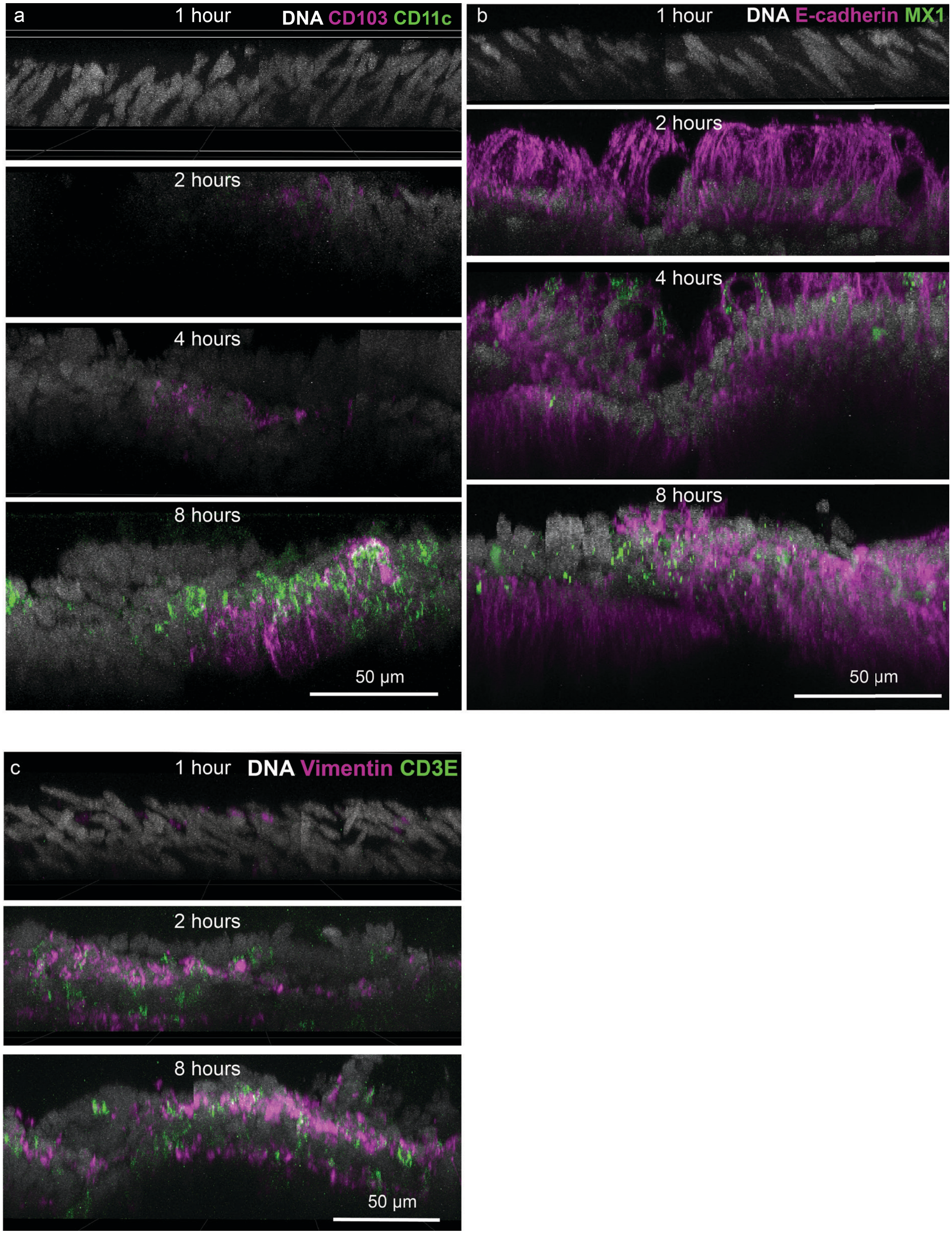


**Supplementary Figure 15: Timelapse of antibody penetration. a-c,** Orthogonal views of different 35 µm thick colorectal cancer tissue of various groups immune, tumour, and stromal markers imaged after 1, 2, 4 or 8 hours of staining at room temperature and washed in PBS. Staining was not apparent until after 2 hours of incubation. Full antibody penetration was observed after 8 hours of staining at room temperature. Tissue specimens prepared and stained on glass coverslips with vimentin-Alexafluor 750 (magenta), CD3e-Alexafluor 555 (green), Hoechst for DNA (grey) and mounted with 70% glycerol. Imaged with 40x/1.3NA oil objective lens on Zeiss LSM980 laser scanning confocal microscope sampled at 414nm (x,y) and 290nm (z).

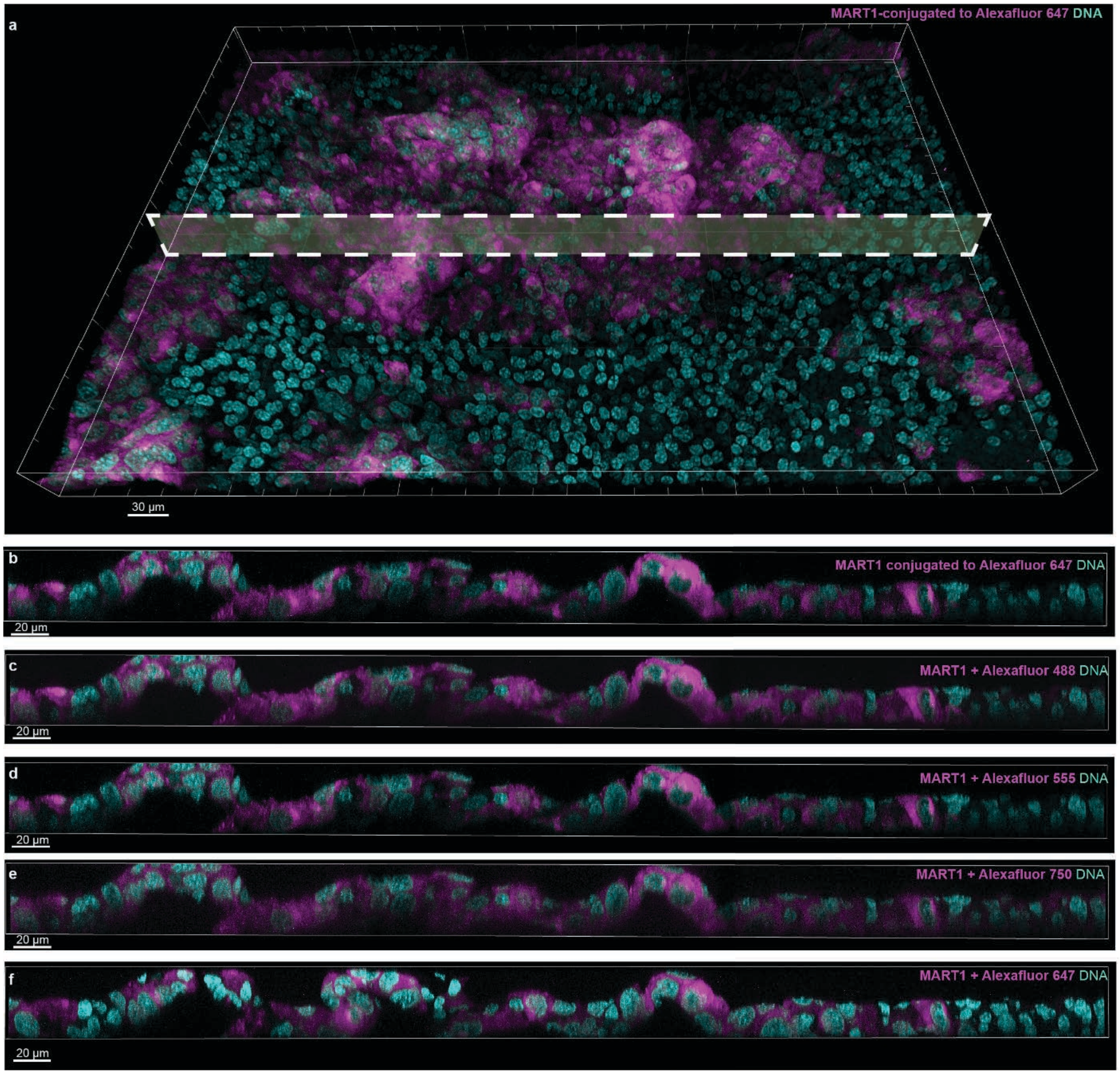


**Supplementary Figure 16. Comparison of antibody penetration with different secondary antibodies against a MART1 primary conjugate. a,** Volume rendering of primary melanoma, with dashed white rectangle indicating the location of the orthogonal views in **b-f**. Scale bar 30 µm. **b-f,** MART1-conjugated to Alexafluor 647 (**b**) Alexafluor 488 (**c**), Alexafluor 555 (**d**), Alexafluor 750 (**e**), or Alexafluor 647 (**f**). All combinations except MART1 + Alexafluor 647 were stained in the same cycle. MART1 + Alexafluor 647 was stained on the next cycle at the same location and same tissue specimen. Results indicate that, for MART1, fluorophore did not affect staining penetration. The use of primary conjugated vs unconjugated also did not influence staining pattern. Tissue section prepared and stained on glass coverslip at 4 degrees C overnight, washed in PBS, and mounted with 70% glycerol. Imaged with 40x/1.3NA oil objective lens on Zeiss LSM980 laser scanning confocal microscope sampled at 207nm (x,y) and 280nm (z). Scale bars 20 µm.

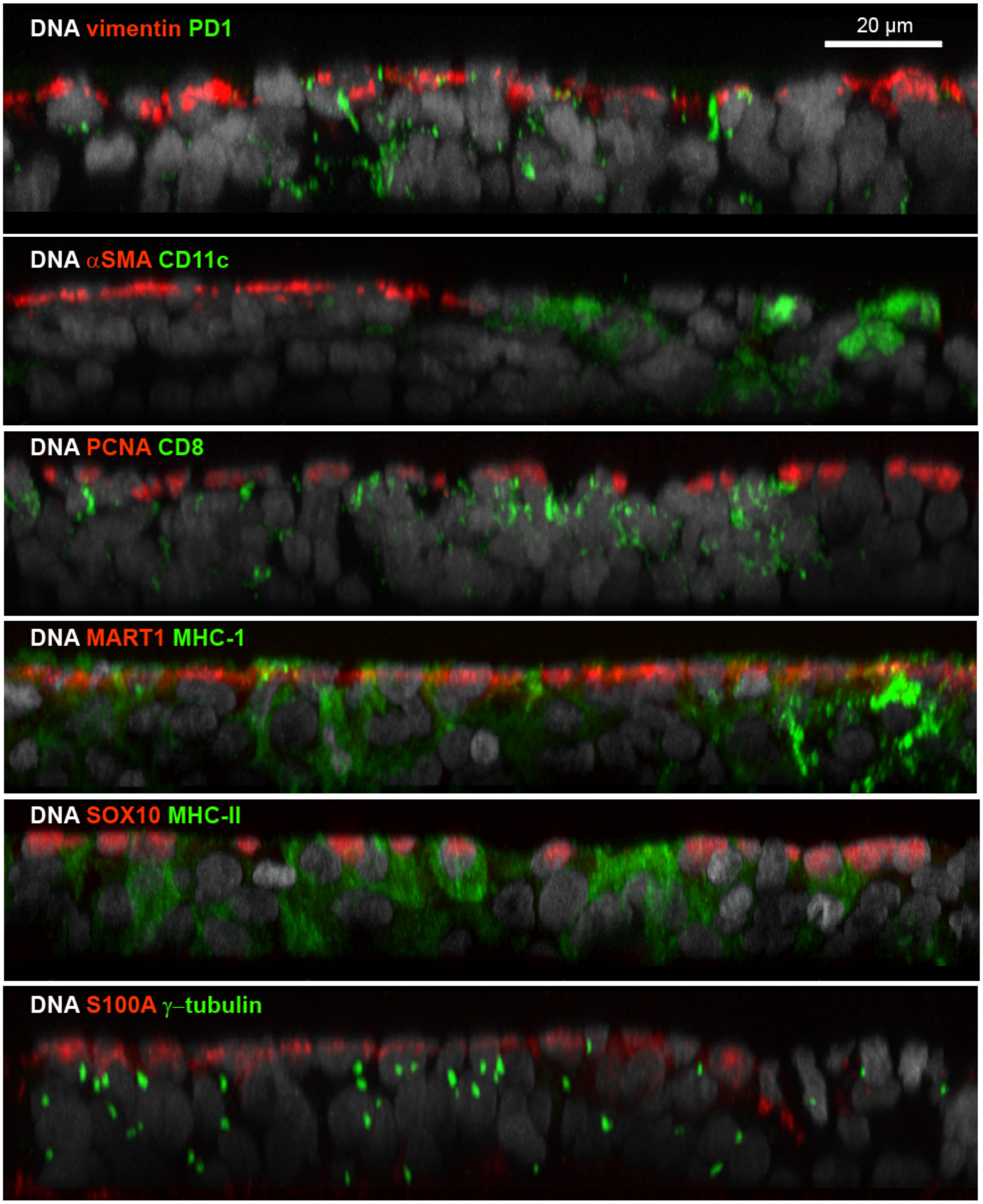


**Supplementary Figure 17. Orthogonal views comparing penetration of different antibodies in the same 35-micron thick melanoma tissue section (Dataset 2 – LSP13625).** Various antibodies (red) can exhibit poor antibody penetration whereas other antibodies (green) penetrate the full thickness of tissue. Tissue specimen was stained with antibodies as indicated across multiple rounds of CyCIF at 4 degrees C overnight on glass slide, washed in PBS, and mounted with 70% glycerol. Imaged with 40x/1.3NA oil objective lens on Zeiss LSM980 laser scanning confocal microscope sampled at 140nm (x,y) and 280nm (z). DNA stained with Hoechst shown in grey. Scale bar 20 µm.

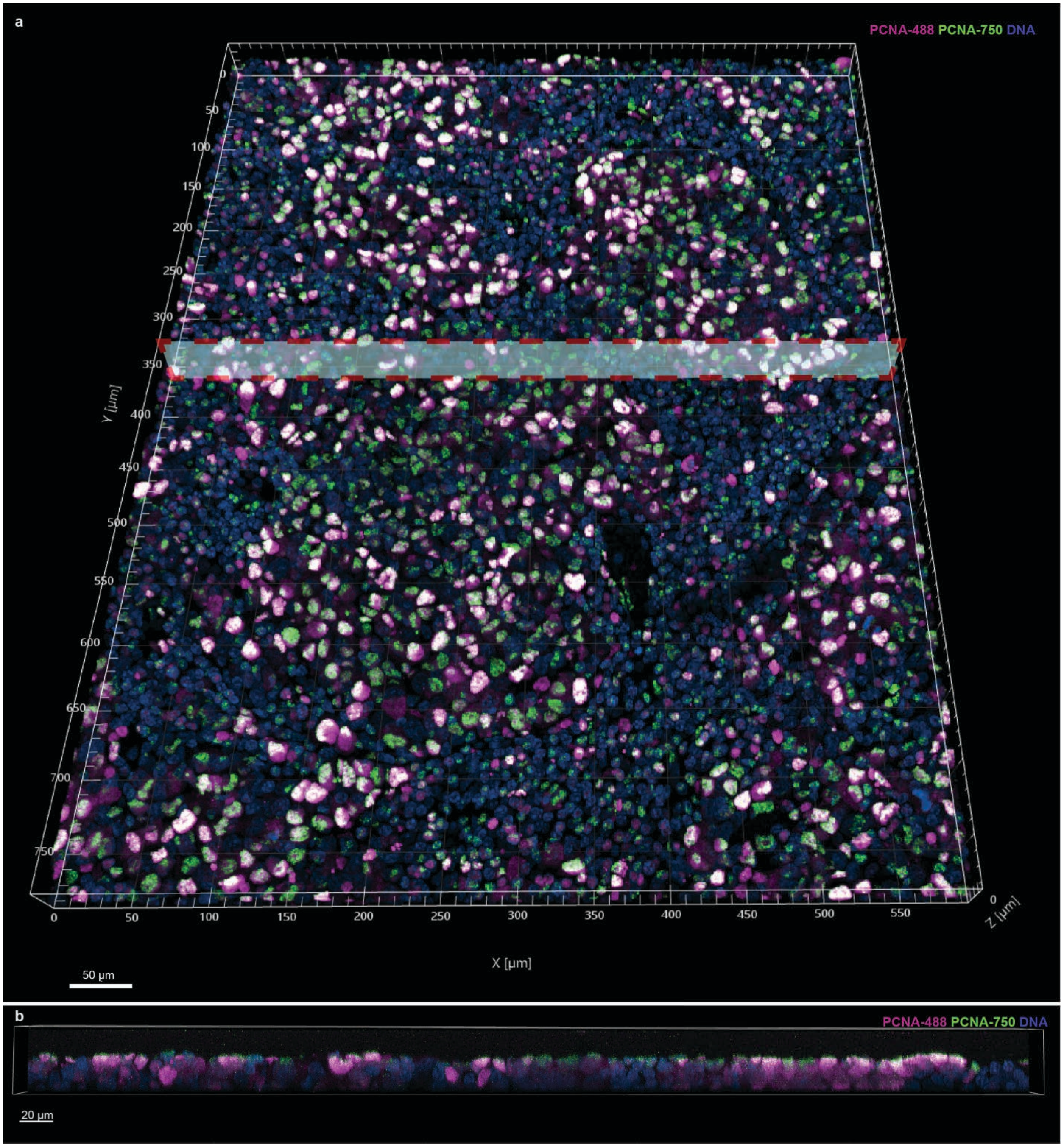


**Supplementary Figure 18. Antibody penetration comparison of PCNA conjugated with Alexafluor 488 and Alexafluor 750. a,** Volumetric rendering of 35 µm thick primary melanoma. Dashed red rectangle indicates location of orthogonal view in **b**. Scalebar 50 µm. **b** Results show that, in some antibodies such as PCNA, changing fluorophores can affect antibody penetration. For example, PCNA-488 (magenta) penetrates tissue more deeply than PCNA-750 (green). Tissue prepared and stained on glass slide at 4 degrees C overnight, washed in PBS and mounted with 70% glycerol. Imaged with 40x/1.3NA oil objective lens on Zeiss LSM980 laser scanning confocal microscope sampled at 140nm (x,y) and 280nm (z). Scalebar 20 µm.

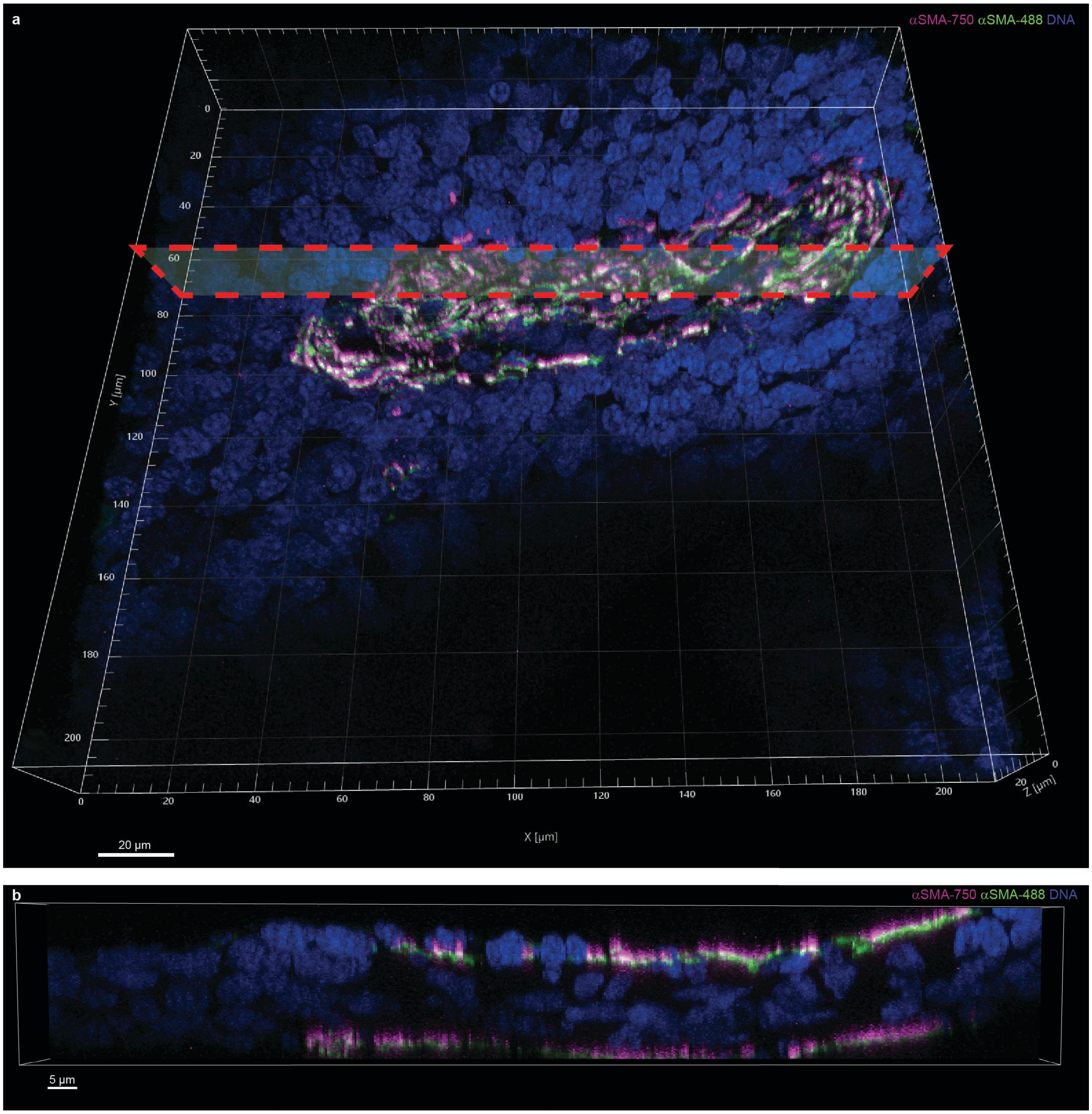


**Supplementary Figure 19. Antibody penetration comparison of αSMA conjugated with Alexafluor 488 (green) and Alexafluor 750 (magenta). a,** Volumetric rendering of primary melanoma with dashed red rectangle indicating location of orthogonal view in **b**. Scalebar 20 µm. **b,** Cross-sectional view of tissue. Both αSMA antibodies stain only the surface of the tissue regardless of fluorophore conjugated. DNA stained with Hoechst (blue). Tissue prepared and stained on glass slide at 4 degrees C overnight, washed in PBS and mounted with 70% glycerol. Imaged with 40x/1.3NA oil objective lens on Zeiss LSM980 laser scanning confocal microscope sampled at 414nm (x,y) and 300nm (z). Scalebar 5 µm.

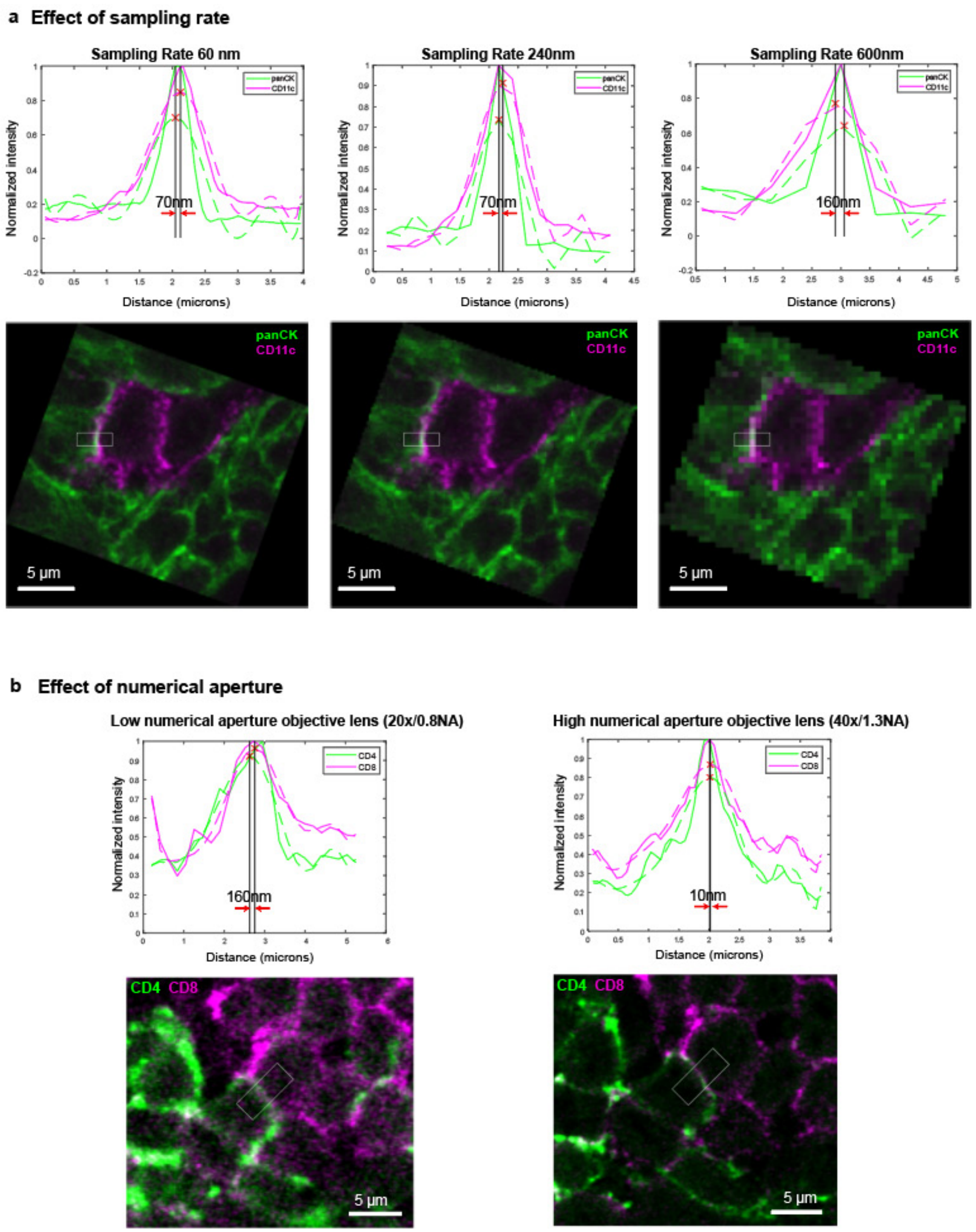


**Supplementary Figure 20. Effect of sampling rate and optical resolution on image quality and ability to identify cell to cell membrane contacts a,** dramatic reduction in sampling rate (increasing pixel size from 60nm to 600nm) showing marginal impact on the ability to detect peaks from line profiles. panCK – green, CD11c – magenta. Box indicates region that line profiles were extracted from. Tissue prepared and stained on glass slide at 4 degrees C overnight, washed in PBS and mounted with 70% glycerol. Imaged with 40x/1.3NA oil objective lens on Zeiss LSM980 laser scanning confocal microscope sampled at 140nm (x,y). Scalebar 5 µm. **b,** Line profiles and single optical z-planes of two interacting immune cells in a 5-micron thick colorectal cancer sample imaged at 20x/0.8NA (left) and 40x/1.3NA (right) objective lens. Using a lower magnification and numerical aperture (20x/0.8NA) results in lower photon sensitivity and a noisier image. Peak detection from line profiles is still possible. CD4 – green, CD8 – magenta.

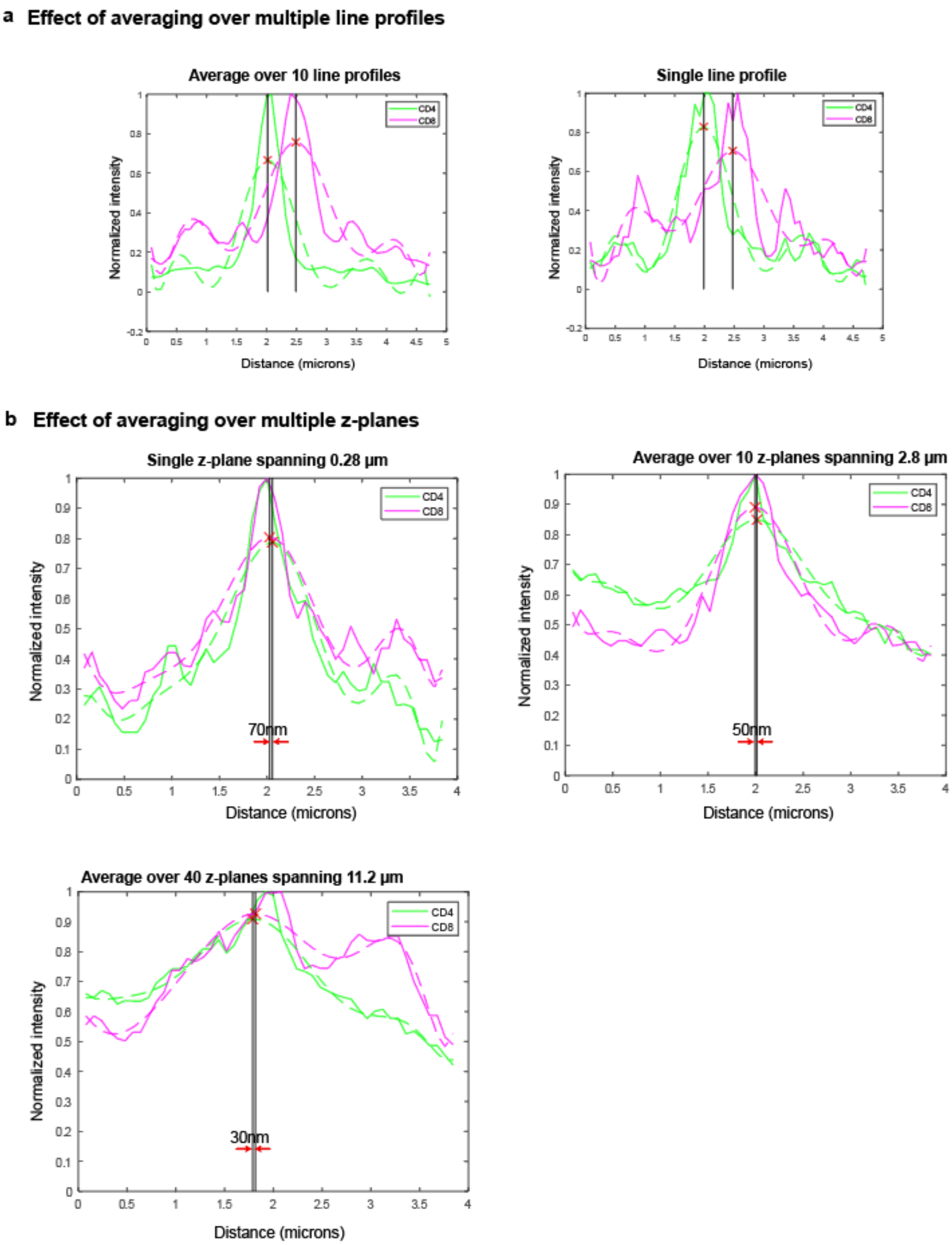


**Supplementary Figure 21. Effect of different line profile measurement methods on ability to identify cell to cell membrane contacts. a,** Comparison of line profiles taken from 10 parallel lines (left) vs 1 line (right) across the membrane of two neighbouring cells (**Supplementary Fig. 22b - right**). A single line profile is noisier than averaging multiple line profiles but peak detection is still possible. **b,** Single line profiles across 1, 10, and 40 z-planes in two cells (**Supplementary Fig. 20b - right)** are compared. Thicker z-planes may extend beyond the actual site of interaction thereby introduce noise and signal from other cells. CD4 – green, CD8 – magenta. Tissue prepared and stained on glass slide at 4 degrees C overnight, washed in PBS and mounted with 70% glycerol. Imaged on a Zeiss LSM980 laser scanning confocal microscope sampled at 140nm (x,y) and 280nm (z).

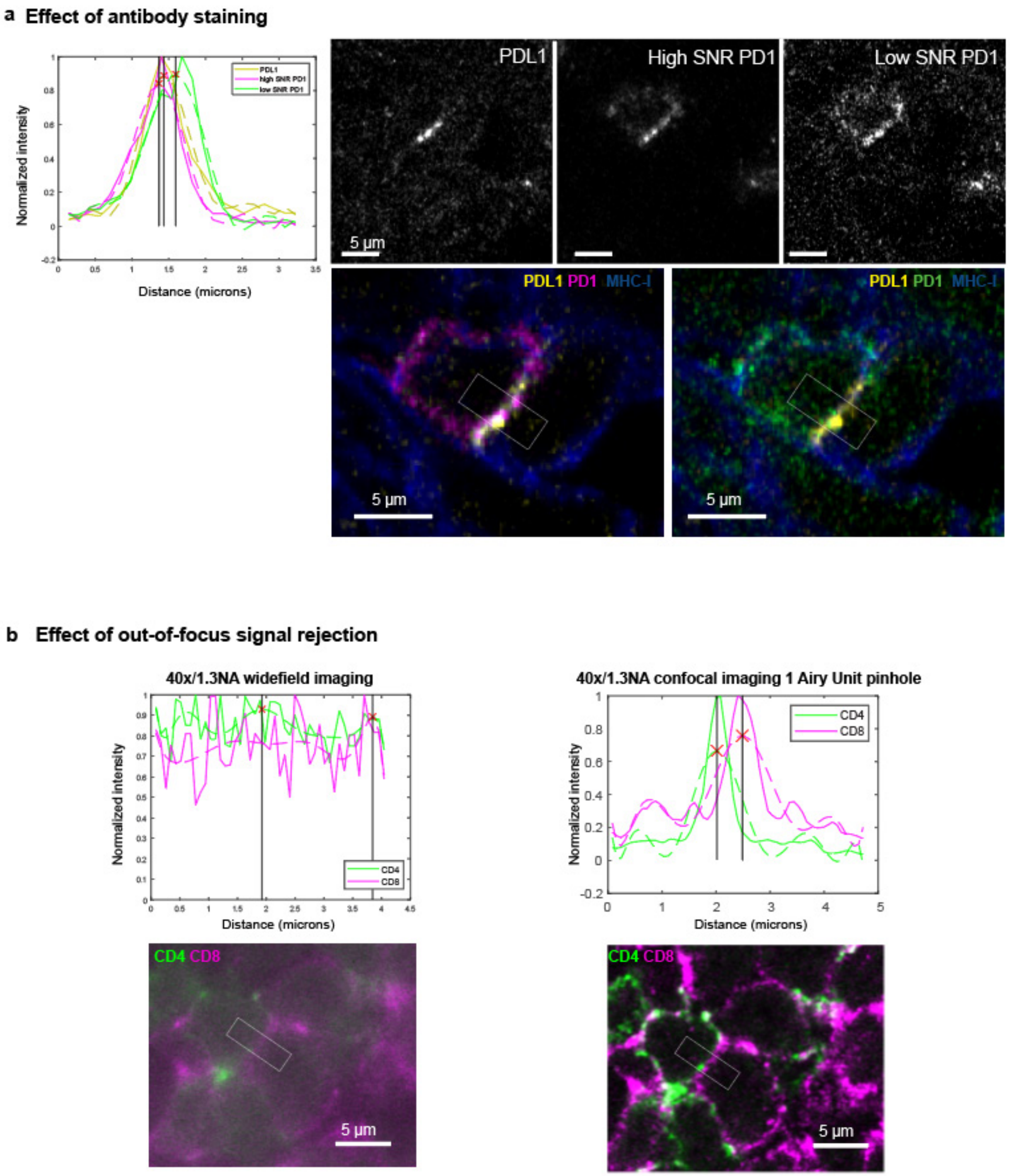


**Supplementary Figure 22. Effect of confocality and antibody signal ability to identify cell to cell membrane contacts. a**, Line profiles and single optical z-planes of two interacting cells stained with PDL1 (yellow), high SNR PD1 (magenta) and low SNR PD1 (green) in a 5-micron thick colorectal cancer tissue. MHC-1 shown in blue to demarcate cells membranes. The peak for low SNR PD1 has shifted due to noise. **b,** Line profiles and fluorescence images comparing widefield microscopy (left) and confocal imaging at 1 Airy Unit pinhole (right) on the exact same cells. Determining peaks in widefield is hindered due to severe out-of-focus signal. CD4 – green, CD8 – magenta. Box indicates region that line profiles were extracted from. Tissue prepared and stained on glass slide at 4 degrees C overnight, washed in PBS and mounted with 70% glycerol. Imaged on a Zeiss LSM980 laser scanning confocal microscope sampled at 140nm (x,y). Scalebar 5 µm.
