## Extended data figure for "Highly Multiplexed 3D Profiling of Cell States and Immune Niches in Human Tumours"

Extended Data Figure 1

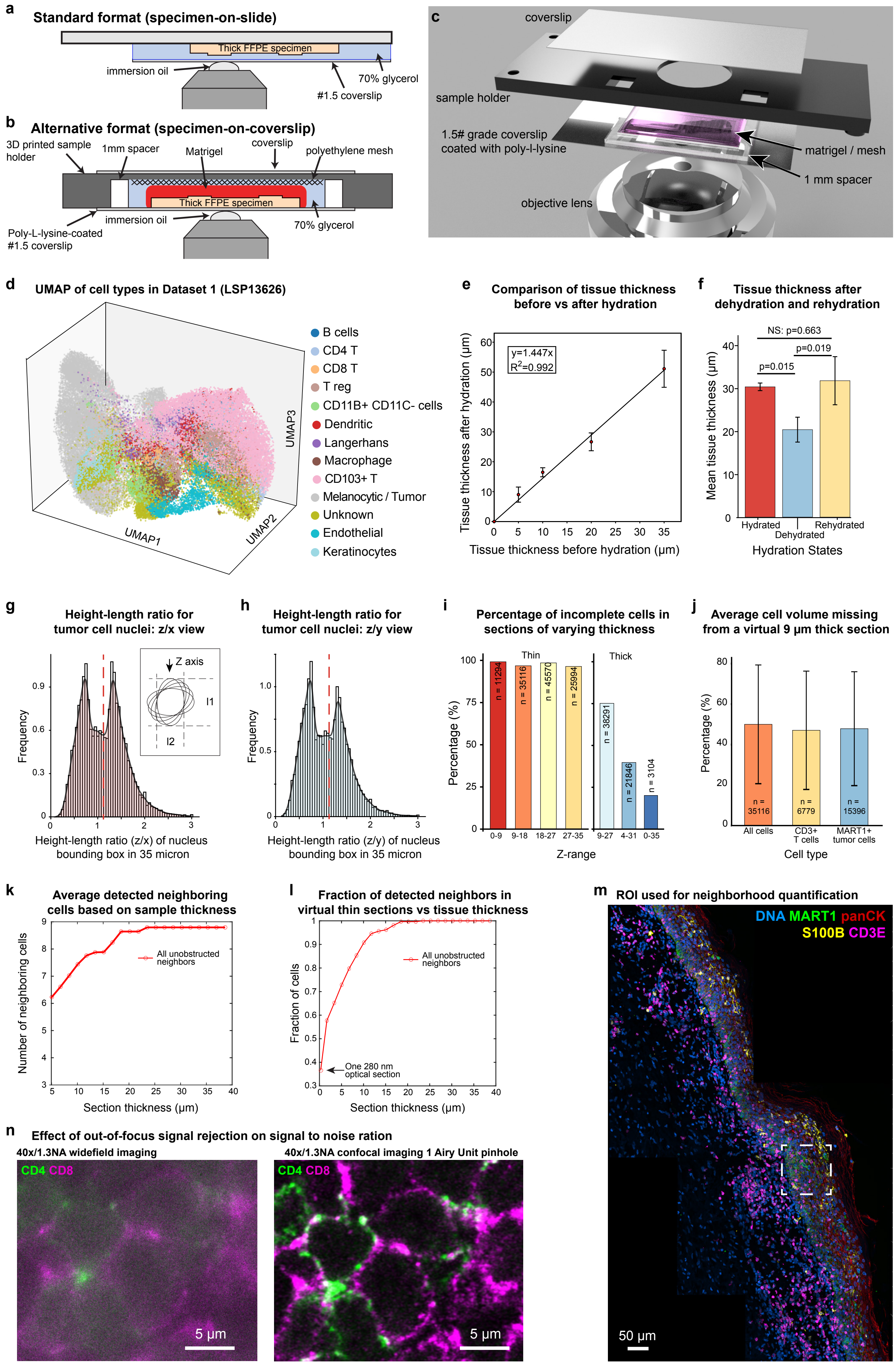

Extended Data Figure 2

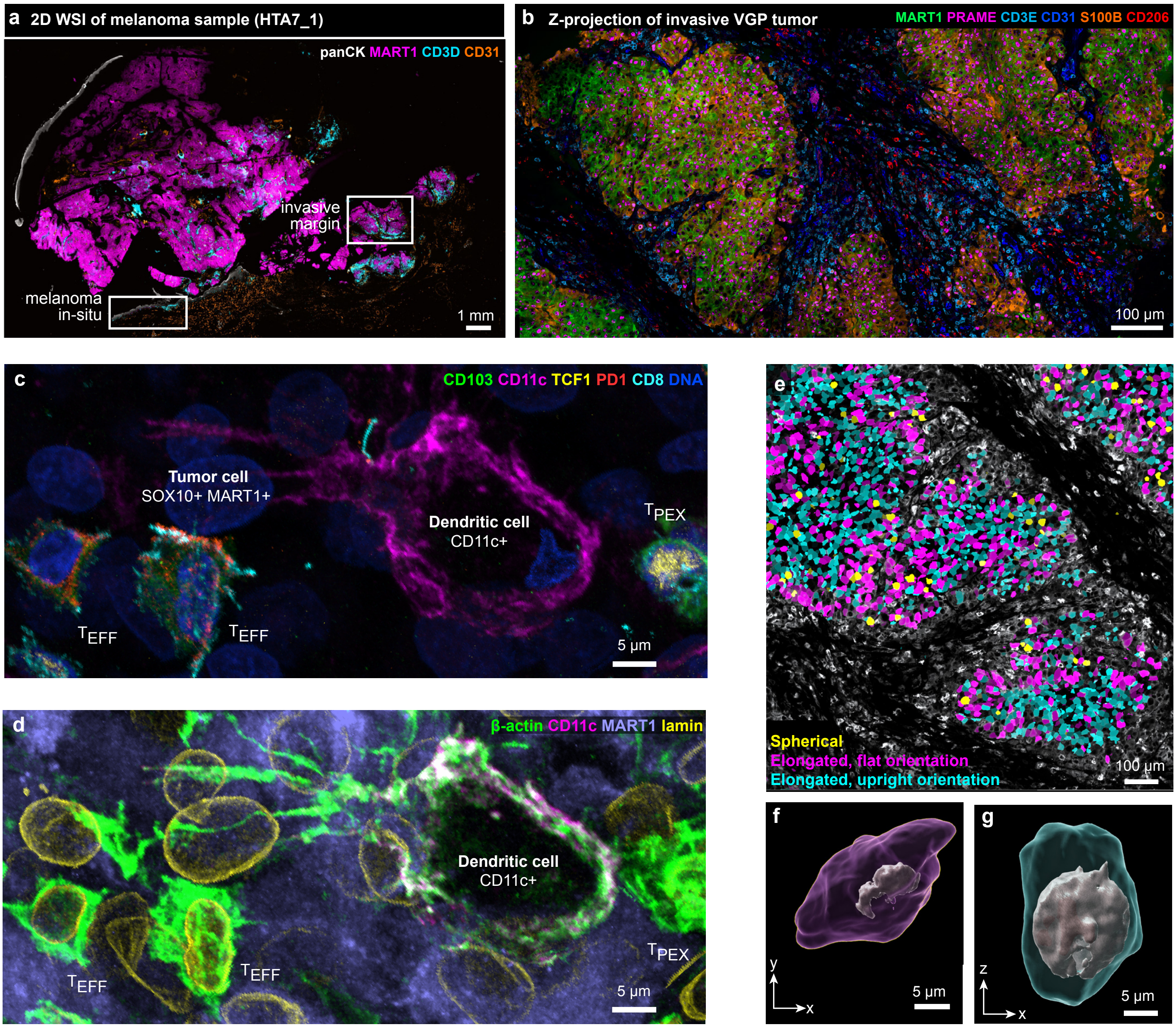

Extended Data Figure 3

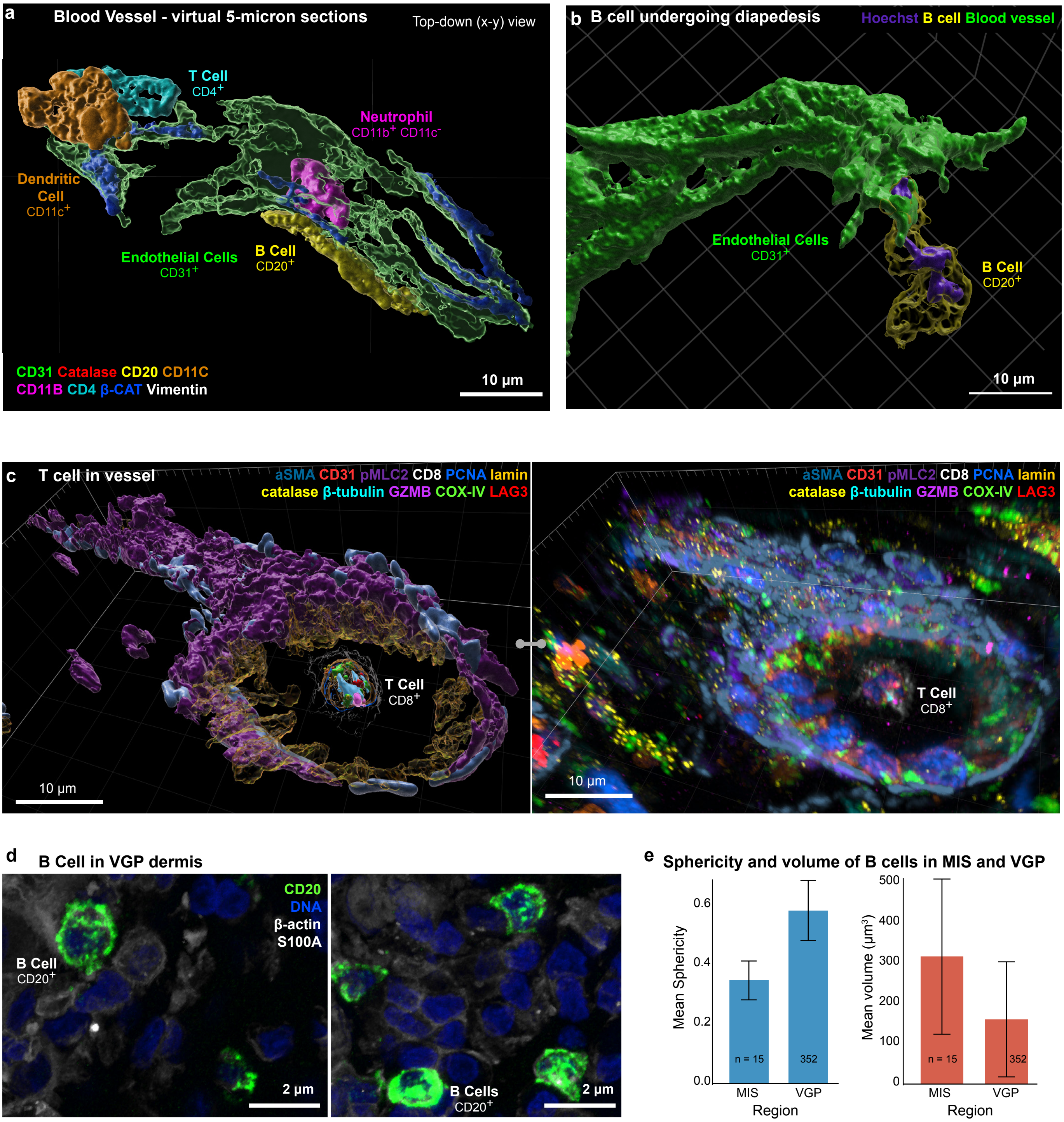

Extended Data Figure 4

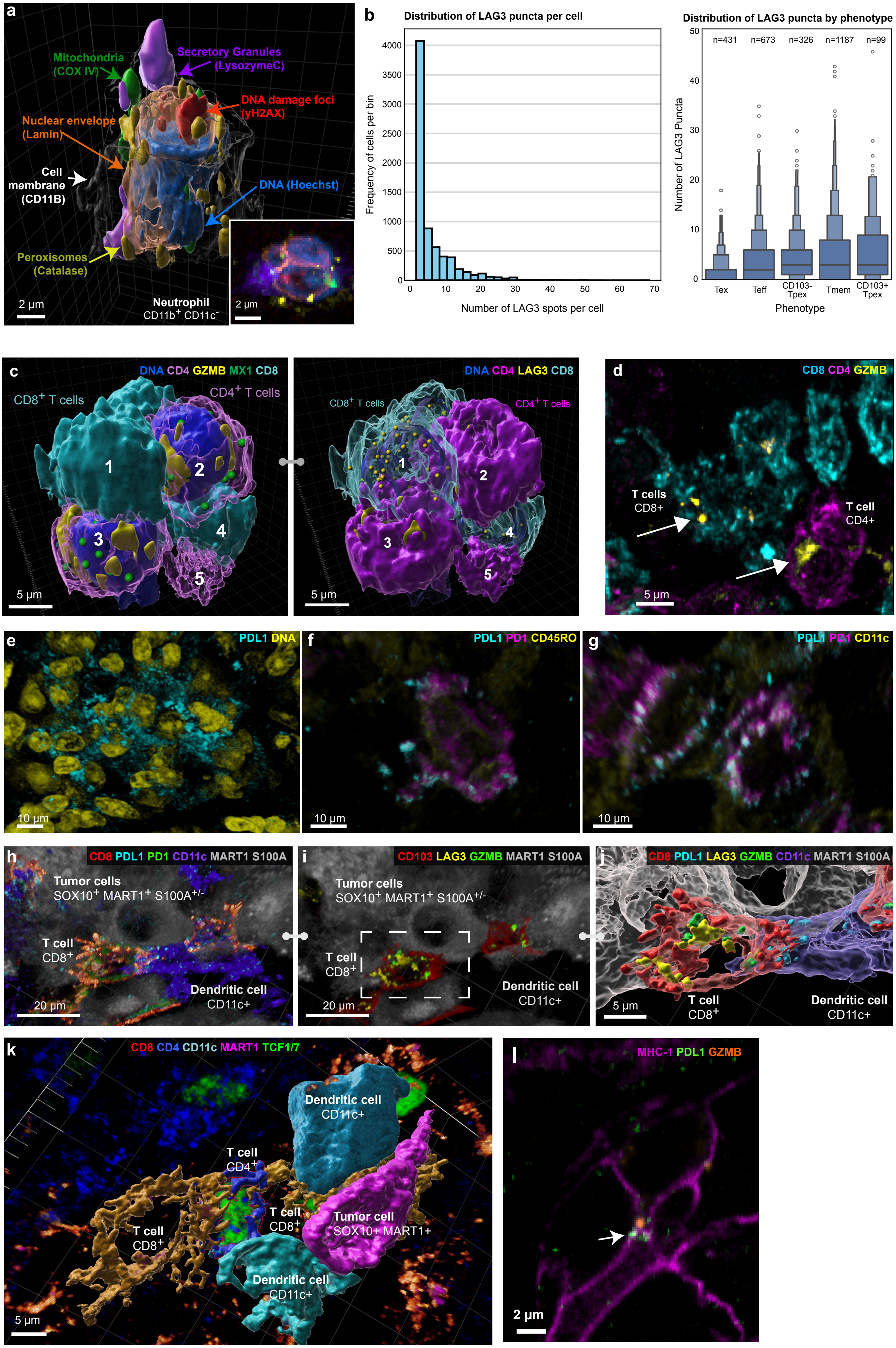

Extended Data Figure 5

**a**

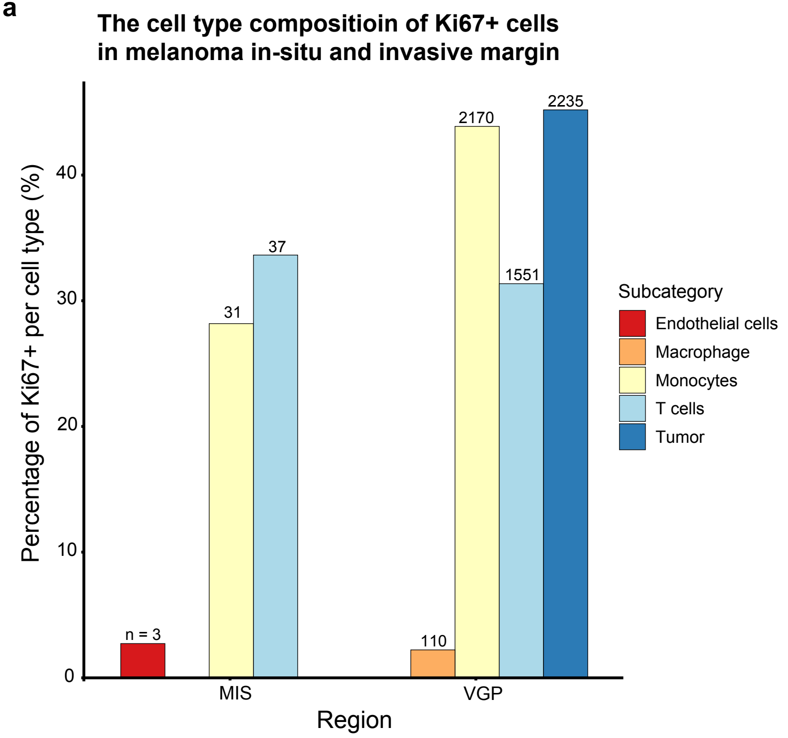


**b**

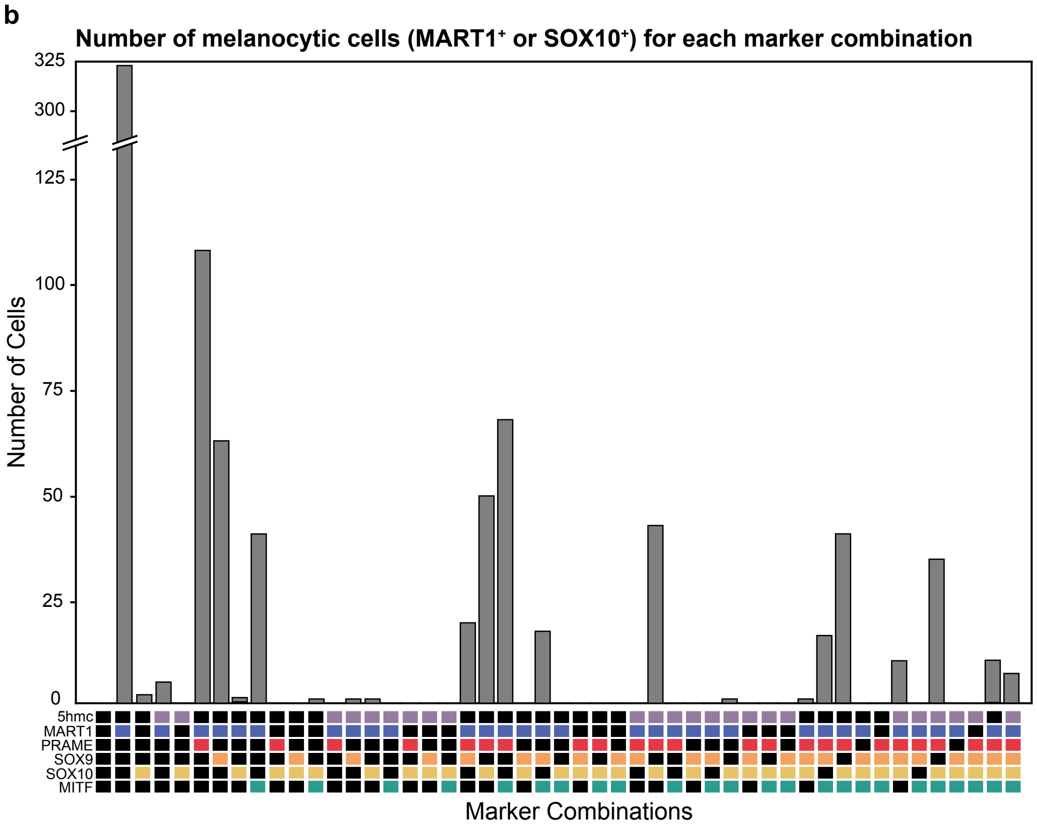

### Extended Data Figure 6

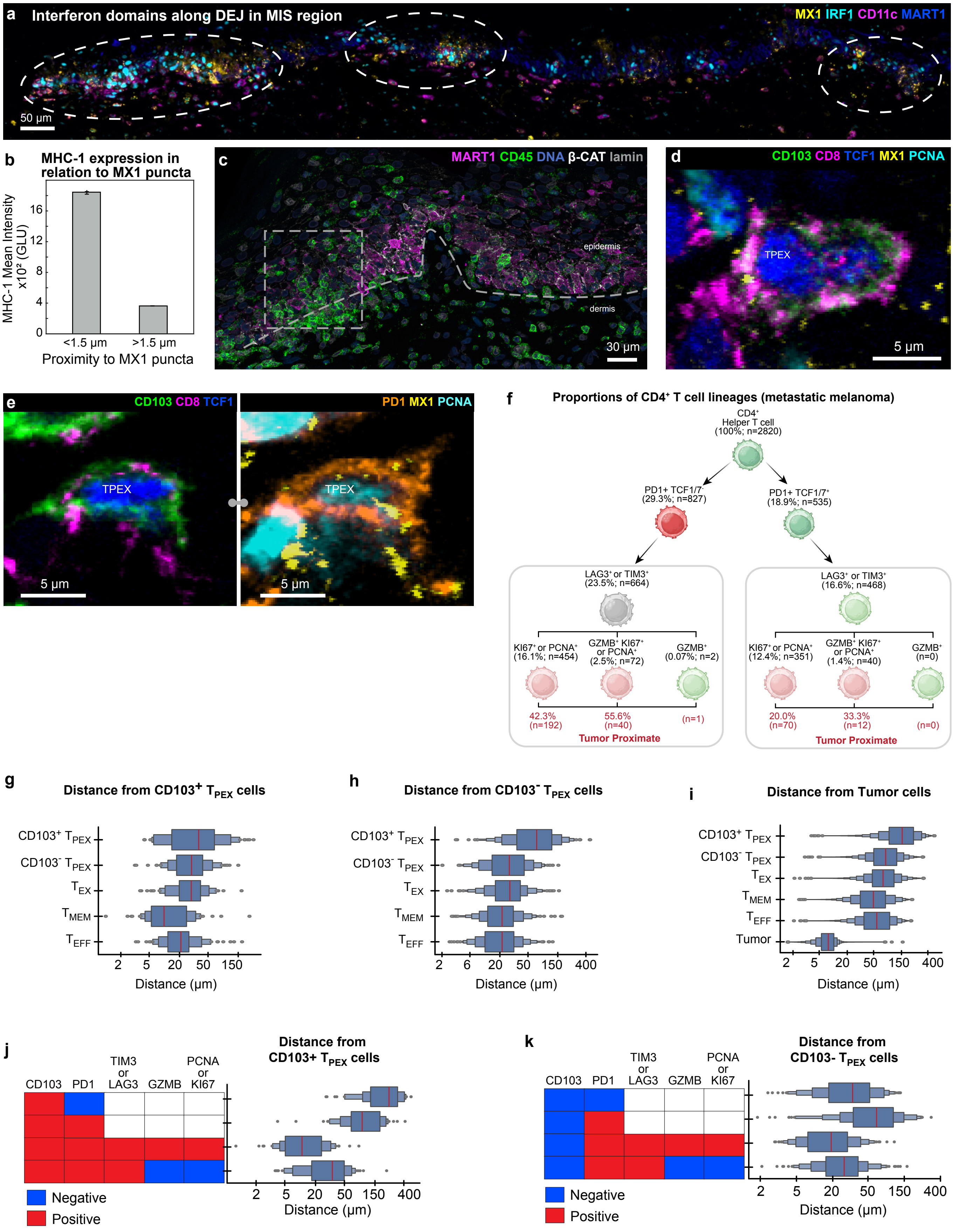

Extended Data Figure 7

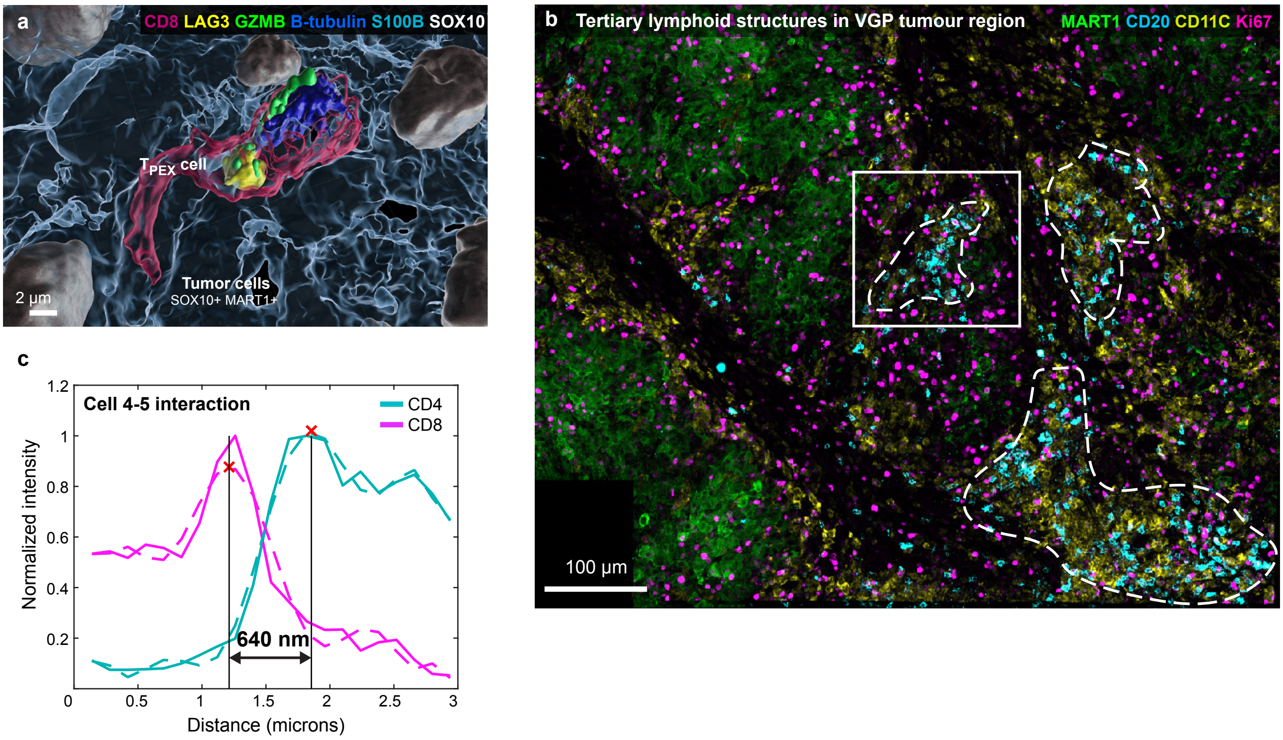

Extended Data Figure 8

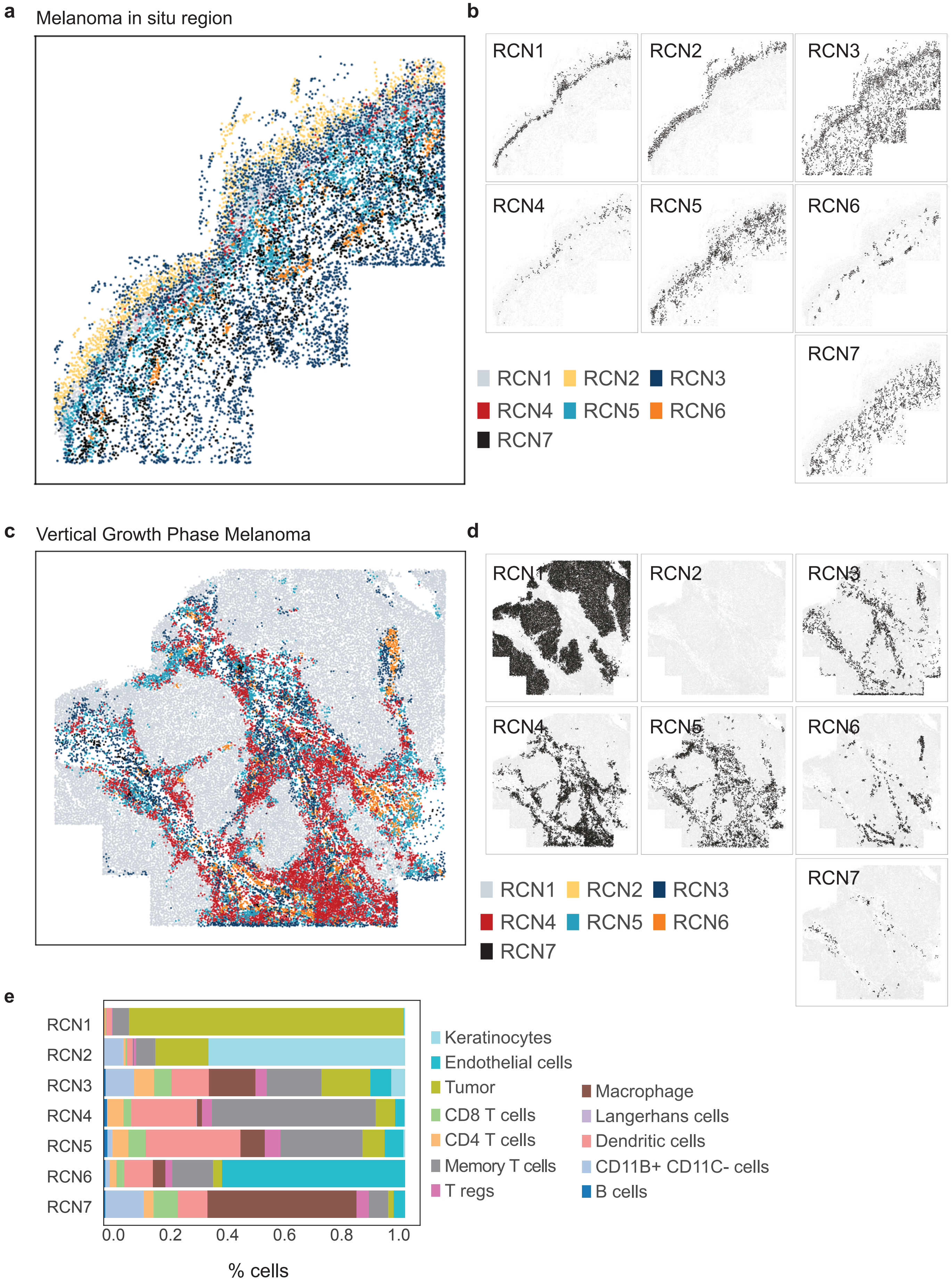
